# Selective inhibition of OSBP blocks retrograde trafficking by inducing partial Golgi degradation

**DOI:** 10.1101/2023.04.01.534865

**Authors:** Nianzhe He, Laura Depta, Cecilia Rossetti, Marko Cigler, Marine Michon, Oliver Rafn Dan, Joseph Hoock, Julien Barbier, Daniel Gillet, Alison Forrester, Georg E. Winter, Luca Laraia

**Affiliations:** Department of Chemistry, Technical University of Denmark, Kemitorvet 207, 2800 Kongens Lyngby, Denmark; CeMM Research Center for Molecular Medicine of the Austrian Academy of Sciences, 1090 Vienna, Austria; Université Paris-Saclay, CEA, INRAE, Département Médicaments et Technologies pour la Santé (DMTS), SIMoS, Gif-sur-Yvette 91191, France; Université de Namur ASBL, Rue de Bruxelles, 61-5000, Namur, Belgium

## Abstract

Sterol-binding proteins are important regulators of lipid homeostasis and membrane integrity; however, the discovery of selective small molecule modulators can be challenging due to structural similarities in the sterol binding domains. We report the discovery of highly potent and selective inhibitors of oxysterol binding protein (OSBP), which we term **oxybipins**. Sterol-containing chemical chimeras aimed at identifying new sterol binding proteins by targeted degradation, led to a significant reduction in Golgi-associated proteins. The degradation was found to occur at lysosomes, concomitant with changes in general protein glycosylation, indicating that the degradation of Golgi proteins was a downstream effect. By establishing a sterol transport protein biophysical assay panel, we discovered that the **oxybipins** potently inhibited OSBP, resulting in blockage of retrograde trafficking and attenuating Shiga toxin toxicity. As the **oxybipins** do not target any other sterol transporters tested, we advocate their use as chemical tools to study OSBP function and therapeutic relevance.

## Introduction

Lipid homeostasis can be regulated by global and local cholesterol levels, as well as by the levels of its metabolites, including oxysterols^1^. As such, discovering (oxy)sterol interacting proteins and tool compounds to modulate them may reveal new regulatory mechanisms and potential drug targets. Identification of sterol binding proteins has been carried out using affinity-based proteomic profiling with cholesterol-derived photoactivatable probes identifying more than two hundred putative cholesterol interacting proteins^2^, while a 20-(*S*)*-*hydroxycholesterol-derived probe identified the sigma-2 receptor as its primary target^3^. We recently developed a thermal proteome profiling-based approach to identify oxysterol target proteins^4^, however the large number of putative targets made validation efforts challenging.

We reasoned that target identification of (oxy)sterols could be achieved using an approach inspired by targeted protein degradation^5–8^. Generating proteolysis targeting chimeras (PROTACs) by linking sterols to E3 ligase ligands via different linkers, could lead to the proteasomal degradation of their targets, which could be retrospectively identified using mass spectrometry-based proteomics. While the requirements for ternary complex formation may reduce the number of potentially identifiable targets, we reasoned that this may actually be an advantage given the relatively promiscuous nature of sterols. Furthermore, a successful sterol-based PROTAC would immediately serve as a tool to study its identified target(s), particularly if it was selective in its degradation profile.

An important target class for sterols and sterol-based ligands are intracellular sterol transport proteins (STP). STPs bind and transport sterols, sharing high structural similarities in their sterol binding domains (SBD). They have distinct tissue distributions, intracellular localizations, and functions^9^. Additionally, they mediate organelle contacts and lipid metabolism^10^. They have been associated with several diseases including different cancer types, lipid storage disorders, and atherosclerosis^11^. The three main families of STPs are Oxysterol-binding Protein (OSBP)-Related Protein (ORP) family, the Steroidogenic Acute Regulatory Protein-related Lipid Transfer Domain (STARD) family and the Aster-domain containing family.

The ORP family contains 12 members, spliced into 16 different isoforms, all harboring the oxysterol-binding-protein-related ligand-binding domain (ORD), responsible for lipid binding^12^. OSBP, the prototypic member of this family, binds cholesterol, 25-hydroxycholesterol and PI4P competitively^13^. OSBP localizes to membrane contact sites formed between the ER and other organelles, including the *trans*-Golgi network (TGN), endosomes, and lysosomes. Additionally, OSBP plays critical roles in diseases. TTP-8307, a known enterovirus replication inhibitor, exerts its antiviral function by targeting OSBP^14^, and ORPphilins, a series of OSBP inhibitors, inhibit the growth of p21 deficient cancer cells^15^.

Herein we describe the preparation of sterol-based chemical chimeras and their degradation profiles. Significant degradation of Golgi-associated proteins was observed, with Golgi integral membrane protein 4 (GPP130 or GOLIM4) emerging as the primary target. Target engagement experiments and analysis of the degradation mechanism revealed that the compounds do not act as PROTACs, or directly target GPP130. Rather, they induce a more general lysosomal, rather than proteasomal, degradation of several Golgi proteins. By hypothesis elimination, the compounds were ultimately found to potently and selectively target OSBP, leading to inhibition of retrograde trafficking and a reduction in Shiga toxin toxicity.

## Results

### Design, synthesis and degradation profiles of cholesterol-bearing PROTACs

As a proof of concept, we initially focused our target identification efforts on cholesterol itself, given the wealth of accessible data on its putative targets^2^. We designed cholesterol (Chol)-bearing PROTACs with a sterol scaffold acting as the warhead for recognizing potential cholesterol-binding proteins without disrupting their critical binding interactions. The alkyl side chain was replaced with a carboxylic acid group for further linker attachment *via* amide coupling. A similar strategy has also been applied to the design of photo-labelling sterol probes^2^. Pomalidomide was chosen as the E3 ligand to recruit cereblon (CRBN), a member of Cullin Ring Ubiquitin Ligases (CRLs)^16^, while flexible linkers of various lengths, composition and hydrophobicity were used to connect the sterol core and the E3 ligand (**C1-C4**, Figure 1a, Extended Data Fig. 1). Introduction of the amide-bond increases the polarity of the whole molecule and may further affect its cell permeability; as such, to exclude proteins affected by the sterol scaffold and the additional amide bond in absence of the linker and E3 ligase recruiter, the *N-*methylated analogue **C5** was designed as a negative control.

**Fig. 1.**
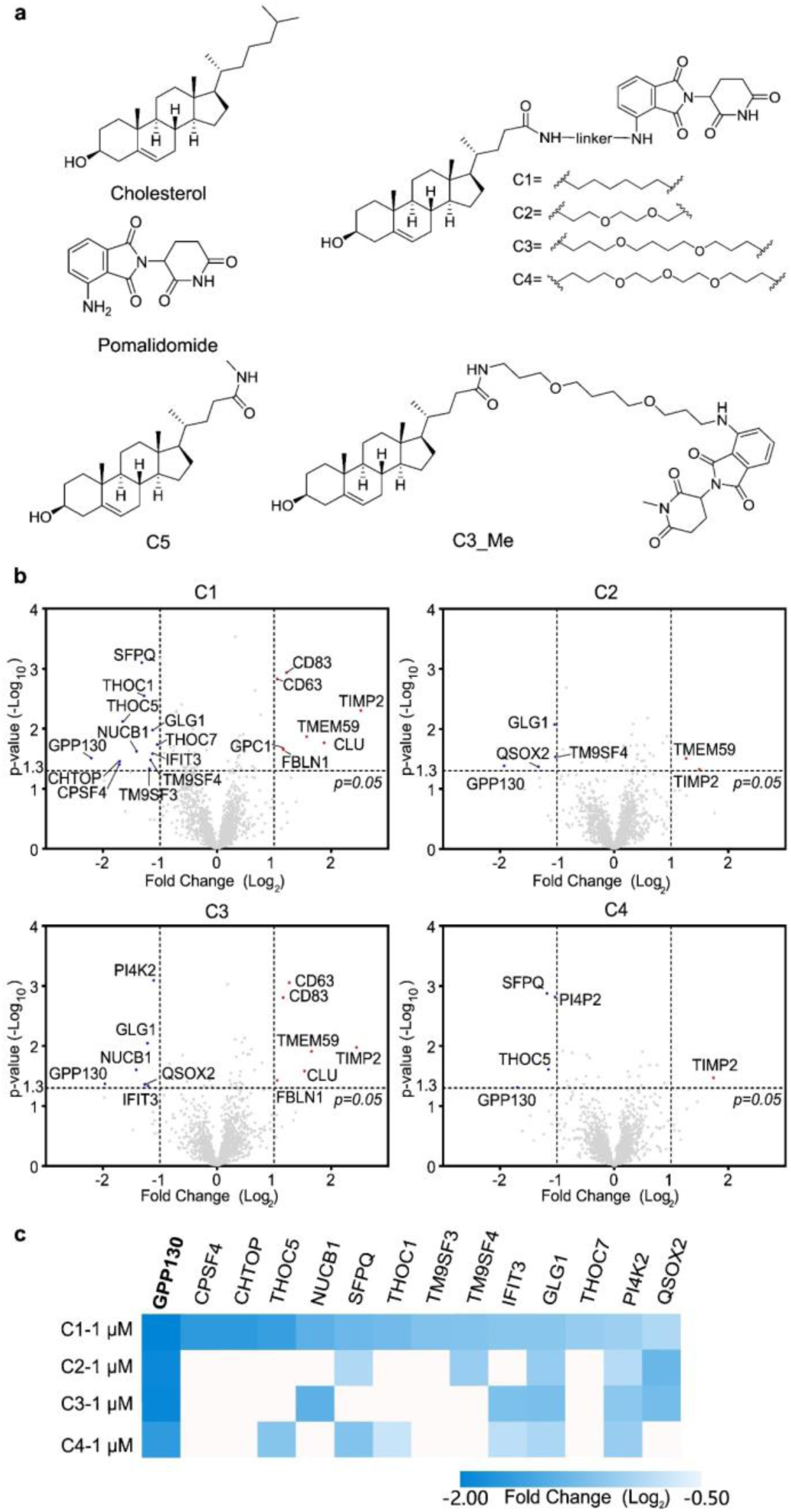
Structures and degradation profiles of Chol-based PROTACs. (a) Structures of cholesterol, pomalidomide, Chol-bearing PROTACs, **C1**-**C4**, negative control **C5** and *N*-methylated pomalidomide-bearing PROTAC, **C3_Me**. (b) Degradation profiles of **C1**-**C4** at 1 μM in HeLa cells treated for 18 hours. The down-regulated proteins (blue dots) and up-regulated proteins (red dots) were plotted as log2 fold change (compound-treated /DMSO-treated) versus -log10 (p-value). P-values were calculated from the data of three technical replicates. The dotted horizontal line marks the significance threshold of p = 0.05 and dotted vertical lines represent the threshold of an absolute log2 fold change of 1. (c) Heatmaps of degraded proteins under the treatment of 1 μM **C1**-**C4**. Please see Extended Dataset 2 for complete proteomics data.

To investigate the proteins affected by **C1**-**C5** in an unbiased way, we used tandem mass tag (TMT)-based proteomics. HeLa cells were treated with 100 nM and 1 μM **C1-C4** and 1 µM **C5** for 18 hours to guarantee a pronounced and sustained degradation of the potential targets. Cells were lysed using PBS containing NP-40 as this detergent has been suggested to aid the solubilization of membrane proteins, where most known sterol targets are located^17^. After protein digestion, the peptides were labelled with the corresponding TMT labels to enable multiplexed MS analysis.

In total, we identified 2454-4064 proteins by expression proteomics (Extended Dataset 1). With a threshold of fold change > 2 or < 0.5, and p < 0.05, no protein was significantly affected by the treatment of 100 nM **C1-C4** or 1 μM **C5** (Extended Data Fig. 2). However, the abundance of 21 proteins was significantly altered upon treatment with 1 μM **C1**-**C4** (Figure 1b), with 14 proteins being degraded and 7 being up-regulated. Among these, five proteins (GPP130, TM9SF3, TM9SF4, QSOX2 and CD63) have previously been identified as cholesterol interacting proteins by photo-affinity cholesterol probes^2^, which provides a potential validation of this method for target identification. Theoretically, PROTACs should reduce protein levels of their primary targets. Therefore, up-regulated proteins were considered as the downstream effects of the degradation of specific proteins and were not further analyzed.

**Fig. 2.**
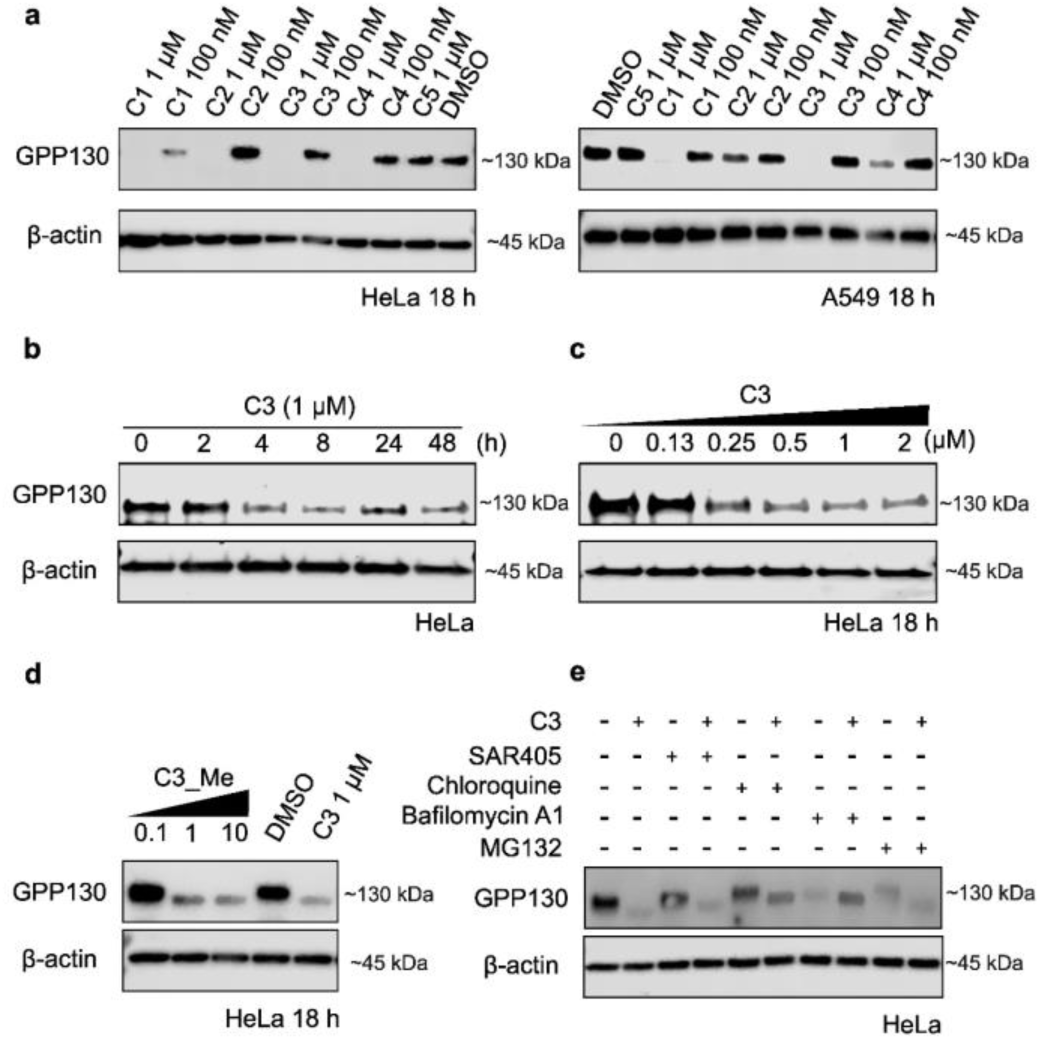
Validation of GPP130 degradation by **C1-C4**.(a) Changes in GPP130 levels in response to **C1-C4** in HeLa and A549 cell lines (18 h, n = 2). (b) Time dependency of GPP130 degradation in HeLa cells by **C3** (n = 4). (c) Dose-dependent degradation of GPP130 in HeLa cells by **C3** (18 h, n = 4). (d) Degradation of GPP130 in HeLa cells by **C3_Me** (18 h, n = 2). (e) Requirement of a functional proteasome or lysosome for the degradation of GPP130 by **C3**. HeLa cells were treated with MG132 (10 μM), chloroquine (20 μM), bafilomycin A1 (100 nM) and SAR405 (100 nM) in the absence/presence of **C3** (1 μM) for 18 hours (n = 3). Please see supplementary Figure 1 for uncropped blots.

**C1**-**C4** exhibited distinct degradation profiles at 1 μM (Figure 1c). **C1**, the only PROTAC containing an alkyl linker, degraded the most proteins (12) while PEG linker containing PROTACs, **C2**-**C4**, led to the decrease in abundance of ≤ 6 proteins. However, the effects of **C1**-**C4** also shared some similarities: all led to the reduction of GLG1, PI4K2, and GPP130. For further target validation, we chose GPP130, the protein that was most significantly degraded by all Chol-PROTACs and a known target of cholesterol, for further study. GPP130 is a Golgi-resident glycoprotein involved in retrograde trafficking^18^. Loss of GPP130 has been reported to protect from Shiga toxin toxicity, making its degradation by small molecules potentially therapeutically relevant^19^.

### The degradation of GPP130 is mediated by the lysosome rather than the proteasome

Consistent with the proteomics data, western blotting showed that significant degradation of GPP130 (> 90%) could be observed in HeLa cells treated with 1 μM **C1-C4** or 100 nM **C1**, while no obvious degradation was observed with 100 nM **C2-C4**. In contrast, no significant depletion of GPP130 was observed in cells treated with 1 μM **C5**, which indicates that the sterol scaffold alone is insufficient to cause GPP130 degradation (Figure 2a). To further assess the activity of **C1**-**C4** across different cell lines, the human lung cancer cell line, A459, was chosen (Figure 2a). Among them, **C1** and **C3** still exhibited effective GPP130 degradation abilities; causing a decrease of over 90 % of GPP130 at 1 μM. The degradation abilities of **C2** and **C4** were limited in A459 cell lines, with less than 50 % of GPP130 degraded. Additionally, **C3** exhibited the best degradation ability and selectivity towards GPP130. Therefore, **C3** was selected for further validation experiments.

**C3** (1 µM) led to a time dependent degradation of GPP130 over a period of 48 hours, with an initial degradation observed at four hours, and a maximal degradation at around 8 hours, which was sustained up to 48 hours (Figure 2b). **C3** also exhibited a concentration dependent degradation of GPP130 (Figure 2c) with maximal degradation achieved at 2 μM. Additionally, the degradation of GPP130 was independent of the choice of E3 ligands as **VH032**, a recruiter or the E3 ligase von Hippel Lindau (VHL)^20, 21^, also led to degradation of GPP130 (Extended Data Fig. 3, 4).

**Fig. 3.**
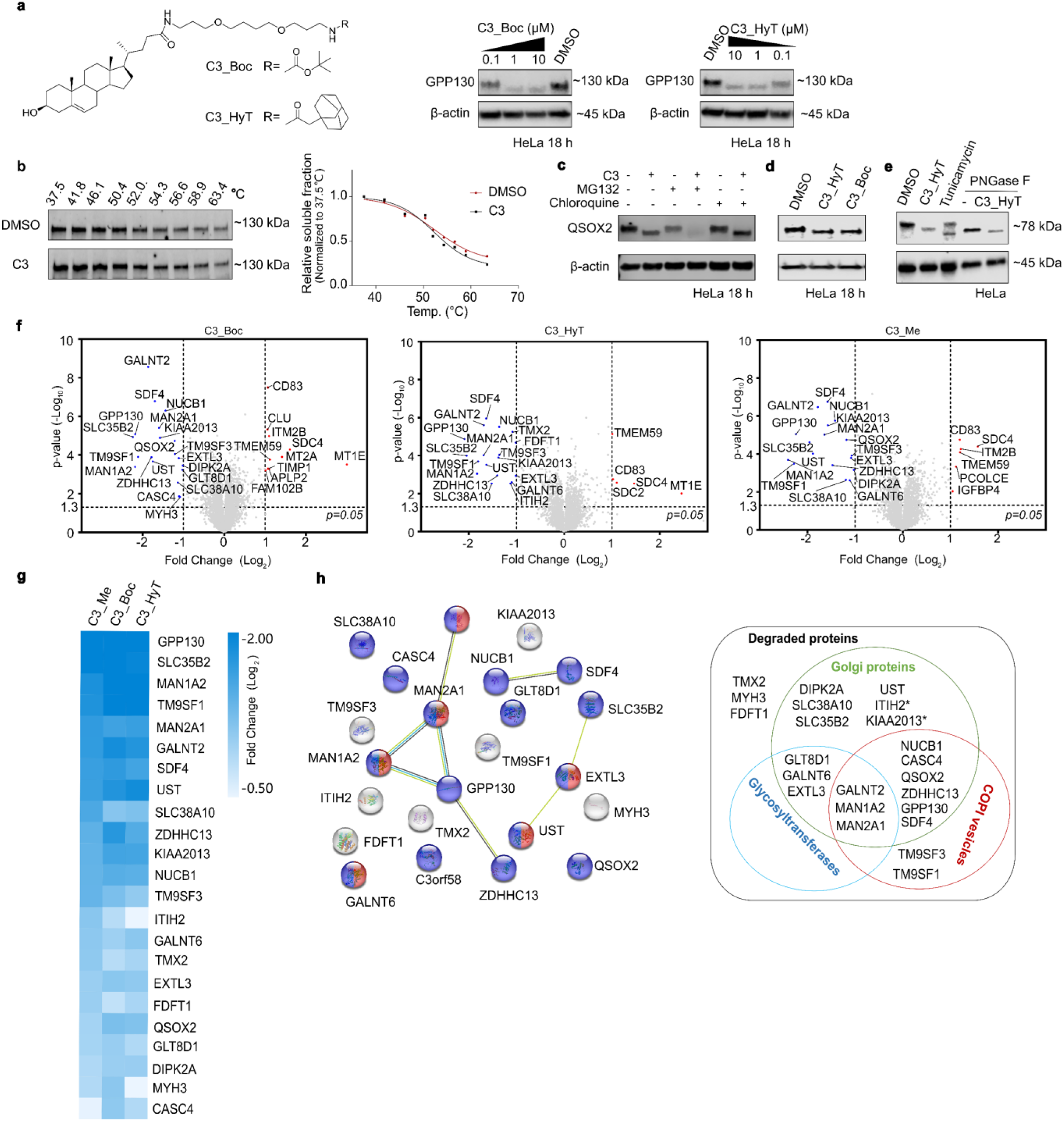
Derivatives of **C3** significantly degrade Golgi proteins. (a) Degradation of GPP130 by **C3_Boc** and **C3_HyT** (n = 2). (b) Thermal stability of GPP130 in HeLa cell lysate in response to **C3** as assessed by a cellular thermal shift assay (n = 2). (c) Mechanism of QSOX2 degradation by **C3**. HeLa cells were treated with **MG132** (10 μM), chloroquine (20 μM) in the absence/presence of **C3** (1 μM) for 18 hours (n = 2). (d) Both **C3_Boc** and **C3_HyT** can degrade QSOX2 at 1 μM in HeLa cells after 18 h (n = 2). (e) *N*-glycosylation of QSOX2 is removed by tunicamycin (5 μg/mL) for 18 hours, PNGase F and **C3_HyT** (n = 2). (f) Degradation profiles of **C3_Me**, **C3_Boc** and **C3_HyT** at 1 μM for 18 hours in HeLa cells. The down-regulated proteins (blue dots) and up-regulated proteins (red dots) were plotted as log_2_ fold change (compound-treated /DMSO-treated) versus -log_10_ (p-value). P-values were calculated from the data of three technical replicates. The dotted horizontal line marks the significance threshold of p = 0.05 and dotted vertical lines represent the threshold of an absolute log_2_ fold change of 1. (g) Heatmaps of degraded proteins under the treatment of 1 μM **C3_Me**, **C3_Boc** and **C3_HyT**. (h) The degraded proteins can be classified into three groups. Golgi proteins (Blue dots and green circle GO: 0005794; FDR 8.76e-10). Golgi proteins not enriched by STRING functional analysis are labelled with a star*; Glycosyltransferases (Blue circle); proteins involved in glycoprotein biosynthetic process (Red dots, GO:0009101; FDR 0.0146); COPI vesicles (Red circle). Please see supplementary Figure 1 for uncropped blots and Extended Dataset 2 for complete proteomics data.

**Fig. 4.**
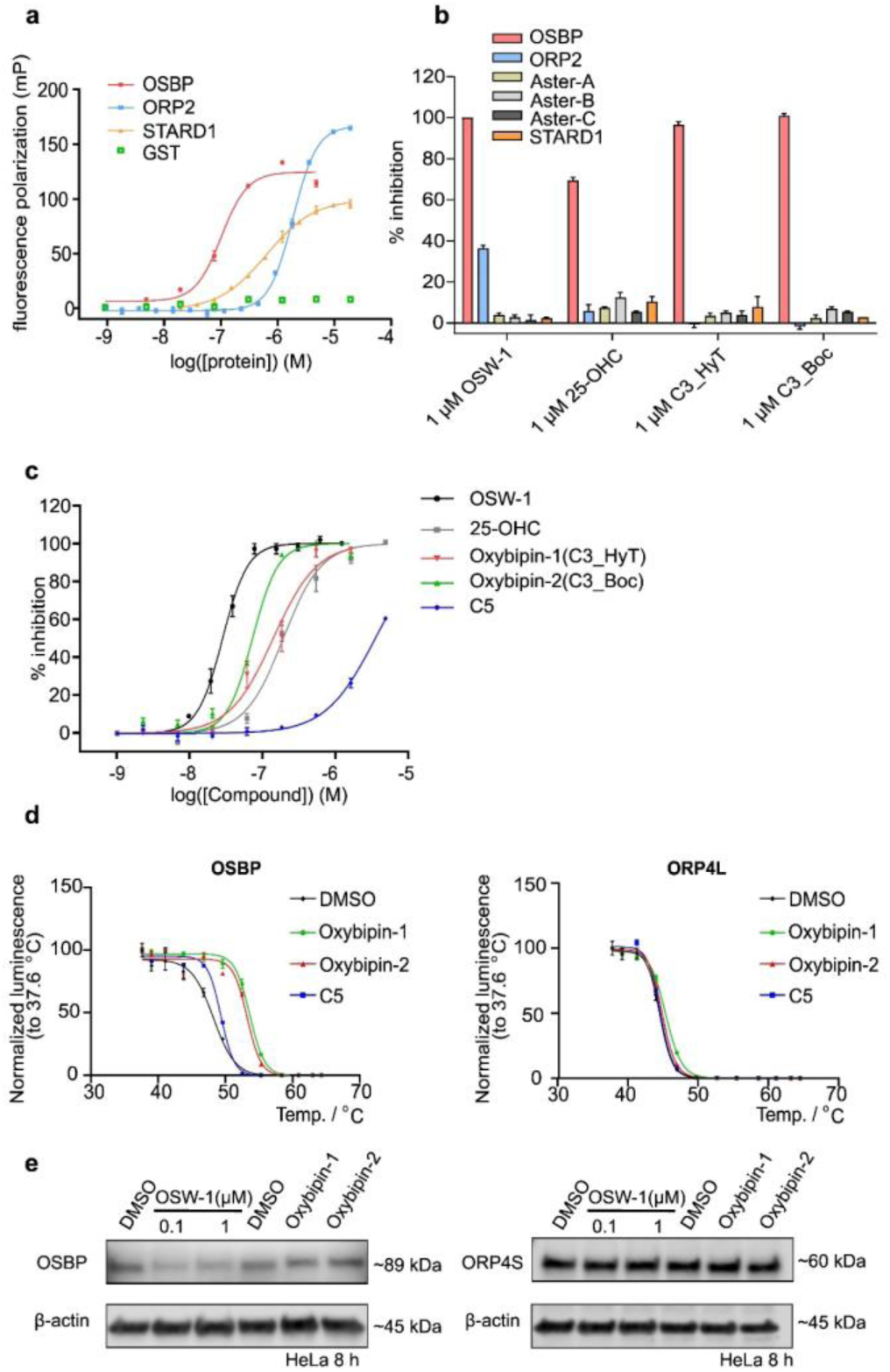
**Oxybipin-1** and **-2** selectively target OSBP. (a) Binding curve of the FP titration of 22-NBD-cholesterol with STPs and GST. (b) Screening of **C3_HyT (oxybipin-1), C3_Boc (oxybipin-2), OSW-1** and **25-OHC** in the STP FP selectivity panel at 1 μM. (c) **Oxybipin-1** and **-2** bind to the ORD domain of OSBP. (d) Effect on the stability of OSBP and ORP by **oxybipin-1** and **-2** in intact KBM7 cells expressing HiBit tagged fusion proteins. (e) Effect on cellular OSBP and ORP4S levels by 1 μM **oxybipin-1** and **-2** as well as **OSW-1** in HeLa cells after 8 hours. For all experiments data is mean ± sem, n = 3. Please see supplementary Figure 1 for uncropped blots.

To explore whether CRBN is needed for GPP130 degradation, **C3_Me** was synthesized by replacing pomalidomide with *N*-methylated pomalidomide, which is known to abrogate binding to CRBN^22^ (Figure 1c). Surprisingly, **C3_Me** still led to GPP130 degradation in a concentration dependent manner, similarly to **C3**, suggesting that CRBN is not involved in GPP130 degradation (Figure 2d). Due to this observation, we investigated whether the degradation of GPP130 was performed by the ubiquitin-proteasome system (UPS) (Figure 2e). HeLa cells were treated with the proteasome inhibitor **MG132** (10 μM), in conjunction with **C3** (1 μM) for 18 hours; however, the GPP130 levels were even lower than when treated with **C3** alone, indicating that GPP130 degradation by **C3** is not mediated by the UPS.

Interestingly, the treatment of **MG132** alone led to a pronounced decrease in GPP130 levels. Based on the fact that the long-term (>18 hours) proteasome inhibition stimulates lysosome degradation^23^, we investigated whether **C3** induced lysosomal degradation of GPP130. To this end, cells were treated with the autophagosome-lysosome fusion inhibitor chloroquine, or with the v-ATPase inhibitor bafilomycin A1. The level of GPP130 was not affected by the treatment of chloroquine (20 μM) itself. Co-treatment with chloroquine (20 μM) and **C3** (1 μM) rescued the degradation of GPP130, strongly suggesting that it is mediated by the lysosome. Interestingly, the co-treatment with bafilomycin A1 (100 nM) and **C3** (1 μM) did not lead to GPP130 degradation, while bafilomycin A1 (100 nM) alone did. GPP130 reversibly redistributes to the endosome upon pH disruption caused by treatments such as chloroquine and bafilomycin A1^24–26^. However, the mechanism by which **C3** rescues the degradation of GPP130 induced by bafilomycin A1 remains unclear. We speculate that prolonged inhibition of lysosomal activity may re-direct protein degradation to the proteasome when the lysosome is strongly de-acidified, while not if the fusion of lysosomes and autophagosomes is impaired^27^, however this remains to be investigated.

Additionally, to investigate whether autophagy is involved in GPP130 degradation, HeLa cells were treated with SAR405 (100 nM), an inhibitor of PIK3C3/VPS34, which leads to inhibition of autophagosome biogenesis^28^. SAR405 did not block the degradation of GPP130 induced by **C3** which indicates that early autophagy is not involved (Figure 2e). Overall, **C3** does not function as a PROTAC in its degradation of GPP130; degradation is mediated by the lysosome and not the UPS or autophagy.

### Cholesterol coupled with a linker is enough to trigger GPP130 degradation

As the degradation of GPP130 by **C3** does not rely on CRBN or the proteasome, we assumed that the pomalidomide part may not be required for GPP130 degradation. Therefore, we attempted to optimize **C3** into smaller GPP130 degraders. The minimization of **C3** was carried out using two approaches. First, the fact that **C3** successfully degrades GPP130 but **C5** does not indicates that the linker still plays a role in GPP130 degradation. Therefore, the pomalidomide part of **C3** was removed (Fig. 3a) but the linker was kept, to obtain **C3_Boc**. Second, **C3_HyT** was designed by replacing pomalidomide with adamantane, which is a representative moiety used in hydrophobic tagging (HyT) strategies^29^, to study whether pomalidomide was recognized as a hydrophobic moiety to trigger GPP130 degradation. Both **C3_Boc** and **C3_HyT** induced GPP130 degradation in a dose-dependent manner, which confirmed that a sterol core connected to a PEG linker is sufficient to trigger GPP130 degradation (Figure 3a, Extended Data Fig. 5). Among all GPP130 degraders, **C3_HyT** exhibited the most potent GPP130 degradation ability; >20% of GPP130 was degraded at 100 nM while no noticeable GPP130 degradation effect was observed for other compounds at this concentration.

**Fig. 5.**
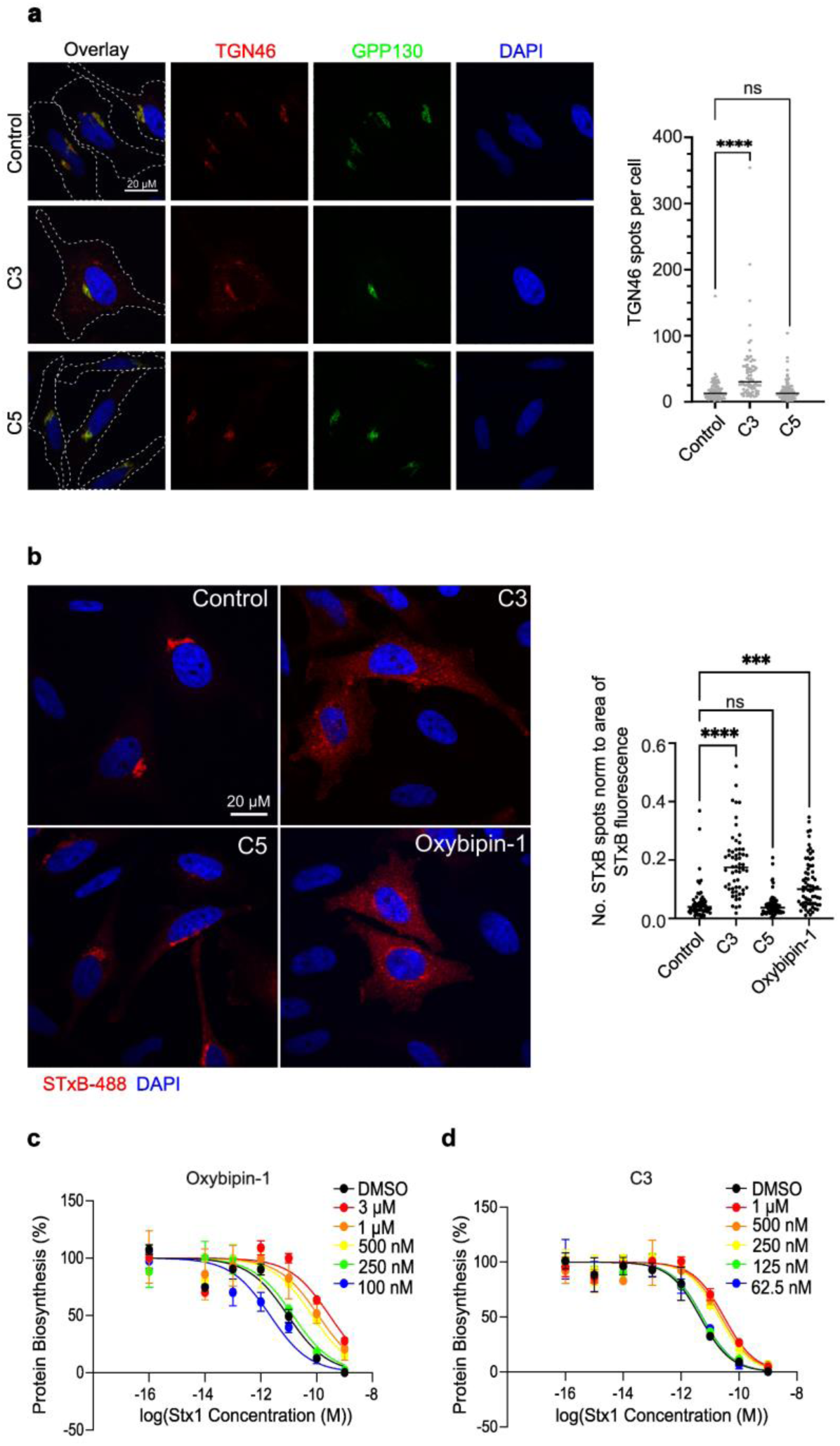
Retrograde trafficking of Shiga toxin was decreased by **oxybipin-1**. (a) Effect of **C3** on Golgi integrity. HeLa cells were treated for 4 h with 1 μM compounds as indicated. Golgi fragmentation was visualized by immunofluorescence using antibodies against the endogenous Golgi marker, TGN46 (red), GPP130 (green) and nuclei (DAPI). Scale bar = 20 μm. Graph shows quantification of number of TGN46 spots per cell, indicating Golgi fragmentation. Graph shows a minimum of 80 cells counted per condition from 3 independent experiments. Each spot represents one cell. Line indicates median. One-way ANOVA was performed (P <0.0001) with Dunnett’s multiple comparisons test, **** P<0.0001. (b) Effect on retrograde trafficking of Shiga toxin by **C3** and Oxybipin-1. Trafficking of STxB-488 (red) to the Golgi is decreased in the presence of **C3** and **Oxybipin-1** compared to **C5** or DMSO control. STxB spots (red) were counted in one central Z-slice per cell, correlating low numbers to STxB arrival at the Golgi, and high numbers correlating with the cytosolic spots that represent colocalization to early endosomes. Fluorescence area was measured in parallel and remained unchanged in all samples. Data is presented as number of STxB spots normalized to area of fluorescence. Scale bar = 20 μm. Graph shows a minimum of 52 cells counted per condition from 3 independent experiments. Each spot represents one cell. Line indicates median. One-way ANOVA was performed (P <0.0001) with Dunnett’s multiple comparisons test, **** P<0.0001, *** P = 0.0001. (c-d) Protection conferred by **oxybipin-1** (left) and **C3** (right) against Stx-1 protein biosynthesis inhibition (n = 3, representative experiment shown).

To study whether **C3** interacts with GPP130 directly, a cellular thermal shift assay (CETSA) with western blotting read-out was carried out. **C3** only exhibited a weak destabilization effect on GPP130 both in cell lysates (Figure 3b) and intact cells (Extended Data Fig. 6a). Additionally, isothermal dose–response fingerprinting (ITDRF) at 54.3 °C or 63.4 °C, where the largest differences between control and **C3** treated samples were observed, showed that **C3** failed to destabilize GPP130 in a dose-dependent manner at both temperatures (Extended Data Fig. 6b). Both CETSA and ITDRF indicate that **C3** does not bind to GPP130 directly and it is therefore likely that the degradation of GPP130 induced by **C3** is a secondary effect. This was further supported by the fact that QSOX2, another significantly down-regulated protein of **C3**, also exhibited a similar degradation pattern to GPP130. **C3** could also cause the degradation of QSOX2 in a dose-dependent manner and the mechanism of QSOX2 degradation was not mediated by the UPS but by the lysosome (Figure 3c). Additionally, **C3_Boc** and **C3_HyT** could also degrade QSOX2 at 1 μM (Figure 3d). Taken together, we postulated that the observed GPP130 or QSOX2 degradation is not due to direct binding of **C3** but a downstream effect of activity at its primary target.

### Glycosylation status and stabilization of Golgi proteins can be affected by C3 and its derivatives

Interestingly, by comparing the bands of GPP130 and QSOX2 by western blotting, it is noticeable that the electrophoretic mobility of GPP130 and QSOX2 are increased under the treatment of **C3** and its derivatives (Fig. 2d and e and 3a, and c-d). The observed band shift often indicates a change in post-translational modifications (PTMs) of the target protein. Based on the fact that both GPP130 and QSOX2 are highly glycosylated^24, 30^, we postulated that the shifts are due to incomplete or mis-glycosylation of GPP130 and QSOX2. To explore whether the *N*-linked glycosylation state was affected by **C3_HyT**, HeLa cells were treated with tunicamycin, a widely used *N*-glycosylation inhibitor to block the first step of biosynthesis of *N*-linked glycoproteins^31^. HeLa cell lysates were also treated with PNGase F to cleave most types of *N*-linked oligosaccharides from glycoproteins^30^. Clearly, band shifts for QSOX2 can be observed under all tested conditions which indicates that the glycosylation state of QSOX2 was successfully affected by tunicamycin and PNGase F (Figure 3e). Each condition yielded QSOX2 bands at different positions, indicating that they may affect the extent or the type of glycosylation. However, PNGase F and tunicamycin did not affect *N*-glycosylation of GPP130 under the same conditions (Extended Data Fig. 7).

Based on the fact that both GPP130 and QSOX2 are localized at the Golgi^24, 30^, we hypothesized that **C3** and its derivatives may affect the stability of other Golgi proteins. Therefore, expression proteomics was conducted to explore the degradation profiles induced by **C3_Me**, **C3_Boc** and **C3_HyT**. In total up to 6556 proteins were identified, where the expression of 37 proteins was significantly altered upon compound treatment (Figure 3f and Extended Dataset 2). **C3_HyT**, **C3_Boc,** and **C3_Me** exhibited nearly identical degradation profiles at 1 μM (Figure 3g). Additionally, the similarity of degradation profiles between **C3** and the negative control **C3_Me** further reconfirm that **C3** does not exert its function as a GPP130 degrader. Of note, manganese (II), which is reported to target GPP130 and leads to its oligomerization and degradation^19, 32–35^, exhibits a distinct degradation profile compared to **C3** derivatives, indicating that theGPP130 degradation mechanisms of Mn^2+^ and **C3** are different (Extended Data Fig. 8).

Moreover, the downregulated proteins induced by **C3** and its derivatives also exhibit similarities (Figure 3h). STRING functional enrichment analysis of the degraded proteins reveals an enrichment of Golgi proteins (16 of 23) and glycosyltransferases (MAN1A2, MAN2A1, GALNT2, EXTL3, GLT8D1 and GALNT6) which further support the fact that the increased mobility of GPP130 and QSOX2 is due to misglycosylation. In addition, several proteins (11 of 23) are also found in coat protein complex I (COPI) vesicles^36^. COPI is mainly responsible for trafficking cargos within Golgi and retrograde trafficking from the Golgi to ER. Therefore, the proteomics data also indicate that COPI-mediated retrograde trafficking was affected by **C3** and its derivatives. In summary, we hypothesized that these compounds interact with a Golgi component, affecting normal Golgi functions and leading to degradation of Golgi-resident proteins.

### C3 selectively targets OSBP

Having ruled out that **C3** and its derivatives interact with GPP130, and given that they contain a sterol scaffold, we hypothesized that the observed phenotypes may be due to the binding and possible inhibition of sterol transport proteins (STPs). In particular, oxysterol-binding protein (OSBP) localizes to ER-Golgi contact sites and acts as a cholesterol sensor to exchange cholesterol/PI4P between the ER and the Golgi^13^. Additionally, the inhibition or depletion of OSBP has been reported to lead to the reduction of mannosidase II (Man II) and the mislocalization of COPI vesicles^37^ as well as Golgi fragmentation and impaired Golgi trafficking^38–40^. Due to the similarities in the sterol binding domains of OSBP and other STPs^9, 12, 41–46^, we sought to establish an STP screening panel comprising ORP2 as a representative member of the ORP family 2, STARD1, as a representative member of the STARD family as well as Aster-A, Aster-B and Aster-C.

To establish a screening and selectivity panel, we developed competitive fluorescence polarization (FP) assays employing 22-NBD-cholesterol as a tracer molecule. The ORD of OSBP(377-807) was expressed harboring a GST tag to maintain sufficient stability and solubility of the protein. Importantly, GST alone does not interfere with 22-NBD-cholesterol binding (Figure 4a). The sterol binding domains of ORP2 and STARD1 were expressed with an N-terminal His6 tag and the ASTER domain of the Aster proteins were also expressed with a GST tag to improve their solubility. During the purification process the GST tag was removed via protease cleavage^47, 48^. GST-OSBP(377-807), ORP2, STARD1 and Aster proteins were titrated against 20 nM 22-NBD-cholesterol to determine the optimal assay protein concentration, which is generally between 60% and 80% saturation to obtain an appropriate assay window. 22-NBD-cholesterol efficiently binds to all STPs (Figure 4a) indicating its suitability as a FP assay tracer molecule. The calculation of Z’ factors clearly indicated that the competitive FP assay for the selected STPs is robust and suitable for the evaluation of ligand binding affinity (Extended Data Fig. 9). We screened **C3_Boc, C3_HyT** towards OSBP, ORP2, STARD1 and the Aster proteins at concentrations of 1 µM (Figure 4b) and 3 µM (Extended Data Fig. 9). **C3_Boc** and **C3_HyT** completely inhibited GST-OSBP(377-807) binding to 22-NBD-Chol at 1 µM, similarly to the endogenous OSBP ligand 25-OH Cholesterol (25-OHC) and the known inhibitor **OSW-1**, while not showing any inhibition of the other sterol transporters. As such, we renamed them **oxybipin-1 (C3_HyT)** and **-2 (C3_Boc)**, for oxysterol binding protein inhibitors, respectively. We subsequently screened **oxybipin-1,** and **-2** as well as **C5, 25-OHC** and **OSW-1** towards GST-OSBP(377-807) in a dose-dependent manner (Figure 4b). Both **oxybipin-1** (IC50 = 142 nM) and **oxybipin-2** (IC50 = 74 nM) were potent inhibitors of the OSBP 22-NBD-Chol interaction.

Following this, we aimed to determine target engagement of OSBP and related ORPs in a cellular setting. For this, a cellular thermal shift assay (CETSA) in a split nanoluciferase (HiBiT-LgBiT) system in KBM7 cells overexpressing HiBiT-tagged OSBP, as well as the closely related ORP4L and ORPs 2 and 9 was carried out^49^. Consistently with the biochemical data, **oxybipin-1** and **oxybipin-2** exhibited a significant stabilization effect on OSBP with ΔT > 5 °C while **C5** only exhibited an insignificant stabilization effect on OSBP with ΔT ∼ 1 °C, which further supports the interaction between the **oxybipins** and OSBP in live cells. Notably, no stabilizing effect of the **oxybipins** could be observed on other ORPs including the highly structurally related ORP4L (Figure 4d), as well as ORP2 and 9 (Extended Data Fig. 9c). Selectivity amongst related sterol transport proteins is challenging due to the similarity in the sterol binding domain, however a recent report highlighted that different oxysterols have differential binding preferences to OSBP and ORP4 based on the site and extent of their oxidation pattern^50^. Our compounds may mimic this effect with the position of the amide and ether heteroatoms on the sterol side chain.

OSBP inhibition by a set of unrelated natural products termed the ORPphilins^15^ has been shown to produce diverse effects on OSBP and ORP4 protein levels, with compounds including **OSW-1** leading to OSBP degradation. OSBP and ORP4S levels were not affected by the treatment of 1 μM **oxybipin-1** and **-2**, unlike the control **OSW-1**, which led to OSBP degradation (Figure 4e). This suggests that different mechanisms operate for **OSW-1** and the **oxybipins**.

### Retrograde trafficking of Shiga-toxin was impaired by C3

In line with our proteomics data, **C3** induced partial fragmentation of the Golgi, which confirms that in some contexts,^40^ inhibition of OSBP affects Golgi integrity (Figure 5a). To test whether protein trafficking in the cell was perturbed, we used classical retrograde transport cargo Shiga toxin, to assess whether OSBP inhibition would affect the efficiency of retrograde transport. The transportation of Shiga toxin starts from being recognized by globotriaosylceramide (Gb3) at the plasma membrane. Shiga toxin is then transported via retrograde trafficking from the early endosomes to the Golgi and finally to the endoplasmic reticulum (ER) to exert its toxicity by inhibiting ribosomes and stopping protein synthesis. If the retrograde trafficking of Shiga toxin is inhibited, it will be transferred from early endosomes to the lysosome for degradation or be directed to recycling endosomes to be expelled from the cell in extracellular vesicles^51^. In HeLa cells in control conditions, after 40 minutes of uptake at 37 °C, the majority of a fluorescently labelled Shiga toxin B-subunit (STxB), had reached the Golgi (Figure 5b). However, for cells treated with **C3** and **oxybipin-1**, the STxB was still trapped in early endosomes due to impaired retrograde trafficking^52^. This was not the case for negative control **C5**, which was similar to control conditions. To further explore whether OSBP inhibition could protect cells against Shiga toxin toxicity, we compared the protein biosynthesis rate in the presence of Shiga toxin-1 (Stx1) with and without **C3** and **oxybipin-1** treatment. The protein synthesis rate was measured by the incorporation of radiolabeled [^14^C]-leucine into newly synthesized proteins. **Oxybipin-1** exhibited a 14-fold protection at 3 μM (Figure 5c) and **C3** exhibited a 5-fold protection at 1 μM (Figure 5d), while **C5** did not exhibit any protection (Extended Data Fig. 10). In summary, the inhibition of OSBP by **C3** and **oxybipin-1** affected the integrity of the Golgi apparatus and impaired the retrograde trafficking of Shiga toxin, thereby further protecting cells against Shiga toxin toxicity.

## Discussion

Here, we describe the identification of a highly potent and selective tool compound for inhibiting OSBP, which inhibits retrograde trafficking and Shiga toxin cytotoxicity. While the original objective of this work was to identify the potential targets of sterols via targeted protein degradation through the synthesis of sterol-containing chemical chimeras, we found that the degradation of GPP130 and a large series of Golgi proteins was a downstream effect of OSBP inhibition, rather than degradation. The lysosomal degradation of Golgi proteins and altered glycosylation states, is likely the source of the observed partial Golgi fragmentation and inhibition of retrograde trafficking. Importantly, inhibition of retrograde trafficking is a powerful strategy for inhibiting Shiga toxin toxicity. As such, we provide evidence that the inhibition of OSBP could block retrograde trafficking of Shiga toxin and protect cells against Shiga toxin toxicity.

The identification of an OSBP inhibitor rather than a degrader demonstrates the challenges of applying PROTACs for target identification. It is challenging to discriminate whether any observed target degradation is the result of a direct binding event of the PROTAC or a secondary effect. The difference, or lack thereof, in effect between inhibition and degradation has also been widely documented for PROTACs derived from inhibitors, rather than binders. Furthermore, the intrinsic selectivity of PROTACs further hinders its application in target identification, in particular for promiscuous molecules such as sterols. Different linkers, E3 ligands and linker attachment positions may increase the stability of the protein-of-interests(POI)-PROTAC-E3 ternary complex and further affect the degradation efficiency^53^, however this level of optimisation is prohibitive for a target ID project. Finally, the test conditions, including time and concentrations of the applied PROTAC will also affect its degradation profile. The optimal concentration can vary depending on the POI. Therefore, it will be challenging to find an optimal PROTAC concentration to trigger the significant degradation of all proteins while avoiding the Hook effect^54^.

Although TPD was not a viable strategy for sterol target ID, the identification of selective OSBP inhibitors still provide highly useful tools to study the function of OSBP, as well as a potential as therapeutic leads. So far only a small number of STP inhibitors have been identified, mostly serendipitously, and their selectivity profiles have not been studied. The natural products termed ORPhilins, including cephalostatin 1, OSW-1, ritterazine B and schweinfurthin A, target OSBP as well as the structurally highly similar ORP4L^15^. However, isolation of these compounds from natural sources yields small amounts, and the total synthesis is highly challenging^55^. Our HiBit CETSA and recombinant protein assays suggest that the oxybipins are highly selective for OSBP, even over the highly similar ORP4L. The establishment of a biophysical selectivity panel utilizing fluorescence polarization assays will enable the profiling of all future STP inhibitors, enabling the standardization of activity and selectivity, which was a significant challenge until now. In the future, the inclusion of other sterol binding domains of STPs as well as the full-length proteins will be necessary to broaden the biophysical selectivity panel and to consolidate OSBP as the main target of oxybipins.

OSBP inhibitors have potential applications in a range of diseases from activity against enteroviruses to specific toxicity towards selected cancer cell lines. Here, we extend the possible applications to the attenuation of Stx cytoxicity. Currently, the treatment strategies of infections by Stx-producing *E. coli* include the neutralization of Stx using antibodies^56, 57^, the inhibition of the binding of Stx by inhibiting its interaction with the plasma membrane receptor Gb3^58^ and the blockage of the intracellular trafficking route of Stx by manganese^19^ or other small molecule inhibitors such as Retro-1/2^59–61^. It is worth noting that antibiotics are generally contraindicated as they lead to sudden release of high amounts of toxins, leading to dangerous complications^62^. Our work further provides an alternative treatment strategy to impair the retrograde trafficking of Stx by targeting OSBP. Oxybipins exhibit a significant protective effect against Stx at 1 μM while no cytotoxicity of oxybipins can be observed at 10 μM suggesting a potential therapeutic window for these compounds, which warrants further studies.

## Acknowledgements

The Laraia Lab was supported by funding from the Novo Nordisk Foundation (NNF19OC0058183, NNF21OC0067188) and the Independent Research Fund Denmark (9041-00241B, 9041-00248B). CeMM and the Winter Lab are supported by the Austrian Academy of Sciences. The Winter lab is further supported by funding from the European Research Council (ERC) under the European Union’s Horizon 2020 research and innovation program (grant agreement 851478), as well as by funding from the Austrian Science Fund (FWF, projects P32125, P31690 and P7909). The Gillet lab was funded by the French National Research Agency (ANR) under Contracts LeishmaStop ANR-18-CE18-0016-01 and SMERSEC ANR-20-CE18-0016-01, by University Paris Saclay under Contracts ReCoVEr and PIMVir, and the Joint ministerial program of R&D against CBRNe risks. We thank Ludger Johannes for helpful discussions.

## Author contributions

N.H. and L.L. designed the project. N.H. carried out the compound synthesis, proteomics sample preparation and analysis, and cell biological validation of compounds. L.D. expressed and purified recombinant proteins and developed and performed fluorescence polarization assays. C.R. designed, supervised, performed and analysed proteomics experiments and supported cell biological validation. O.R.D. synthesized vHL probes and conducted their biological analysis. A.F. performed the Golgi integrity and retrograde trafficking experiments. M.C. performed the HiBit CETSA experiments under supervision of G.E.W. M.M. performed the Stx1 inhibition experiments under the supervision of J.B. and D.G. J.H. supported recombinant protein expression and synthesis. L.L. supervised the project. N.H., L.D. and L.L. wrote the paper with input from all authors.

## Extended data

**Extended Data Fig. 1.**
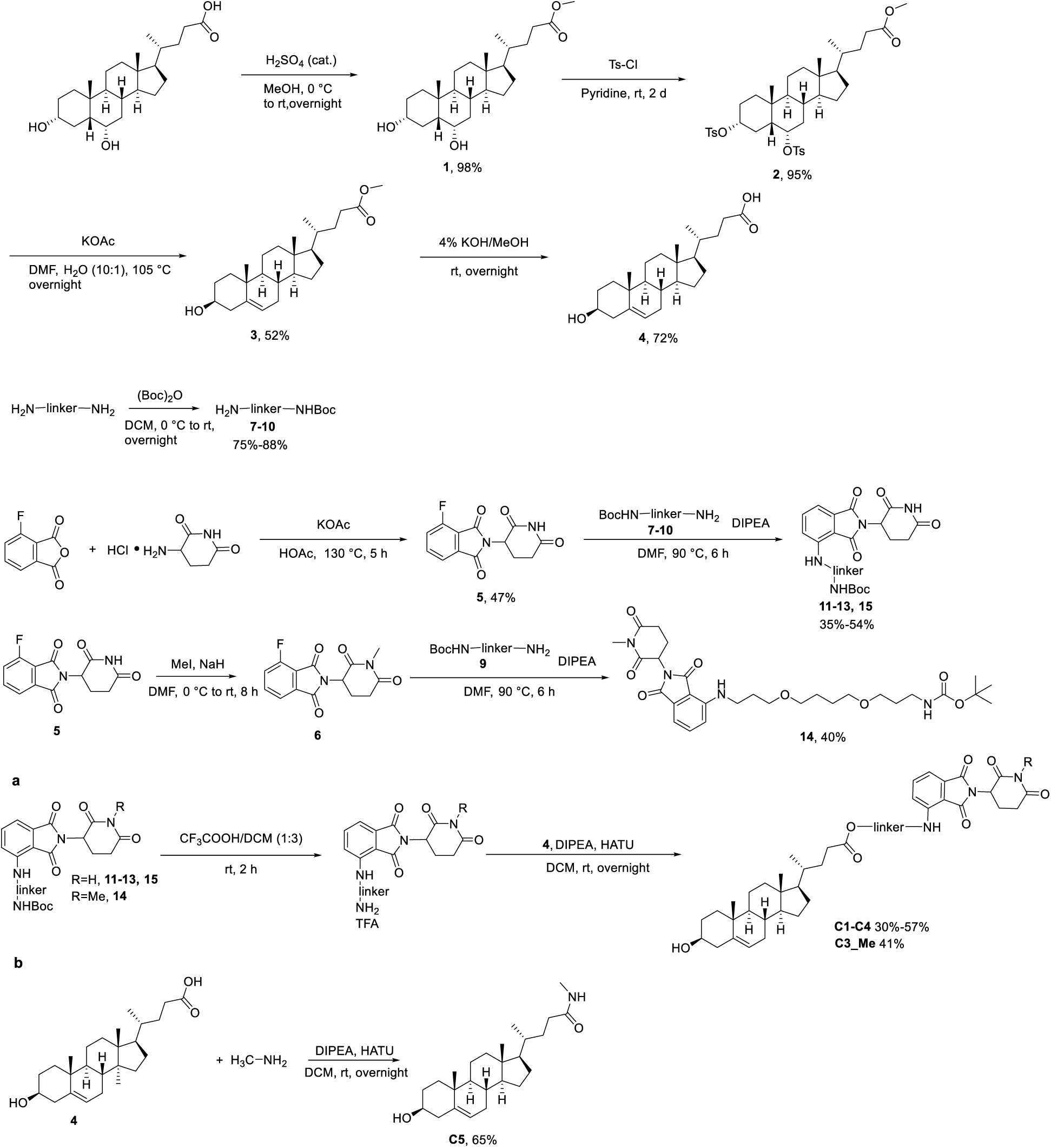
Synthesis of Cholesterol-bearing PROTACs **C1-C4**, **C3_Me** (a) and **C5** (b).

**Extended Data Fig. 2.**
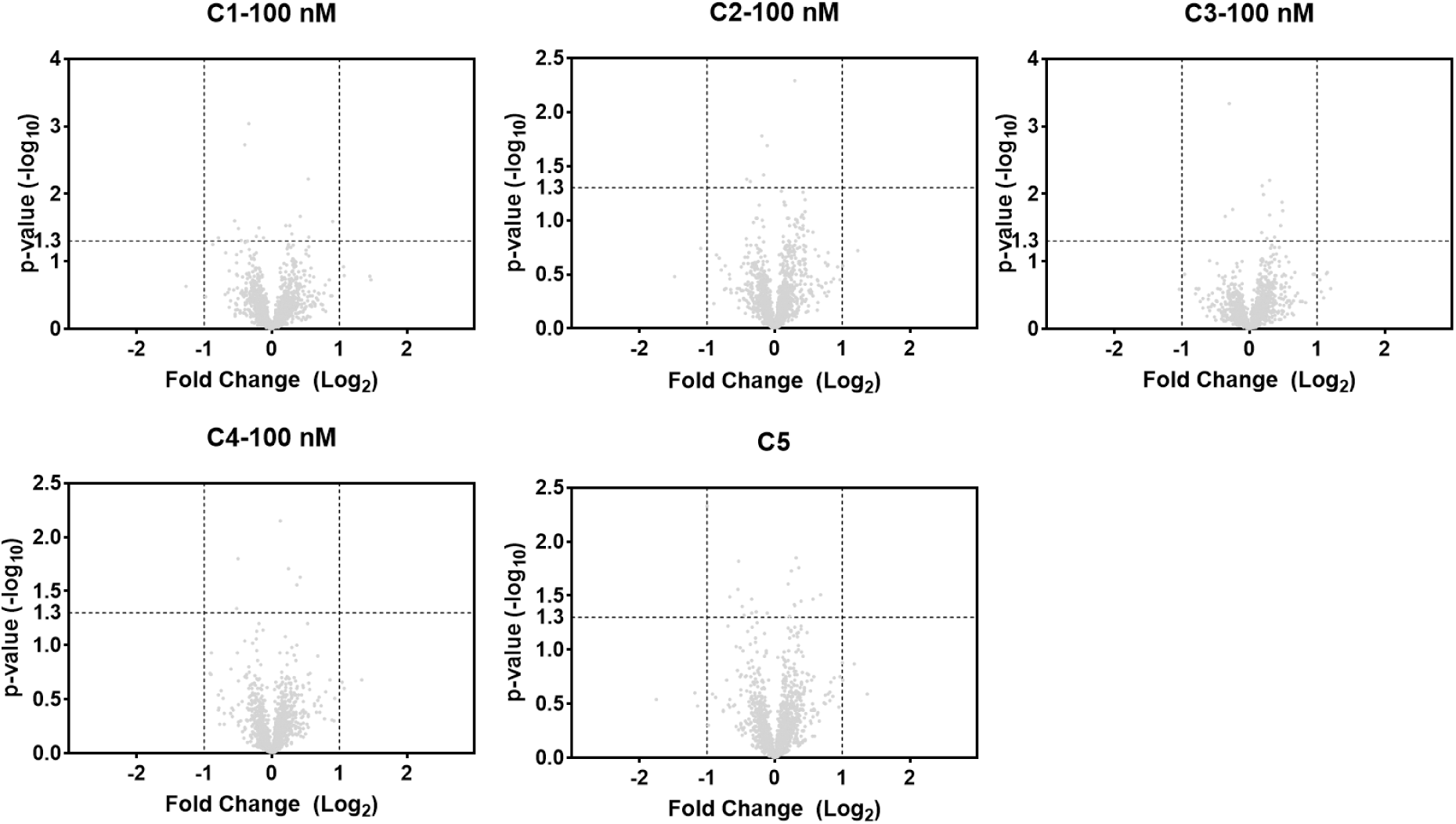
Degradation profiles of **C1-C4** at 100 nM and **C5** at 1 μM in HeLa cells treated for 18 hours. Identified proteins were plotted as log2 fold change (compound-treated /DMSO-treated) versus - log10 (p-value). P-values were calculated from the data of three technical replicates. The dotted horizontal line marks the significance threshold of p = 0.05 and dotted vertical lines represent the threshold of an absolute log2 fold change of 1.

**Extended Data Fig. 3.**
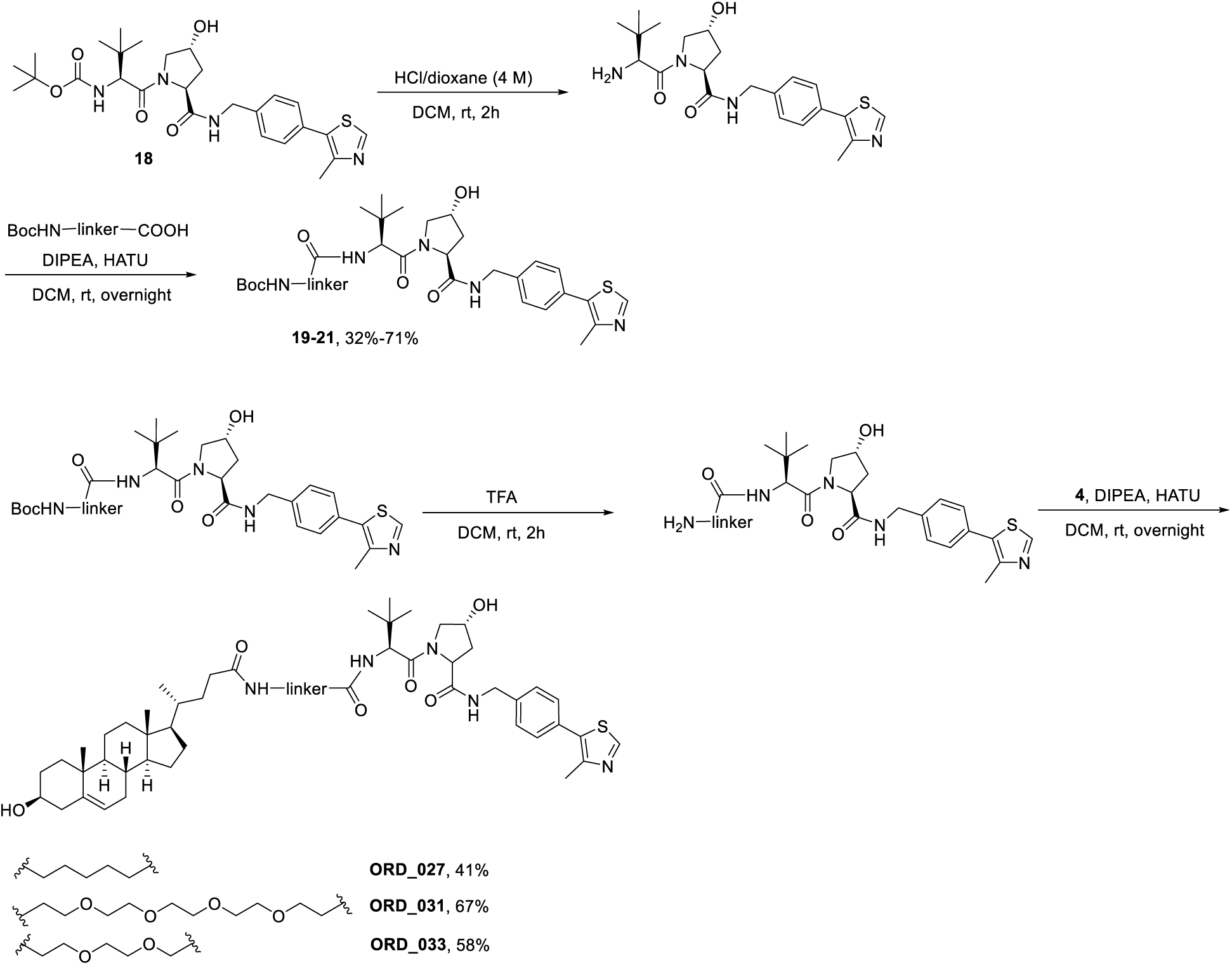
Synthesis of VHL-based PROTACs, **ORD_027**, **031** and **033**.

**Extended Data Fig. 4.**
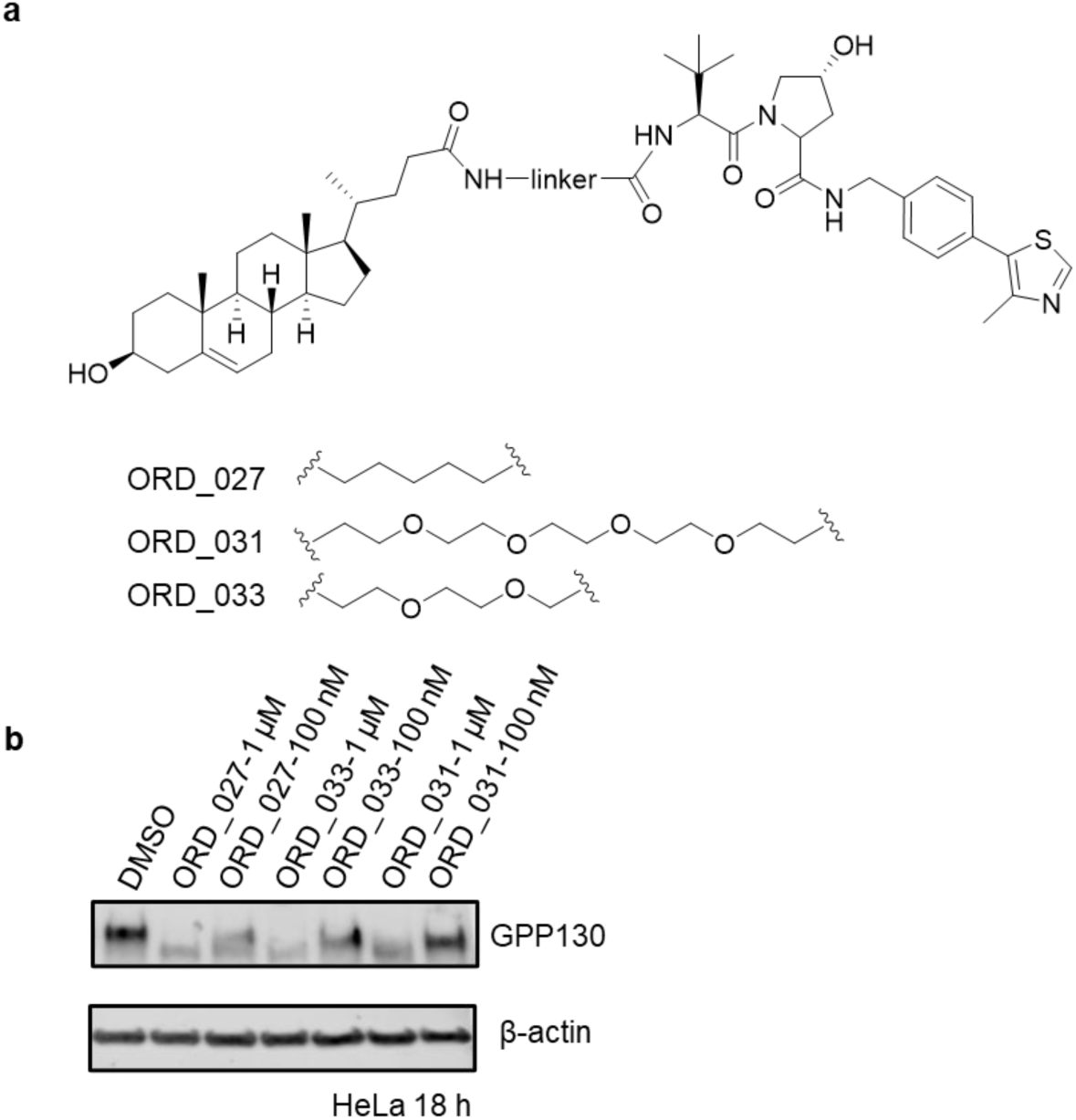
GPP130 can be degraded by VHL-based PROTACs. (a) Structures of VHL-based PROTACs, **ORD_027,031** and **033**. (b) Effect of VHL-based PROTACs **ORD_027,031** and **033** on GPP130 levels in HeLa cells for 18 hours (n = 3).

**Extended Data Fig. 5.**
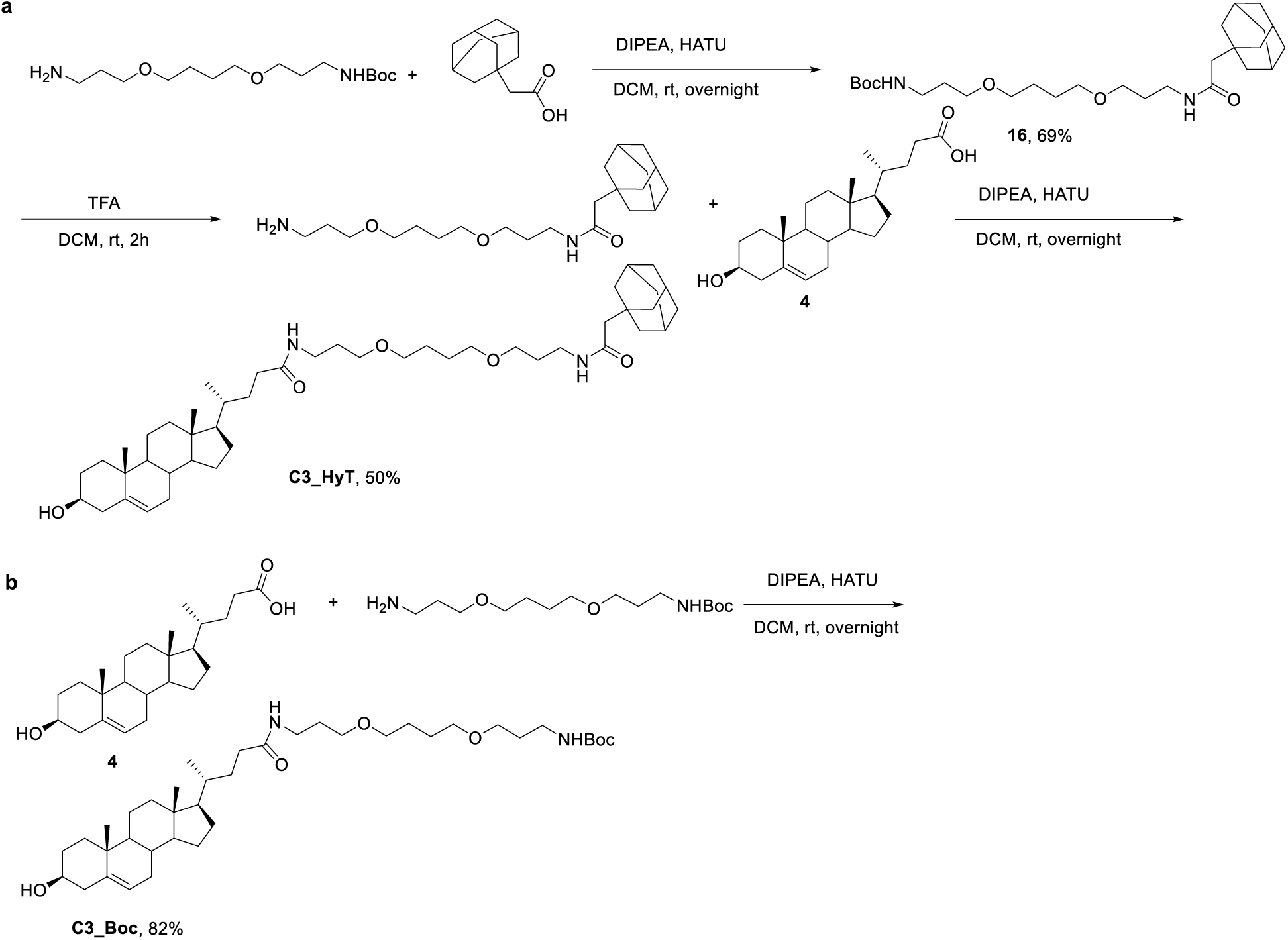
Synthesis of **C3_HyT** (a) and **C3_Boc** (b).

**Extended Data Fig. 6.**
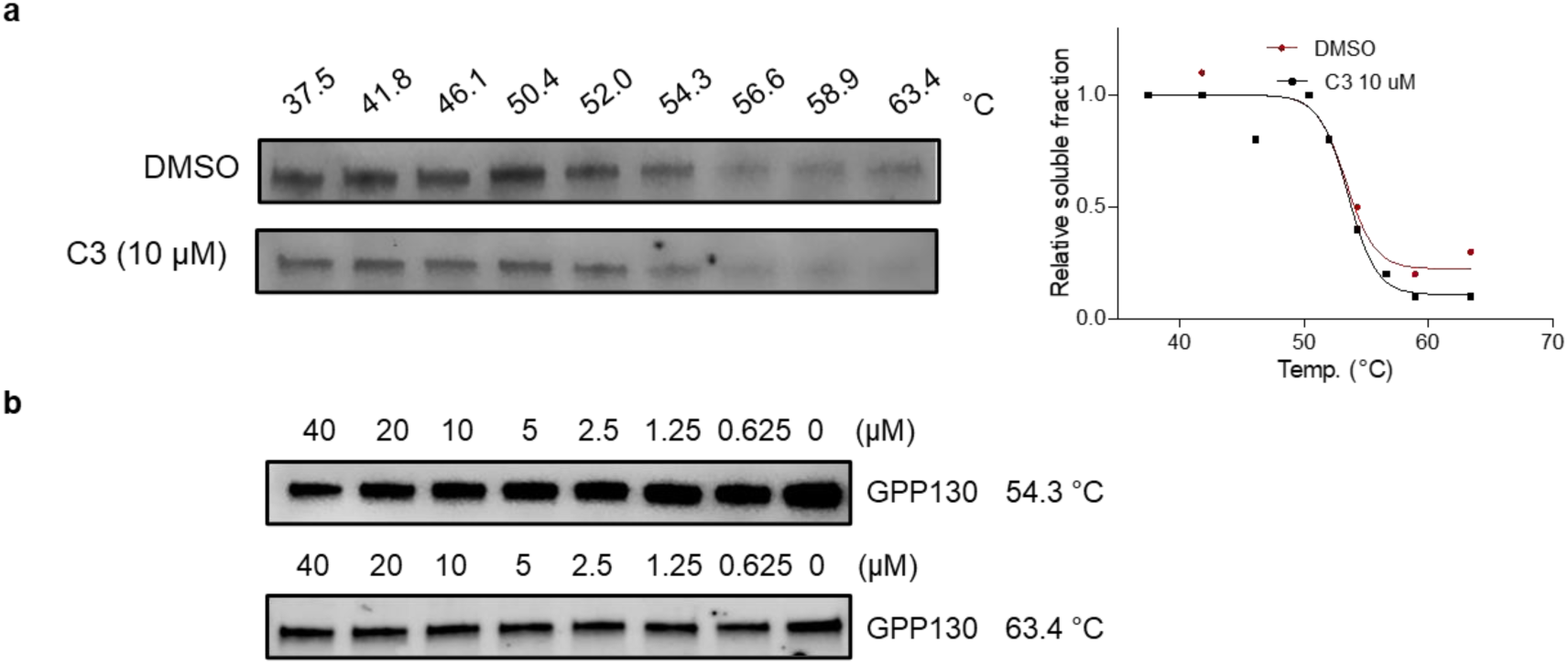
**C3** does not bind to GPP130 directly. (a) Cellular thermal shift assay shows **C3** exhibits a weak destabilization effect on GPP130 in intact HeLa cells (n = 2). (b) Isothermal dose response fingerprints assay shows **C3** failed to destabilize GPP130 in a dose-dependent manner at 54.3 ℃ (n = 3) and 63.4 ℃ (n = 1) in intact HeLa cells.

**Extended Data Fig. 7.**
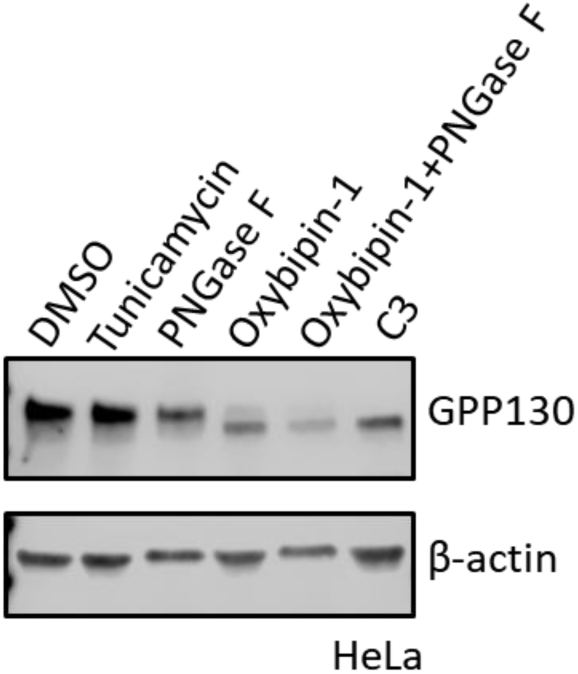
Effect of Tunicamycin, PNGase F, **C3** and **Oxybipin-1** on the *N-*glycosylation of GPP130 (n = 2). HeLa cells were treated with tunicamycin (5 μg/mL) for 18 hours (n = 2).

**Extended Data Fig. 8.**
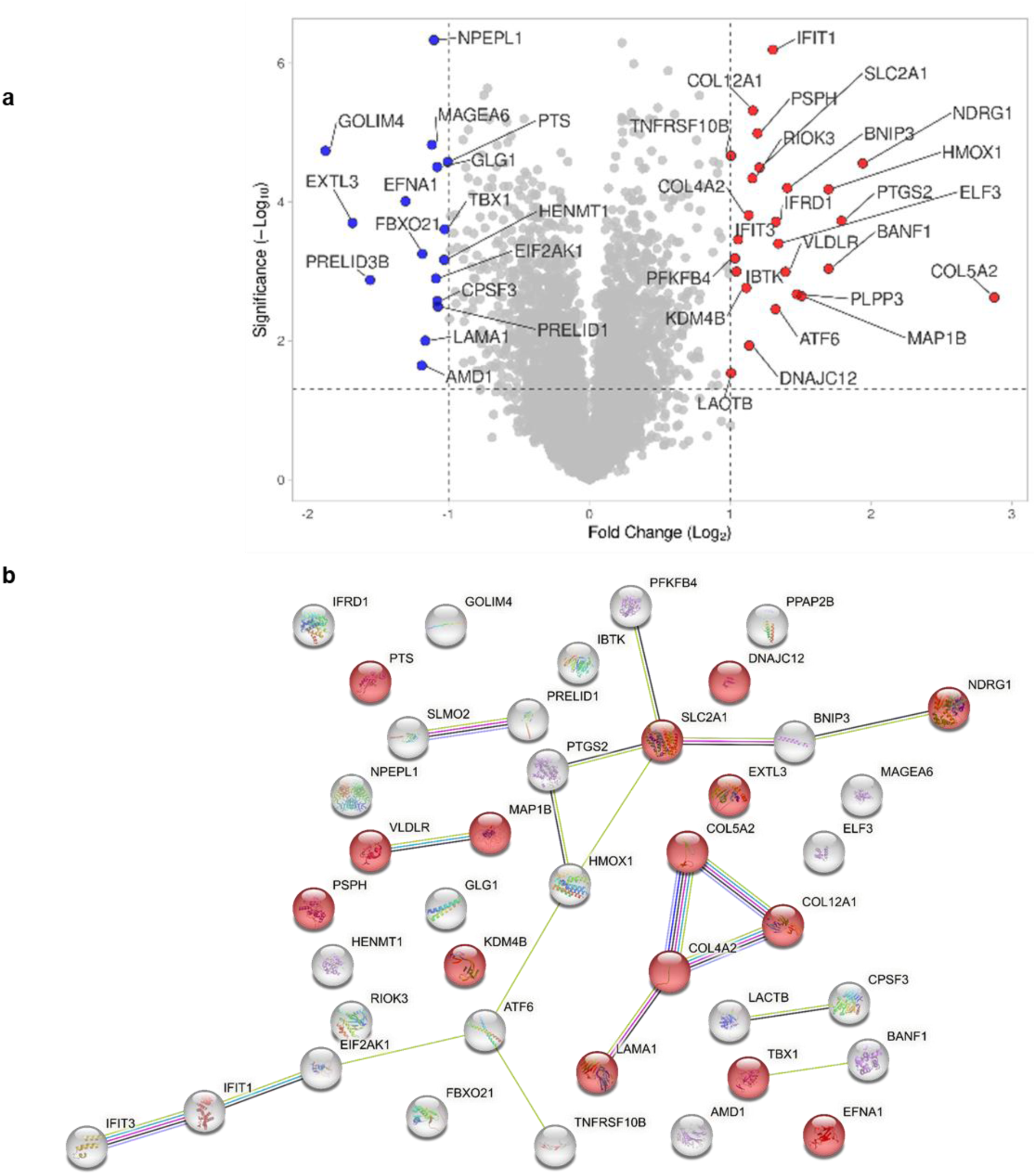
Degradation profiles of manganese(II) (a) Degradation profiles of manganese(II) at 150 μM in HeLa cells. The down-regulated proteins (blue dots) and up-regulated proteins (red dots) were plotted as log2-fold change (Mn(II)-treated/DMSO-treated) versus -log10-(p-value). P-values were calculated from the data of three technical replicates. The dotted horizontal line marks the significance threshold of p = 0.05 and dotted vertical lines represent the threshold of an absolute log2 fold change of 1..(b) STRING analysis of affected proteins by Mn^2+^ exhibits an enrichment of proteins involved in neurodevelopmental delay (Red dots; HP:0012758; FDR: 0.0419)

**Extended Data Fig. 9.**
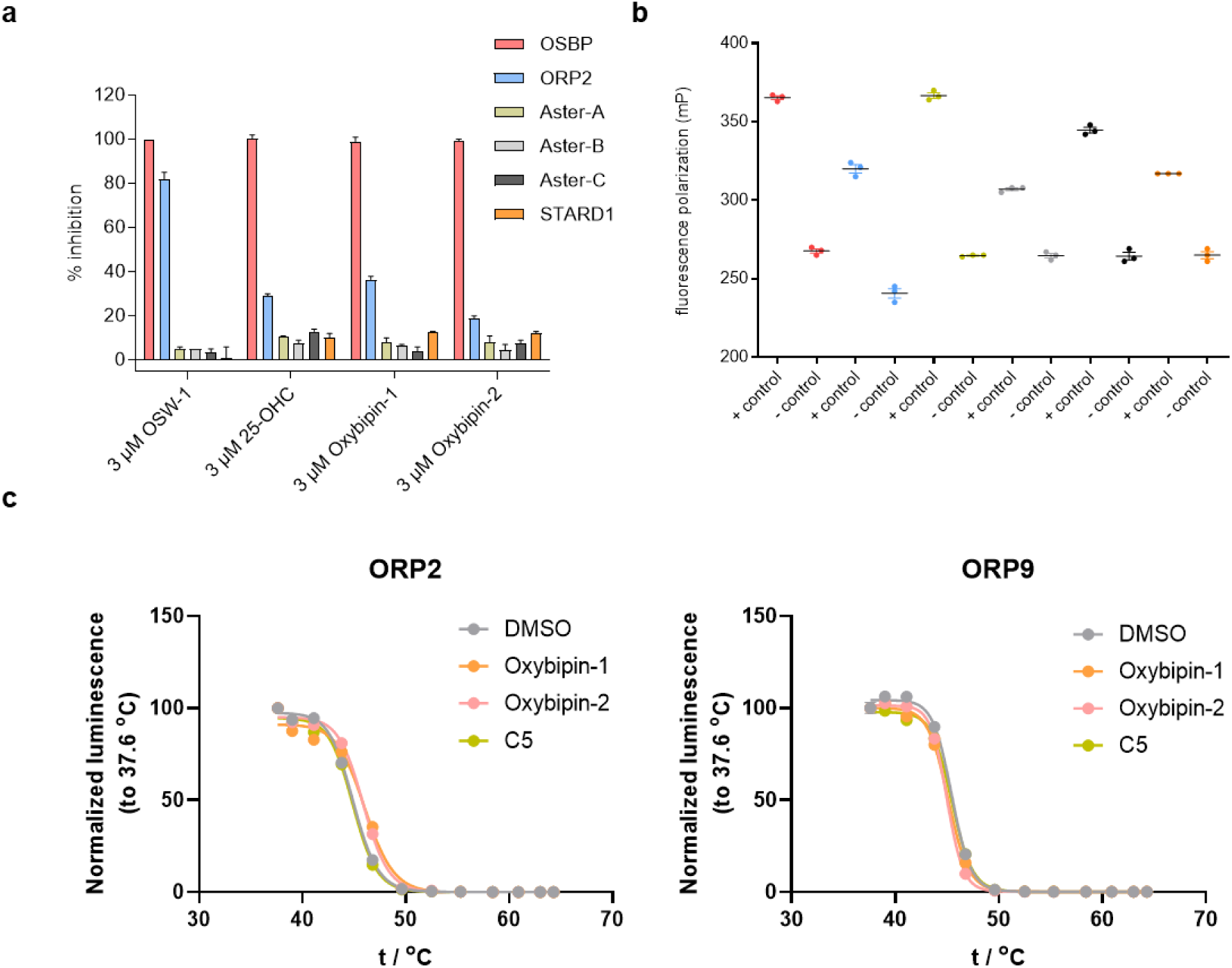
In vitro binding to sterol transport proteins. (a) **oxybipin-1** and **oxybipin-2** profiling against ORPs, STARDs and Asters at 3 μM (n = 3). 25-OH cholesterol and **OSW-1** are control OSBP inhibitors. (b) Assay controls for fluorescence polarization assays (n = 3); red = OSBP, blue = ORP2, green = Aster-A, light grey = Aster-B, dark grey = Aster-C, orange = STARD1. (c) Effect of **oxybipin-1** and **oxybipin-2** on the thermal stability of ORP2 and 9 in a HiBit CETSA experiment (n = 3).

**Extended Data Fig. 10.**
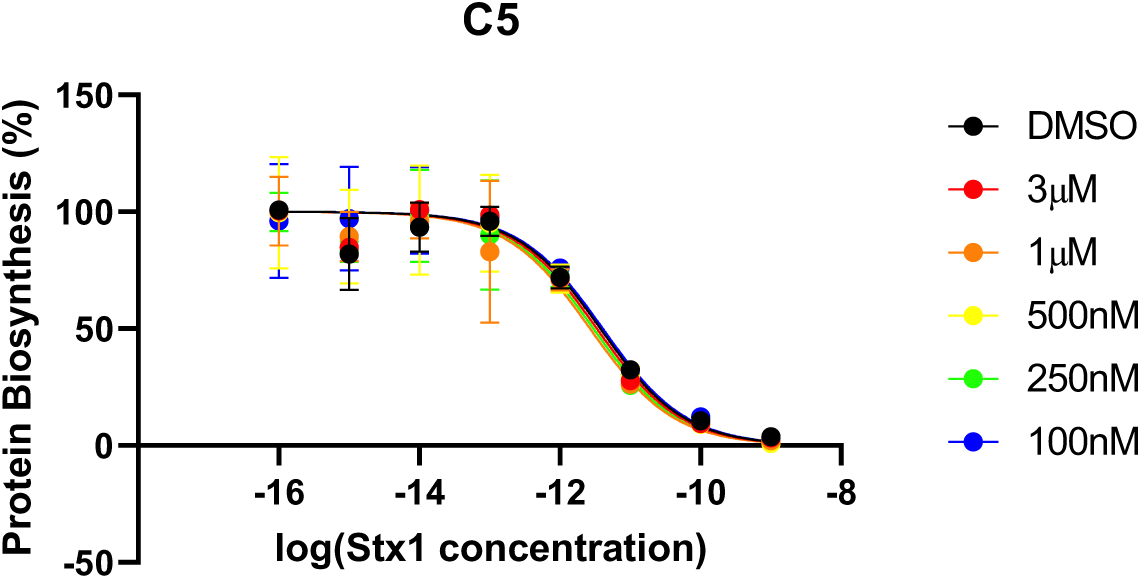
No protection effect against Shiga toxin can be observed under the treatment of **C5**.

## Methods

### Proteomics

#### Reagents and suppliers

| Reagent or resource | Source | Identifier |
| --- | --- | --- |
| **Biological reagents** |  |  |
| Dulbecco's Modified Eagle's Medium (DMEM) | Gibco | 41966-029 |
| Fetal bovine serum (FBS) | Gibco | 11563397 |
| Penicillin/streptomycin (P/S) | Gibco | 11548876 |
| Nonidet P40 Substitute (NP-40) | Sigma-Aldrich | 11754599001 |
| Triethylammonium bicarbonate buffer (TEAB) | Sigma-Aldrich | T7408 |
| Tris(2-carboxyethyl)phosphine hydrochloride (TCEP) | Sigma-Aldrich | C4706 |
| Iodoacetamide (IAA) | Sigma-Aldrich | I1149 |
| Acetone | Sigma-Aldrich | 650501 |
| Trypsin | Sigma-Aldrich | T6567 |
| TMT 11plex reagents kits | Thermo Scientific | A34808 |
| Hydroxylamine for TMT experiments | Thermo Scientific | 90115 |
| Acetonitrile | Pierce | 51101 |
| Trifluoroacetic acid (TFA) | Honeywell Fluka | 14264 |
| Ammonium hydrogen carbonate | Supelco | 533005 |
| 0.1% Formic Acid | Pierce | 85174 |
| **Materials** |  |  |
| Sep-Pak® Plus C18 cartridges | Waters | WAT020515 |

#### Cell culture

A549 and HeLa cells were cultured in DMEM supplemented with 10% FBS and 1% P/S at 37 °C with 5% CO_2_.

#### Cell lysis and sample preparation for MS analysis

2 mL cells (Cell density 2 ×10^5^ cell/ mL) were seeded in 6-well plates and incubated overnight at 37 °C and 5% CO_2_. The following day, cells were incubated with the corresponding compound (or DMSO for the control sample) for 18 hours at 37 °C and 5% CO_2_. At the end of the incubation, cells were washed with ice-cold PBS twice and lysed by the addition of lysis buffer consisting of 0.4% (v/v) NP-40 in PBS on ice. Lysis was finalized by four freeze-thaw cycles using liquid nitrogen. Cellular debris were removed by centrifugation at 25000 × g for 25 minutes at 4 °C. The protein concentration of the cell lysate was measured by DC assay (BioRad) and further diluted to a protein concentration of 2 mg/mL in lysis buffer. For each sample, 150 µg of total proteins were further diluted in 100 mM TEAB, and with 200 mM TCEP for cysteine reduction for 1 h at 55 °C. 375 mM IAA was successively used for alkylation for 30 min at room temperature in the dark. After this step proteins were precipitated overnight with 6 volumes of cold acetone at -20 °C. The following day the protein pellets were collected by centrifugation at 8000 × g at 4 °C for 10 min and resuspended in TEAB 100 mM before digestion. Protein digestion was performed by adding trypsin in a 1:80 enzyme: substrate ratio and left overnight at 37 °C, with gentle shaking in a ThermoMixer (Eppendorf®). Half of the sample volume corresponding to 75 µg of peptides were successively labelled with TMT 11-plex reagents kits. Experimental replicates were labelled with different TMT-batches concatenated by a pooled internal reference sample for each TMT-batch, which was consisting of an additional sample obtained by pooling equal amounts of each sample from the same TMT-batch. TMT reaction was allowed for 2 hours and quenched with 5% hydroxylamine. All the 11 samples related to each batch were pooled and dried in SpeedVac (Eppendorf EP022822993) before desalting with Sep-Pak® Plus C18 cartridges. Peptide were eluted with 40% and 60% of acetonitrile in 0.1% of trifluoroacetic acid (TFA) and dried before injection of 30 µg in the UHPLC system (Dionex U3000) for high-pH fractionation.

The separation of the peptides was carried out at a constant flowrate of 5 µl min^-1^ on a CSH C18 Acquity UPLC M-Class Peptide column, 130 Å, 1.7 µm, 300 µm × 150 mm (Waters, 186007563) using a 100 min linear gradient from 5 to 35% of mobile phase B (acetonitrile) with a subsequent 15 min gradient to 70%, before 5 min re-equilibration with 95% of mobile phase A (5mM ammonium bicarbonate, pH 10). 60 time-based fractions were pooled in 30 fractions in the collection plates. Clean-up of the fractions was performed by EvoTip according to the manufacturer’s instructions.

#### LC-MS analysis

The EvoTips (EvoSep, EV2003) were loaded on the Evosep One module (EvoSep EV-1000) coupled to an Orbitrap Eclipse™ Tribid™ mass spectrometer (ThermoFisher Scientific). Peptides were loaded onto the EASY-Spray™ C18 column, 2 µm, 100 Å, 75 µm × 15 cm (ThermoFisher Scientific, ES804) using the standard “30 samples per day” Evosep method. The method eluted the peptides with a 44 min gradient ranging from 5% to 90% acetonitrile with 0.1% formic acid. The MS acquisition was performed in data dependent-MS3 with real-time-search (RTS) and a FAIMS interface switching between CVs of −50 V and −70 V with cycle times of 2 s and 1.5 s, respectively. The data dependent acquisition mode was run in a MS1 scan range between 375 and 1500 *m/z* with a resolution of 120000, and a normalized gain control (AGC) *target* of 100%, with a maximum injection time of 50 ms. RF Lens set at 30%. Filtering of the precursors was performed using peptide monoisotopic peak selection (MIPS), including charge states from 2 to 7, dynamic exclusion of 120 s with ±10 ppm tolerance excluding isotopes, and a precursor fit of 70% in a windows of 0.7 *m/z* with an intensity threshold of 5000. Selected precursors for further MS2 analysis were isolated with a window of 0.7 *m/z* in the quadrupole. The MS2 scan was performed over a range of 200-1400 *m/z*, collecting ions with a maximum injection time of 35 ms and normalized AGC target of 300% MS2 fragmentation was operated with normalized HCD collision energy at 30%. Fragmentation spectra were searched against the fasta files from the human Uniprot database (reviewed) in the RTS, set with tryptic digestion, TMT-11plex as fixed modification on Lysine (K) and N-Terminus together with cysteine (C) carbamidomethylation, and oxidation of methionine (M) as variable modification. 1 missed cleavage and 2 variable modifications were allowed with a maximum search time of 35 ms. FDR filtering was enabled with 1 as Xcorrelation and 5 ppm of precursor tolerance. Precursors identified via RTS in the MS2 scan were further isolated in the quadrupole with a 2 *m/z* window, maximum injection time of 86 ms and normalized AGC target of 300%. The further MS3 fragmentation was operated with a normalized HCD collision energy at 50% and fragments were scanned with a resolution of 50000 in the range of 100 to 500 *m/z*. The MS performances were monitored by quality control of an in-house standard of HeLa cell lysate, both at the beginning and the end of each sample set.

#### Data analysis

Mass spectrometric raw files were analyzed with MaxQuant (version 1.6.2.10, https://maxquant.org/) setting all the TMT11 labels as reporter ion MS3 for lysine and N-terminus with mass tolerance 0.003 Da. The search settings included carbamidomethylation of cysteine (C) as fixed modifications, methionine (M) oxidation and acetylation of protein N-termini as variable modification and trypsin as enzyme (allowing maximum 2 missed cleavages). Peptide-spectrum match (PSM) and protein false discovery rates (FDR) were 0.01 with at least 1 minimum razor + unique peptides. Peptide sequences were searched against UniProtKB/ TrEMBL human proteome6 fasta file (downloaded on 08/08/2018, https://www.uniprot.org/proteomes/). The protein group file generated from the MaxQuant analysis were processed for further normalization and scaling.

Contaminants and proteins identified in the reverse database were removed, as well as all the proteins identified with a sum of Unique + Razor peptides below 2. Loading normalization was performed on the first 10 TMT channels, summing the intensities corrected for isotopic error for each TMT channel and calculating the respective correction factor on the average of the summed intensities. Hence, each protein intensity was normalized for the respective channel correction factor. For comparison of the experimental replicates measured in the multi-batch TMT experiment, the internal reference scaling (IRS) normalization method was applied following the procedure described in Plubell et al^63^. Briefly, the geometric mean of the channels measuring the pooled internal reference sample in the 11th channel was calculated across the replicates and an inter-batch scaling factor was calculated and applied to each protein.

Normalized intensities were further considered to calculate the average among the replicates and the fold changes (FC) for PROTACs-treated samples over the control samples. The obtained values were log2 transformed. A two-sided T-test was performed on the normalized data and obtained p-values were -log10 transformed.

Volcano plots were obtained plotting the log2 Fold Change vs -log10 (p-value) in both Prism 5 (Graphpad Software, Inc. Version 5.3) and VolcanoseR^64^.

### Immunoblotting

#### Reagents and suppliers

| Antibodies |  |  |  |
| --- | --- | --- | --- |
| Products | Source | Identifier | Dilution |
| GOLIM4 Polyclonal Antibody | Invitrogen | PA5-51624 | 1: 1000 |
| Anti-QSOX2 antibody | Abcam | ab121376 | 1: 1000 |
| Anti-OSBP antibody produced in rabbit | Sigmaaldrich | HPA039227 | 1: 1000 |
| Anti-OSBP2 antibody produced in rabbit | Sigmaaldrich | HPA021514 | 1: 1000 |
| Beta Actin Monoclonal Antibody | Invitrogen | AM4302 | 1: 5000 |
| Goat anti-Rabbit IgG (H+L) Secondary Antibody, HRP | Invitrogen | 31460 | 1: 10 000 |
| F(ab')<sub>2</sub>-Goat anti-Mouse IgG (H+L) Secondary Antibody, HRP | Invitrogen | A24512 | 1: 10 000 |
| IRDye® 800CW Goat anti-Mouse IgG (H + L) | LI-COR | 925-32210 | 1: 15 000 |
| IRDye® 680RD Goat anti-Rabbit IgG (H + L) | LI-COR | 925-68071 | 1: 15 000 |

#### Immunoblotting analysis

For degradation tests, 2×10^5^ cells/mL A549 and HeLa cells in 2 mL medium were seeded in six-well plates and incubated (37 °C and 5% CO_2_) overnight. The media was then removed and replaced with the fresh media containing the required concentration of compounds dissolved in DMSO. At the specific time point, cells were washed with ice-cold PBS twice and then lysed in lysis buffer (100 mM Tris–HCl (pH 6.8), 4% (w/v) sodium dodecyl sulfate (SDS), 20% (v/v) glycerol) and sonicated by ultrasonic processor (UP100H, Hielscher). The protein concentrations of the lysate were determined using NanoDrop ND-1000 spectrophotometer (Saveen Werner AB, Limhamn, Sweden) according to the manufacturer’s instructions and further diluted to 4 mg/mL. SDS–PAGE was carried out using 4-15% or 4-20% precast polyacrylamide gels (Bio-Rad, 4561086 or 4561093) and run at a constant voltage of 80 V for 5 min followed by 120 V for 1 h. Semi-dry transfer onto a PVDF membrane was performed using Bio-Rad Trans-Blot Turbo Transfer System at 1.3 A for 7 min. For Chemiluminescent detection, membranes were blocked in 5% milk in TBST (137 mM NaCl, 19 mM Tris-base, 2.7 mM KCl and 0.1% (v/v) Tween-20, blocking buffer) for 1 h at room temperature. The membrane was incubated with the primary antibody in blocking buffer overnight at 4 °C. After washing with TBST (3×5 min), the membrane was incubated with the secondary antibody in blocking buffer for 1 h at room temperature. Signals were visualized using the SuperSignal West Pico Chemiluminescent Substrate (Thermo Fisher, 34579) or the SuperSignal West Femto Maximum Sensitivity Substrate (Thermo Fisher, 34094) on a Li-COR Odyssey Fc. For fluorescent detection, membranes were blocked in Intercept® (TBS) Blocking Buffer (Li-COR, 927-60001) for 1 h at room temperature. The membrane was incubated with the primary antibody in blocking buffer overnight at 4 °C. After washing with TBST (3×5 min), the membrane was incubated with the secondary antibody in blocking buffer for 1 h at room temperature. Signals were visualized on a Li-COR Odyssey Fc.

### In-Cell Cellular Thermal Shift Assay (CETSA)

Two T75 cell culture flasks were seeded with each 6×10^5^ HeLa cells in media and incubated at 37°C and 5% CO_2_ for three days. The media was then removed and replaced with 3 mL medium containing either 10 µM of the given compound or DMSO. After incubation for 1 h at 37°C and 5% CO_2_ the medium was removed and the cells were washed by PBS and detached using 1.5 mL trypsin. And then the media and trypsin were removed by centrifuge at 250 × g for 3 min. After three times washing by PBS, 1×10^6^ cells were collected in 0.6 mL PBS each. Treated and non-treated cell suspensions were divided into ten aliquots, each 100 µL in PCR tubes. The aliquots were individually heated at different temperatures (Doppio 2×48 well Thermal cycler, VWR). After the heat treatment, 10 µL PBS containing 4.4% (v/v) NP-40 added to each sample and cells were lysed by freeze and thaw four times. The cell lysates were completely transferred to polycarbonate tubes and centrifuged (Micro Star 30, VWR) at 25,000 × g, 4°C for 25 min. The cleared supernatants were collected and further analyzed by western blotting assay.

### In-Cell Isothermal dose-response fingerprinting (ITDRF) experiments

Two six well plates were seeded with each 3×10^5^ HeLa cells in 2 mL DMEM per well and incubated at 37°C and 5% CO_2_ overnight. The media was subsequently removed and replaced with 0.5 mL medium containing either different concentrations (40, 20, 10, 5, 2.5, 1.25, 0.625, 0.3 µM) of the test compound or DMSO. After incubation for 1 h at 37°C and 5% CO_2_, the medium was removed and cells were detached using 200 mL trypsin and collected in 0.5 mL MEM each. Treated and non-treated cell suspensions were centrifuged (350 × g, 5 min), and resuspended in 50 µL PBS in PCR tubes. The samples were heated at the given temperature (Eppendorf Mastercycler ep Gradient S). After the heat treatment, 5 µL PBS containing 4.4% (v/v) NP-40 were added to each sample and cells lysed by freeze and thaw four times and then the cell lysates were completely transferred to polycarbonate tubes and centrifuged (Micro Star 30, VMR) at 25,000 × g, 4 °C for 25 min. The cleared supernatants were collected and further analyzed by western blotting assay.

### Cell lysate Cellular Thermal Shift Assay (CETSA)

HeLa cell lysates (protein concentration 2 mg/mL) were separated in the two aliquots (1.0 mL) to be incubated for 10 min at room temperature with either 0.1% DMSO (1.0 µL) or with 10 µM of the tested compound. After incubation, each aliquot (each 100 µL) was heated at different temperatures for 3 min using PCR gradient-cyclers.

After heating, the removal of the precipitated proteins from the solution was performed by ultracentrifugation at 4 °C with a speed of 25000 × g for 25 min. The cleared supernatants were collected and further analyzed by western blotting.

### Cell lysate Isothermal dose-response fingerprinting (ITDRF) experiments

HeLa lysates (protein concentration 2 mg/mL) were separated in 9 aliquots (each 100 µL) to be incubated for 10 min at room temperature with either 0.1% DMSO or with decreasing concentration of the tested compounds (40, 20, 10, 5, 2.5, 1.25, 0.625, 0.3 µM). All the aliquots were successively heated for 3 min at the given temperature using PCR gradient-cyclers.

After heating, the removal of the precipitated proteins from the solution was performed by ultracentrifugation at 4 °C with a speed of 25000 × g for 25 min. The cleared supernatants were collected and further analyzed by western blotting.

### Deglycosylation assay

HeLa cell lysates were incubated with PNGase F (New England Biolabs, NEB-P0704S) according to the manufacturer’s instructions. Briefly, 10 μL of reaction mixture (9 μL of 1 mg/mL HeLa cell lysate and 1 μL of Glycoprotein Denaturing Buffer (10×)) was denatured by heating at 100 °C for 10 min. The denatured protein mixture was chilled on ice and centrifuged for 10 s. Then, 2 μL of GlycoBuffer 2 (10×), 2 μL of 10% NP - 40, 6 μL of H_2_O were added to the previous reaction mixture to make a total reaction volume of 20 μL. Next, 1 μL PNGase F was added to the reaction mixture and the mixture was incubated at 37 °C for 2 h and the analyzed by western blotting as previously described.

### Protein expression

#### Protein expression constructs

Human ASTER domains of Aster-A(359-547), -B(364-552) and -C(318-504) were subcloned into a pGEX-6p-2rbs vector, thus introducing the cloning artifact ‘GPLGS ’. The pGEX-6P-1-GST-OSBP(377-807) and pET28(+)-ORP2(49-480) plasmids were purchased from Genscript. The pET22b_His6_STARD1(66-284) plasmid was a gift from James H. Hurley (University of California ).

#### Protein expression and purification

ASTER domains of human Aster-A(359-547), -B(364-552) and -C(318-504) in the pGEX-6p-2rps vector with an N-terminal PreScission-cleavable GST-tag were expressed in Escherichia coli OverExpress C41 in Terrific Broth (TB) medium for approximately 16 h at 18 °C after induction with 0.1 mM IPTG. Cells were collected at 3,500g for 15 min and lysed by sonication in buffer containing 50 mM HEPES pH 7.5, 300 mM NaCl, 10% (vol/vol) glycerol, 5 mM DTT, 0.1% (vol/vol) Triton X-100 and protease inhibitor mix HP plus (Serva). The cleared lysate was purified by affinity chromatography on a GSTrap FF column (Cytiva) using an ÄKTA Start (Cytiva) in buffer containing 50 mM HEPES pH 7.5, 300 mM NaCl, 10% (vol/vol) glycerol, 5 mM DTT and 0.01% (vol/vol) Triton X-100. The GST-tag was cleaved on the column overnight at 4 °C. Proteins were further purified by size-exclusion chromatography on a HiLoad 16/600 Superdex 75 pg (Cytiva) in buffer containing 20 mM HEPES pH 7.5, 300 mM NaCl, 10% (vol/vol) glycerol and 2 mM DTT.

The START domain of human STARD1(66-284) harboring an N-terminal His6-Tag was expressed in Escherichia coli BL21(DE3) in Luria-Bertani Broth (LB) medium for approximately 16 h at 18 °C after induction with 0.15 mM IPTG. Cells were collected at 3,500g for 15 min and lysed by sonication in buffer containing 50 mM HEPES pH 7.5, 150 mM NaCl, 5% (vol/vol) glycerol, 5 mM DTT, 0.1% (vol/vol) Triton X-100 and EDTA-free protease inhibitor cocktail (Sigma-Aldrich). The cleared lysate was purified by affinity chromatography on a Ni-NTA Superflow Cartridge (Qiagen) using an ÄKTA Start (Cytiva) in buffer containing 50 mM HEPES pH 7.5, 150 mM NaCl, 5 % (vol/vol) glycerol, 5 mM DTT. START domains were eluted by using elution buffer containing 50 mM HEPES pH 7.5, 150 mM NaCl, 5% (vol/vol) glycerol, 5 mM DTT and 500 mM imidazole. Proteins were further purified by size-exclusion chromatography on a HiLoad 16/600 Superdex 75 pg (Cytiva) in buffer containing 20 mM HEPES pH 7.5, 150 mM NaCl, 5% (vol/vol) glycerol and 2 mM DTT.

The ORP domain of human OSBP(377-807) in the pGEX-6p-1 vector with an N-terminal PreScission-cleavable GST-tag was expressed in Escherichia coli OverExpress C41 in LB medium for approximately 16 h at 18 °C after induction with 0.1 mM IPTG. Cells were collected at 3,500g for 15 min and lysed by sonication in buffer containing 20 mM HEPES pH 7.5, 300 mM NaCl, 10% (vol/vol) glycerol, 5 mM DTT, 0.1% (vol/vol) Triton X-100 and EDTA-free protease inhibitor cocktail (Sigma-Aldrich). The cleared lysate was purified by affinity chromatography on a GSTrap HF column (Cytiva) using an ÄKTA Start (Cytiva) in buffer containing 20 mM HEPES pH 7.5, 300 mM NaCl, 10% (vol/vol) glycerol, 5 mM DTT. OSBP(377-807) was eluted by using elution buffer containing 20 mM HEPES pH 7.5, 300 mM NaCl, 10% (vol/vol) glycerol, 5 mM DTT and 10 mM reduced glutathione. Proteins were further purified by size-exclusion chromatography on a HiLoad 16/600 Superdex 75 pg (Cytiva) using an ÄKTA Explorer (Cytiva) in buffer containing 20 mM HEPES pH 7.5, 150 mM NaCl, 10% (vol/vol) glycerol and 2 mM DTT.

The ORP domain of human ORP2(49-480) in the pET24b(+) vector including an N-terminal His6-Tag was expressed in Escherichia Coli BL21(DE3) in Terrific Borth medium for approximately 16 h at 18 °C after induction with 0.1 mM IPTG. Cells were collected at 3,500 g for 15 min and lysed by sonification in buffer containing 10 mM Tris-HCl pH 8, 300 mM NaCl, 5 % glycerol, 2 mM DTT, 0.1 % Triton X-100 and EDTA-free protease inhibitor cocktail (Sigma-Aldrich). The cleared lysate was purified by affinity chromatography on a Ni-NTA Superflow Cartridge (Qiagen) using an ÄKTA Start (Cytiva) in buffer containing 10 mM Tris-HCl, pH 8, 300 mM NaCl, 5 % glycerol, 2 mM DTT. ORP2(49-480) was eluted using elution buffer containing 10 mM Tris-HCl, pH 8, 300 mM NaCl, 5 % glycerol, 500 mM imidazole, 2 mM DTT. The protein was further purified by size-exclusion chromatography on a HiLoad 16/600 Superdex 75 pg (Cytiva) using an ÄKTA Explorer (Cytiva) in buffer containing 10 mM Tris-HCl, pH 8, 150 mM NaCl, 5 % glycerol, 2 mM DTT.

### Fluorescence polarization

Fluorescence polarization experiments were performed in a buffer composed of 20 mM HEPES pH 7.5, 300 mM NaCl, 0.01% (vol/vol) Tween-20, 0.5% glycerol and 2 mM DTT in a final volume of 30 µl in black, flat-bottom, non-binding 384-well plates (Corning). For competition experiments, 20 nM 22-NBD-cholesterol was mixed with protein and incubated with desired concentrations of screening compounds. The fluorescence polarization signal was measured using a Spark Cyto multimode microplate reader (Tecan) with filters set at 485 ± 20 nm for excitation and at 535 ± 20 nm for emission. The data was analysed using GraphPad Prism 5. Measured mP values were normalized setting 100% inhibition as the FP signal from the protein + fluorophore control well and 0% as the FP signal from the fluorophore control well. Curves were fitted to the normalized data via non-linear regression to allow the determination of IC50 values.

| Protein | Protein conc. | Incubation temp. | Incubation time<br>(Protein/22-NBD cholesterol) | Incubation time<br>(Protein-22-NBD-cholesterol/compound) |
| --- | --- | --- | --- | --- |
| Aster-A | 1 μM | room temperature | 20 min | 20 min |
| Aster-B | 1 μM | room temperature | 20 min | 20 min |
| Aster-C | 0.5 μM | room temperature | 20 min | 20 min |
| STARD1 | 2 μM | 4 °C | 30 min | 20 min |
| OSBP | 0.3 μM | 4 °C | 60 min | 20 min |
| ORP2 | 2 | 4 °C | 60 min | 20 min |

### HiBiT-Cellular thermal shift assay (CETSA) in intact cells

After treatment with test compounds or DMSO (1 million cells/mL), KBM7 cells overexpressing HiBiT-tagged ORPs were spun down (1450 rpm, 5 min), washed once with PBS, and re-suspended in PBS (100 μL of PBS per 1 million of cells). The cell suspension (100 μL) was distributed into PCR strips, spun down (1450 rpm, 5 min) and 80 μL of the supernatant were removed without disturbing the cell pellet. The samples were then thermally destabilized (3-min temperature gradient of choice followed by 3 min at 25 °C) and re-suspended in 30 uL of the lysis buffer (20 mM Tris-HCl pH 8.0, 120 mM NaCl, 0.5 % NP-40). Three freeze/thaw cycles with liquid nitrogen were then performed and the lysed samples were cleared out via centrifugation (full speed, 20 min, 4 °C).

Luminescence was assessed with the Nano-Glo^®^ HiBiT Lytic Detection System (Promega, N3040) in 384-well plate format and in technical triplicates. Briefly, a 6x detection master mix was prepared containing the substrate, LgBiT and the provided lysis buffer and 2.5 μL were distributed into well plates. Cleared lysates of cells expressing HiBiT-tagged ORPs (12.5 μL) were then added into respective wells and the plates were incubated at room temperature for 10 mins with mild shaking. Luminescence signal was measured on the Multilabel Plate Reader Platform Victor X3 model 2030 (Perkin Elmer). Melting curves and *T*_m_ were determined in GraphPad Prism (v9.5.1) by fitting a non-linear regression curve or interpolation of a sigmoidal standard curve. Each data point was normalized to the mean luminescence at the lowest temperature (typically 37.6 °C).

### Shiga toxin trafficking assay

HeLa cells were seeded one day before use, and used at 70% confluency. Cells were treated for 4 h with 1 μM of each compound or DMSO vehicle, in complete media at 37°C. Media was changed to fresh media containing 0.5 μg/mL STxB-488 (obtained from Ludger Johannes, Institut Curie) and compounds as before. HeLa cells were incubated at 37°C for 40 min, then fixed in 4% PFA for 10 min at RT. Immunofluorescence was performed.

### Golgi Fragmentation assay

HeLa cells were used at 70% confluency. Cells were treated with 1 μM compounds or DMSO vehicle for 4 h at 37°C, then fixed with 4% PFA for 10 min at RT, followed by immunofluorescence analysis.

### Immunofluorescence

Fixed cells were blocked and permeabilized for 20 min in blocking buffer (0.05% (w/v) saponin, 0.5% (w/v) BSA, 50 mM NH_4_Cl and 0.02% NaN_3_ in PBS, pH 7.4). Cells were incubated for 1 h with primary antibodies against GPP130 (a gift from A. Linstedt), TGN46 (Biorad, AHP500GT, 1:200), washed 3 times with PBS and incubated in blocking buffer for 45 min with fluorescently labelled secondary antibodies at 1:400, purchased from Beckman Coulter (donkey anti-mouse-FITC no. 715196150, lot 92566; donkey anti-sheep-Cy3 no. 713166147, lot 106361). Nuclei were labelled for 5 min with 5 μg/mL 4’6-diamidino-2-phenylindole (DAPI, Vector Laboratories), and mounted using Mowiol (Sigma-Aldrich).

### Image acquisition

Images were acquired on an inverted Eclipse Ti-E (Nikon) microscope equipped with a spinning disk CSU-X1 (Yokogawa) module, a x60 CFI Plan Apo VC oil objective and Metamorph software by Gataca Systems. *Z*-stacks were acquired in a step size of 0.4 μm. The pixel size was 0.267 μm.

### Image analysis

#### STxB trafficking

STxB trafficking was previously characterised^59^ by quantification of STxB arrival at Golgi (Giantin), or localisation to early endosomes (EEA1). Based on this, STxB spots were counted in one central Z-slice per cell, correlating low numbers to STxB arrival at the Golgi, and high numbers correlating to the cytosolic spots that represent colocalization to early endosomes. Fluorescence area was measured in parallel and remained unchanged in all samples. Data is presented as number of STxB spots normalised to area of fluorescence.

#### Semi-automated quantification

Fiji ImageJ2 software (National Institute of Health)^65^ was used for image processing and quantification. Quantification was semi-automated by applying Max Entropy automatic threshold to individual cells to determine areas of positive fluorescence (for STxB or TGN46), then counting the number of fragments above threshold per cell by using the *analyze particles* command.

#### Statistics

Statistics were performed in GraphPad PRISM9 software. One way ANOVA was applied with Dunnett’s multiple comparisons test to compare each treatment group to CTRL. A P-value of 0.05 or less was considered significant.

### Protein synthesis assays in cells exposed to Shiga toxins^66^

HeLa cells were grown at 37 °C in an atmosphere of 5% CO_2_ and 95% air on 150 cm^2^ tissue culture flasks in Dulbecco’s Modified Eagle’s Medium (Invitrogen) containing 10% SVF, 2 mM L-glutamine, 0.1 mM non-essential amino acids, 100 U/mL penicillin and 100 µg/mL streptomycin.

First, the cells were plated at a density of 10,000 cells per well in 96-well Cytostar T scintillating microplates with scintillator incorporated into the polystyrene plastic (100 µL/well). Then, complete DMEM medium supplemented with compound was added (50 µL/well). After a short incubation time, complete DMEM medium supplemented with Stx-1, (List Lab) was added to each well of HeLa-seeded scintillating microplates (dilutions of Stx-1 ranging from 10^-9^ M to 10^-16^ M; 50 µL/well). After incubation for 20 hours, the media (200 µL/well) were removed and replaced with DMEM without leucine containing 10% SVF, 2 mM L-glutamine, 0.1 mM non-essential amino acids, 1% Penicillin/Streptomycin supplemented by 0.5 µCi/mL [^14^C]-leucine. The cells were grown for an additional 5 hours at 37°C in an atmosphere of 5% CO_2_ and 95% air. Protein biosynthesis was determined by measuring the incorporation of radiolabeled leucine using a Wallac MicroBeta Trilux scintillation counter.

The mean percentage of protein biosynthesis was determined and normalized from duplicate wells. All values are expressed as means ± SD. Data were fitted with Prism v5 software (Graphpad Inc., San Diego, Calif.) to obtain the 50% effective toxin concentration (EC_50_). Plotting (1 - EC_50_)*100/(1 - EC_50max_) as a function of compound concentration enables the calculation of the IC_50_, which represents the concentration of compound giving 50 % of its inhibitory effect. The goodness of fit for each toxin and drug was assessed by R^2^ and confidence intervals.

