## Supporting Information for "Selective inhibition of OSBP blocks retrograde trafficking by inducing partial Golgi degradation"

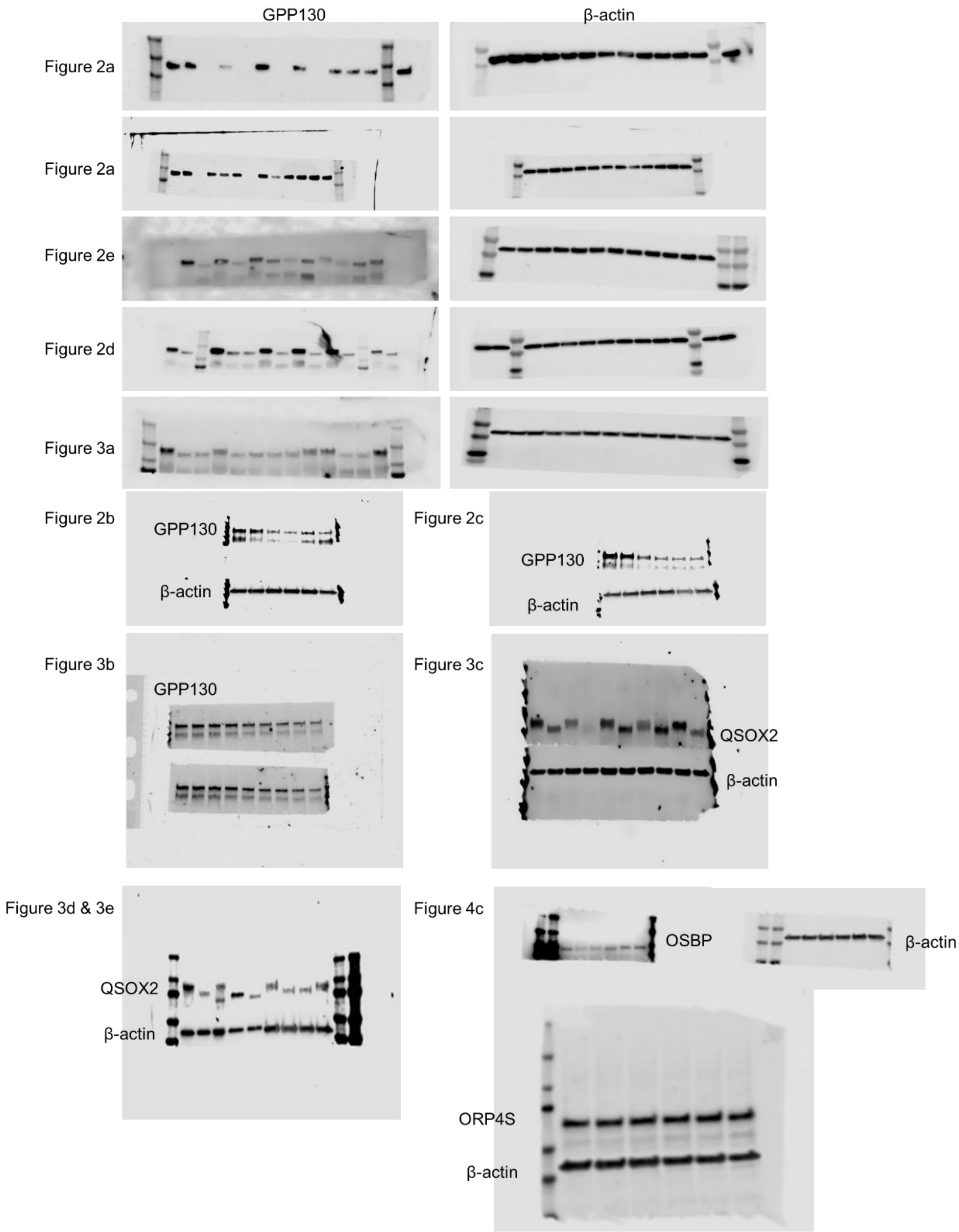


Supplementary Figure 1. Uncropped blots from Figures 2a-2e, 3a, 3c-e and 4c.

Extended Data Fig. 4b

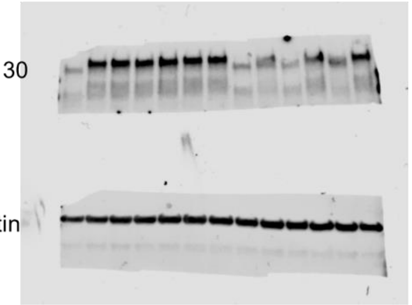


Extended Data Fig. 5a

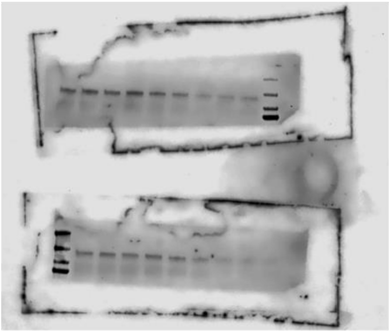

Extended Data Fig. 5b

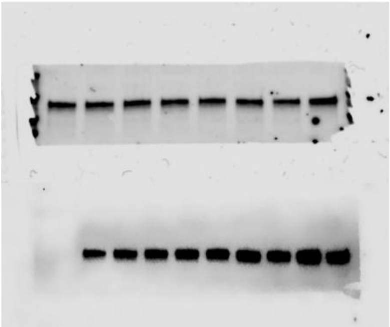


Extended Data Fig. 6

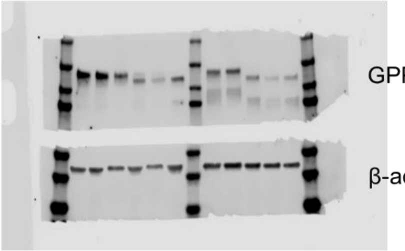


Supplementary Figure 2. Uncropped blots from Extended Data Fig. 4b, 5, and 6.

#### Synthesis

General directions

Commercially available reagents were used without further purification and all solvents were of HPLC quality. All reactions were monitored by thin layer chromatography (TLC) and/or reversed-phase ultra-performance liquid chromatography mass spectrometry (RP-UPLC-MS)

Analytical TLC was conducted on Merck aluminium sheets covered with silica (C60). The plates were either visualized under UV-light or stained by dipping in a developing agent followed by heating. $KMnO_4$ [3 g in water (300 mL) along with $K_2CO_3$ (20 g) and 5% aqueous NaOH (5 mL)] was used as developing agents. Flash column chromatography was performed using Merck Geduran® Si60 (40-63 μm) silicagel.

All new compounds were characterized by $^1$H NMR, $^{13}$C NMR, MS (ESI), HRMS (ESI) and optical rotation (byproducts and intermediates were not fully characterized). For the recording of $^1$H NMR and $^{13}$C NMR a Bruker Ascend with a Prodigy cryoprobe (operating at 400 MHz for proton and 100 MHz for carbon) was used. The chemical shifts (δ) are reported in parts per million (ppm) and the coupling constants (J) in Hz. For spectra recorded in DMSO, signal positions were measured relative to the signal for DMSO (δ 2.50 ppm for $^1$H NMR and δ 39.43 ppm for $^{13}$C NMR). For spectra recorded in $CDCl_3$, signal positions were measured relative to the signal for $CHCl_3$ (δ 7.26 ppm for $^1$H NMR and δ 77.0 ppm for $^{13}$C NMR).

Analytical RP-UPLC-MS (ESI) analysis was performed on a S2 Waters AQUITY RP-UPLC system equipped with a diode array detector using an Thermo Accucore C18 column (d 2.6 μm, 2.1 x 50 mm; column temp: 50 °C; flow: 1.0 mL/min). Eluents A (0.1% $HCO_2H$ in $H_2O$) and B (0.1% $HCO_2H$ in MeCN) were used in a linear gradient (5% B to 100% B) in 2.4 min and then held for 0.1 min at 100% B (total run time: 2.6 min). The LC system was coupled to a SQD mass spectrometer.

Ultra-high Performance Liquid Chromatography-High Resolution Mass Spectrometry (UHPLC-HRMS) was performed on an Agilent Infinity 1290 UHPLC system (Agilent Technologies, Santa Clara, CA, USA) equipped with a diode array detector. Separation was obtained on an Agilent Poroshell 120 phenyl-hexyl column (2.1 × 150 mm, 1.9 μm) with a linear gradient consisting of water (A) and acetonitrile (B) both buffered with 20 mM formic acid, starting at 10% B and increased to 100% in 10 min where it was held for 2 min, returned to 10% in 0.1 min and remaining for 2 min (0.35 mL/min, 60 °C). An injection volume of 1 μL was used. MS detection was performed in both positive and negative detection on an Agilent 6545 QTOF MS equipped with Agilent Dual Jet Stream electrospray ion source with a drying gas temperature of 250 °C, gas flow of 8 L/min, sheath gas temperature of 300 °C and flow of 12 L/min. Capillary voltage was set to 4000 V and nozzle voltage to 500 V. Mass spectra were recorded at 10, 20 and 40 eV as centroid data for *m/z* 85–1700 in MS mode and *m/z* 30–1700 in MS/MS mode, with an acquisition rate of 10 spectra/s. Lock mass solution in 70:30 methanol:water was infused in the second sprayer using an extra LC pump at a flow of 15 μL/min using a 1:100 splitter. The solution contained 1 μM tributylamine (Sigma-Aldrich) and 10 μM Hexakis(2,2,3,3-tetrafluoropropoxy)phosphazene (Apollo Scientific Ltd., Cheshire, UK) as lock masses. The $[M + H]^+$ ions (*m/z* 186.2216 and 922.0098 respectively) of both compounds was used.

**Hyodeoxycholic acid methyl ester (1)**

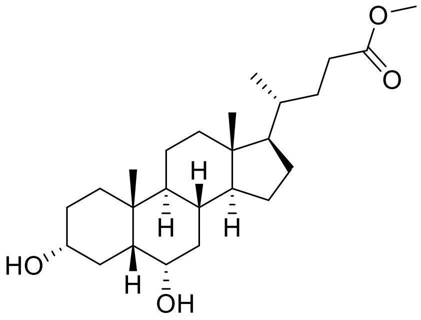

Hyodeoxycholic acid (20 g, 51 mmol) was dissolved in 100 mL methanol and then concentrated $H_2SO_4$ (2.5 mL) was added dropwise at 0 °C under $N_2$ atmosphere. The solution was stirred overnight at room temperature. Then, the solvent was removed under vacuum and the crude was dissolved in 40 mL EtOAc, washed by saturated $NaHCO_3$ (2 × 25 mL) and brine (2 × 25 mL). The organic phase was collected, dried over anhydrous sodium sulfate and evaporated under reduced pressure to obtain compound **1**.

White solid (20 g, 98%).

**$^1$H NMR** (400 MHz, $CDCl_3$) δ 4.04 (m, 1H), 3.66 (s, 3H), 3.64 – 3.55 (m, 1H), 2.36 (m, 1H), 2.22 (m, 1H), 2.02 – 1.30 (m, 17H), 1.23 – 0.98 (m, 7H), 0.91 (d, *J* = 7.1 Hz, 6H), 0.64 (s, 3H).

**$^{13}$C NMR** (101 MHz, $CDCl_3$): δ 174.89, 71.80, 68.32, 56.26, 56.05, 51.65, 48.48, 42.98, 40.07, 39.94, 36.10, 35.68, 35.48, 35.09, 34.97, 31.19, 31.09, 30.31, 29.28, 28.25, 24.33, 23.61, 20.88, 18.39, 12.16.

**MS** *m/z* (ES+) calcd for $C_{25}H_{42}O_4$, $[2M+H]^+$ 813.6, found: 813.5.

The NMR data matched those reported in the literature[1].

**Methyl 3α,6α-ditosyloxy-5β-cholan-24-oate (2)**

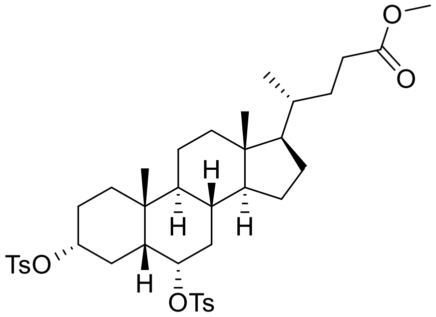

**1** (5 g, 12.5 mmol) was dissolved in dry pyridine (30 mL) and then a solution of TsCl (7 g, 37.5 mmol) dissolved in 20 mL dry pyridine was added dropwise. The solution was stirred at room temperature for 2 days. Then, 200 mL 10% HCl was poured into the solution and ice chips were added gradually. The precipitate was filtered off and washed with water to give compound **2**.

White solid (8.5 g, 95%).

**$^1$H NMR** (400 MHz, $CDCl_3$) δ 7.83 – 7.72 (m, 4H), 7.36 (t, *J* = 8.0 Hz, 4H), 4.81 (m, 1H), 4.32 (m, 1H), 3.68 (s, 3H), 2.48 (s, 6H), 2.36 (m, 1H), 2.22 (m, 1H), 1.98 – 0.93 (m, 24H), 0.90 (d, *J* = 6.3 Hz, 3H), 0.82 (s, 3H), 0.61 (s, 3H).

**$^{13}$C-NMR** (101 MHz, $CDCl_3$): δ 174.81, 144.84, 134.64, 134.63, 129.98, 129.94, 127.75, 127.66, 127.14, 81.92, 79.82, 55.95, 55.90, 51.65, 46.47, 42.96, 39.72, 39.58, 36.29, 35.39, 34.96, 32.22, 31.14, 31.01, 28.10, 27.53, 26.60, 24.06, 23.02, 21.82, 20.65, 18.34, 12.09.

The NMR data matched those reported in the literature[1].

**Methyl 3β-hydroxy-5-cholen-24-oate (3)**

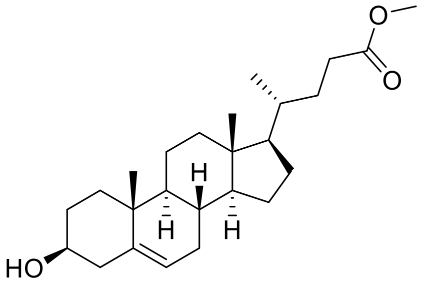


**3** (1 g, 1.4 mmol) and KOAc (100 mg, 1 mmol) were dissolved in 10 mL DMF and 1mL $H_2O$. The solution was refluxed at 105 °C overnight. And then, the solution was cooled down to room temperature, extracted by EtOAc (3 × 20 mL) and washed by brine (3 × 20 mL). The organic phase was collected, dried over anhydrous sodium sulfate and evaporated under reduced pressure. The residue was purified by column chromatography (gradient of pentane/EtOAC 8:1 to 2:1).

White solid (282 mg, 52%).

**$^{1}$H NMR** (400 MHz, $CDCl_3$) δ 5.35 (m, 1H), 3.66 (s, 3H), 3.57 – 3.46 (m, 1H), 2.41 – 2.16 (m, 4H), 2.05 – 1.74 (m, 8H), 1.65 – 1.23 (m, 9H), 1.20 – 1.02 (m, 4H), 1.00 (s, 3H), 0.92 (d, $J$ = 6.5 Hz, 3H), 0.68 (s, 3H).

**$^{13}$C-NMR** (101 MHz, $CDCl_3$): δ 174.93, 140.88, 121.82, 71.94, 56.87, 55.91, 51.64, 50.22, 42.51, 42.43, 39.88, 37.39, 36.64, 35.52, 32.03, 32.01, 31.79, 31.21, 31.16, 28.26, 24.40, 21.21, 19.54, 18.46, 12.01.

**MS** *m/z* (ES+) calcd for $C_{25}H_{40}O_3$ $[M+H]^+$ 389.3, found: 389.1.

The NMR data matched those reported in the literature[2].

**3β-hydroxy-5-cholenoic acid (4)**

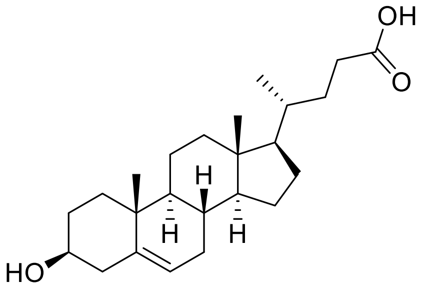


3 (500 mg, 1.28 mmol) was dissolved in 10 mL 4% KOH/MeOH and the solution was stirred overnight at room temperature. Then, 1 M HCl was added gradually and the precipitate was filtered off and washed with $H_2O$ to obtain compound 4.

White solid (344 mg, 72%).

**$^1$H NMR** (400 MHz, $CDCl_3$) δ 5.37 (m, 1H), 3.55 (m, 1H), 2.38 – 2.28 (m, 1H), 2.25 – 2.14 (m, 3H), 2.08 – 1.75 (m, 6H), 1.70 – 1.04 (m, 15H), 1.03 (s, 3H), 0.96 (d, *J* = 6.6 Hz, 3H), 0.73 (s, 3H).

**$^{13}$C-NMR** (101 MHz, $CDCl_3$): δ 178.17, 142.23, 122.43, 72.44, 58.14, 57.30, 51.70, 43.55, 43.02, 41.12, 38.55, 37.68, 36.71, 33.26, 33.01, 32.34, 32.30, 32.01, 29.14, 25.30, 22.19, 19.86, 18.79, 12.31.

The NMR data matched those reported in the literature[3].

**2-(2,6-dioxopiperidin-3-yl)-4-fluoroisoindoline-1,3-dione (5)**

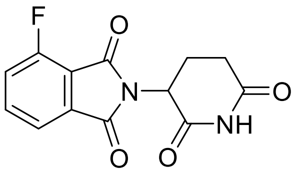


3-Fluorophthalic anhydride (996 mg, 6 mmol), 3-aminopiperidine-2,6-dione hydrogen chloride (658 mg, 4 mmol) and NaOAc (558 mg, 6.8 mmol) were added to HOAc (20 mL) and the solution was heated at 135 °C for 8 h. Then, the reaction was cooled down to room temperature and the residue was suspended in 100 mL ice water. The solid was collected by vacuum filtration and used without further purification.

Grey solid (512 mg, 47%).

The NMR data matched those reported in the literature[4].

**$^1$H NMR** (400 MHz, DMSO-$d_6$) δ 11.16 (s, 1H), 7.96 (m, 1H), 7.80 (d, *J* = 7.3 Hz, 1H), 7.75 (t, *J* = 8.9 Hz, 1H), 5.17 (m, 1H), 2.90 (m, 1H), 2.69 – 2.53 (m, 2H), 2.07 (m, 1H).

**MS** *m/z* (ES+) calcd for $C_{13}H_9FN_2O_4$ $[M+H]^+$ 277.1, found 276.9.

**4-fluoro-2-(1-methyl-2,6-dioxopiperidin-3-yl)isoindoline-1,3-dione (6)**

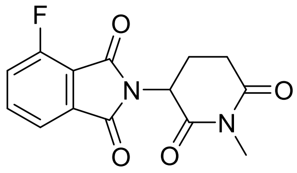

**5** (100 mg, 0.36 mmol) and NaH (21 mg, 0.54 mmol) were dissolved in 6 mL DMF and stirred for 15 min under 0 °C. Then, MeI (76.6 mg, 0.54 mmol) was added to the mixture dropwise under 0 °C and the reaction mixture was stirred at room temperature for 8 h. Then, the solution was diluted with 30 mL water and extracted by EtOAc (3 × 10 mL), washed with brine (3 × 20 mL), dried over anhydrous $Na_2SO_4$ and concentrated to obtain **6** as a green oil without further purification.

The data matched those reported in the literature[5].

**MS** *m/z* (ES+) calcd for $C_{14}H_{11}FN_2O_4$ $[M+H]^+$ 290.3, found 290.9.

**General procedure I: Mono-Boc protection of diamines.**

A solution of Di-*tert*-butyl dicarbonate (1 equiv.) in $CH_2Cl_2$ was added dropwise to a solution of the corresponding diamine (6 equiv.) in $CH_2Cl_2$ at 0 °C. The solution was stirred at 0 °C for 3 h and then at room temperature overnight. And then the mixture was washed by $H_2O$ and brine. The organic phase was collected, dried over $Na_2SO_4$ and evaporated under reduced pressure.

The data for the following compounds matched those reported in the literature[6].

***tert*-butyl (6-aminohexyl)carbamate (7)**

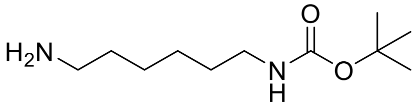

The compound was prepared using General procedure I to give a colorless oil. Yield 1.9 g, 88%.

**$^1$H NMR** (400 MHz, $CDCl_3$) δ 4.59 (s, 1H), 3.11 (m, 2H), 2.71 (t, *J* = 7.0 Hz, 2H), 2.10 (s, 2H), 1.52 – 1.46 (m, 4H), 1.44 (s, 9H), 1.34 (m, 4H).

***tert*-butyl (2-(2-(2-aminoethoxy)ethoxy)ethyl)carbamate (8)**

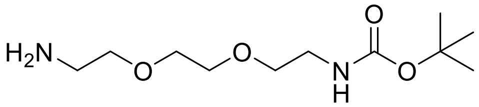

The compound was prepared using General procedure I to give a colorless oil. Yield 2.0 g, 81%.

**$^1$H NMR** (400 MHz, $CDCl_3$) δ 5.21 (s, 1H), 3.59 (s, 4H), 3.51 (m, 4H), 3.29 (m, 2H), 2.86 (t, *J* = 5.2 Hz, 2H), 1.41 (s, 9H).

***tert*-butyl (3-(4-(3-aminopropoxy)butoxy)propyl)carbamate (9)**

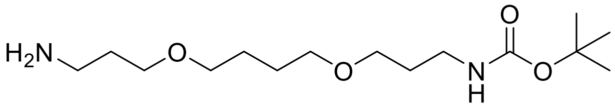

The compound was prepared using General procedure I to give a colorless oil. Yield 2.5 g, 82%.

**$^1$H NMR** (400 MHz, $CDCl_3$) δ 5.01 (s, 1H), 3.52 – 3.35 (m, 8H), 3.19 (m, 2H), 2.80 (t, *J* = 6.7 Hz, 2H), 1.77 – 1.67 (m, 4H), 1.65 – 1.56 (m, 4H), 1.41 (s, 9H).

***tert*-butyl (3-(2-(2-(3-aminopropoxy)ethoxy)ethoxy)propyl)carbamate (10)**

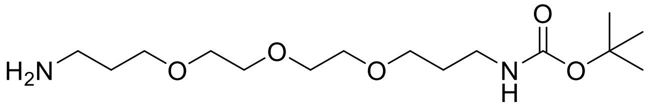

The compound was prepared using General procedure I to give a colorless oil. Yield 3.2 g, 75%.

**$^1$H NMR** (400 MHz, $CDCl_3$) δ 5.39 (t, *J* = 5.7 Hz, 1H), 3.42 – 3.25 (m, 12H), 2.94 (m, 2H), 2.53 (t, *J* = 6.7 Hz, 2H), 1.47 (m, 4H), 1.18 (s, 9H).

**General procedure II: Nucleophilic aromatic substitution ($S_NAr$).**

4-fluoro-pomalidomide (**5**, 1 equiv.) or 4-fluoro-2-(1-methyl-2,6-dioxopiperidin-3-yl)isoindoline-1,3-dione (**6**, 1 equiv.), DIPEA (4 equiv.) and the corresponding mono-protected diamine (1 equiv.) were dissolved in dry DMF. The mixture was stirred at 90 °C for 6 h. After cooling to room temperature, the mixture was diluted by $H_2O$ (20 mL) and then extracted by EtOAc (3 × 30 mL). The combined organic phase was further washed by brine (3 × 30 mL), dried over anhydrous sodium sulfate and evaporated under reduced pressure. The residue was purified by column chromatography (gradient of pentane/EtOAc 4:1 to 1:4).

The data for the following compounds matched those reported in the literature[6].

***tert*-butyl (6-((2-(2,6-dioxopiperidin-3-yl)-1,3-dioxoisoindolin-4-yl)amino)hexyl)carbamate (11)**

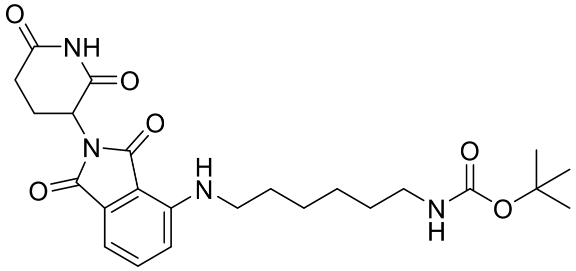

**5** (400 mg, 1.45 mmol) and **7** (313 mg, 1.45 mmol) were combined according to general procedure II. The product was a yellow solid. Yield 309 mg, 45%.

**$^1$H NMR** (400 MHz, $CDCl_3$) δ 7.99 (s, 1H), 7.54 – 7.49 (m, 1H), 7.12 (d, *J* = 7.1 Hz, 1H), 6.90 (d, *J* = 8.6 Hz, 1H), 4.97 – 4.90 (m, 1H), 4.53 (s, 1H), 3.29 (t, *J* = 7.0 Hz, 2H), 3.14 (s, 2H), 2.96 – 2.71 (m, 3H), 2.16 (m, 1H), 1.75 – 1.57 (m, 4H), 1.53 (m, 2H), 1.47 (s, 9H), 1.45 (m, 2H).

**MS** *m/z* (ES+) calcd for $C_{24}H_{32}N_4O_6$ [M+ $H^+$] 473.2, found 473.6.

***tert*-butyl (2-(2-(2-((2-(2,6-dioxopiperidin-3-yl)-1,3-dioxoisoindolin-4-yl)amino)ethoxy)ethoxy)ethyl)carbamate (12)**

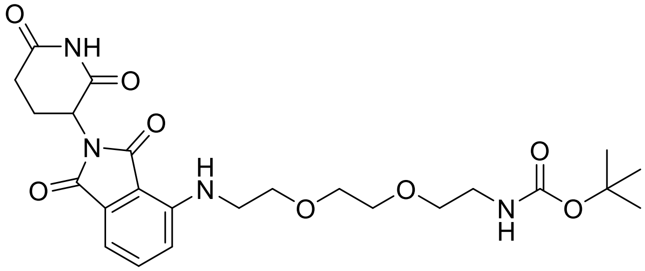

**5** (800 mg, 2.9 mmol) and **8** (720 mg, 2.9 mmol) were combined according to general procedure II. The product was a yellow oil. Yield 512 mg, 35%.

**$^1$H NMR** (400 MHz, $CDCl_3$) δ 8.48 (s, 1H), 7.51 (dd, *J* = 8.5, 7.1 Hz, 1H), 7.13 (d, *J* = 7.1 Hz, 1H), 6.93 (d, *J* = 8.5 Hz, 1H), 5.09 (s, 1H), 4.97 (s, 1H), 3.74 (t, *J* = 5.3 Hz, 2H), 3.67 (m, 4H), 3.58 (t, *J* = 5.2 Hz, 2H), 3.49 (t, *J* = 5.3 Hz, 2H), 3.33 (t, *J* = 5.2 Hz, 2H), 2.93 – 2.71 (m, 3H), 2.15 (m, 1H), 1.45 (s, 9H).

**MS** *m/z* (ES+) calcd for $C_{24}H_{32}N_4O_8$ [M+ $H^+$] 505.2, found 505.1.

***tert*-butyl (3-(4-(3-((2-(2,6-dioxopiperidin-3-yl)-1,3-dioxoisoindolin-4-yl)amino)propoxy)butoxy)propyl) carbamate (13)**

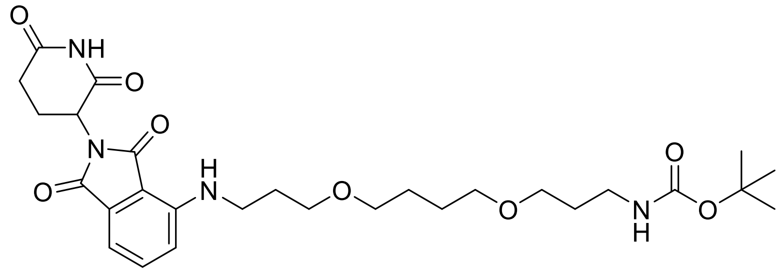

**5** (400 mg, 1.45 mmol) and **9** (441 mg, 1.45 mmol) were combined according to general procedure II. The product was a yellow oil. Yield 439 mg, 54%.

**$^1$H NMR** (400 MHz, $CDCl_3$) δ 8.25 (s, 1H), 7.51 (dd, *J* = 8.5, 7.1 Hz, 1H), 7.11 (d, *J* = 7.1 Hz, 1H), 6.94 (d, *J* = 8.5 Hz, 1H), 6.47 (s, 1H), 4.93 (m, 1H), 3.56 (t, *J* = 5.7 Hz, 2H), 3.52 – 3.39 (m, 8H), 3.23 (q, *J* = 6.2 Hz, 2H), 2.96 – 2.70 (m, 3H), 2.15 (m, 1H), 1.94 (p, *J* = 6.2 Hz, 2H), 1.76 (m, 2H), 1.72 – 1.61 (m, 4H), 1.46 (s, 9H).

**MS** *m/z* (ES+) calcd for $C_{28}H_{40}N_4O_8$ [M+ H$^+$] 561.3, found 561.2.

***tert*-butyl (3-(4-(3-((2-(1-methyl-2,6-dioxopiperidin-3-yl)-1,3-dioxoisoindolin-4-yl)amino)propoxy)butoxy) propyl)carbamate (14)**

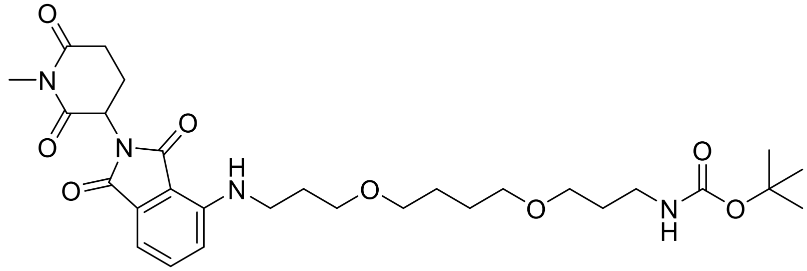

**6** (150 mg, 0.52 mmol) and **9** (157 mg, 0.52 mmol) were combined according to general procedure II. The product was a green oil. Yield 120 mg, 40%.

**$^1$H NMR** (400 MHz, $CDCl_3$) δ 7.50 (dd, *J* = 8.5, 7.1 Hz, 1H), 7.10 (dd, *J* = 7.2, 0.6 Hz, 1H), 6.94 (d, *J* = 8.6 Hz, 1H), 4.91 (m, 1H), 3.54 (m, 2H), 3.51 – 3.37 (m, 8H), 3.23 (s, 5H), 3.02 – 2.93 (m, 1H), 2.84 – 2.71 (m, 2H), 2.15 – 2.05 (m, 1H), 1.96 – 1.89 (m, 2H), 1.76 (m, 2H), 1.67 (m, 4H), 1.46 (s, 9H).

**MS** *m/z* (ES+) calcd for $C_{29}H_{42}N_4O_8$ [M+ H$^+$] 575.3, found 575.3.

***tert*-butyl (3-(2-(2-(3-((2-(2,6-dioxopiperidin-3-yl)-1,3-dioxoisoindolin-4-yl)amino)propoxy)ethoxy)ethoxy)propyl)carbamate (15)**

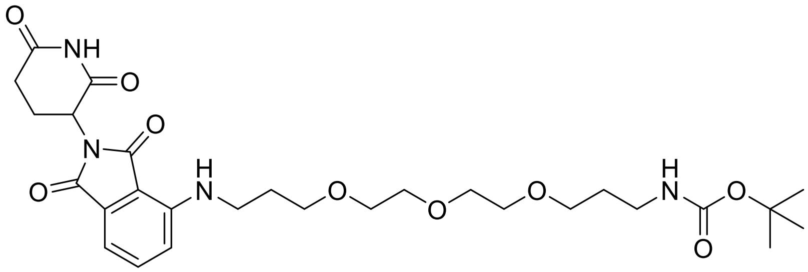

**5** (500 mg, 1.81 mmol) and **10** (580 mg, 1.81 mmol) were combined according to general procedure II. The product was a yellow oil. Yield 564 mg, 54%.

**$^1$H NMR** (400 MHz, $CDCl_3$) δ 8.17 (s, 1H), 7.51 (dd, *J* = 8.5, 7.1 Hz, 1H), 7.11 (d, *J* = 7.1 Hz, 1H), 6.96 (d, *J* = 8.5 Hz, 1H), 4.93 (m, 1H), 3.65 (m, 10H), 3.55 (t, *J* = 6.0 Hz, 2H), 3.43 (t, *J* = 6.6 Hz, 2H), 3.24 (m, 2H), 2.96 – 2.68 (m, 3H), 2.15 (m, 1H), 1.96 (m, 2H), 1.77 (m, 2H), 1.46 (s, 9H).

**MS** *m/z* (ES+) calcd for $C_{28}H_{40}N_4O_9$ [M+ $H^+$] 577.3, found 577.7.

**General procedure III: Boc-deprotection and HATU-mediated amide coupling.**

The corresponding mono-Boc protected linker coupled pomalidomide was dissolved in $CH_2Cl_2$ (3 mL) and then trifluoroacetic acid (1 mL) was added dropwise at 0 °C. The mixture was then stirred at rt for 2 h. Then the solvent was removed, washed by $NaHCO_3$ and extracted by $CH_2Cl_2$. The orange oily residue was further dried under vacuum. The carboxylic acid (1 equiv.), DIPEA (2 equiv.) and HATU (1.5 equiv.) were dissolved in 10 mL $CH_2Cl_2$. After 5 min, deprotected amine (1.2 equiv.) was dissolved in 5 mL $CH_2Cl_2$ and added to the same bottle. The mixture was stirred at rt overnight. Then, the mixture was poured into $H_2O$ (10 mL) and extracted by $CH_2Cl_2$ (3 × 10 mL). The organic phase was collected, dried over anhydrous sodium sulfate and evaporated under reduced pressure. The residue was purified by column chromatography (gradient of $CH_2Cl_2$/MeOH 80:1 to 30:1).

**(4R)-N-(6-((2-(2,6-dioxopiperidin-3-yl)-1,3-dioxoisoindolin-4-yl)amino)hexyl)-4-((3S,8S,9S,10R,13R,14S,17R)-3-hydroxy-10,13-dimethyl-2,3,4,7,8,9,10,11,12,13,14,15,16,17-tetradecahydro-1H-cyclopenta[a]phenanthren-17-yl)pentanamide (C1)**

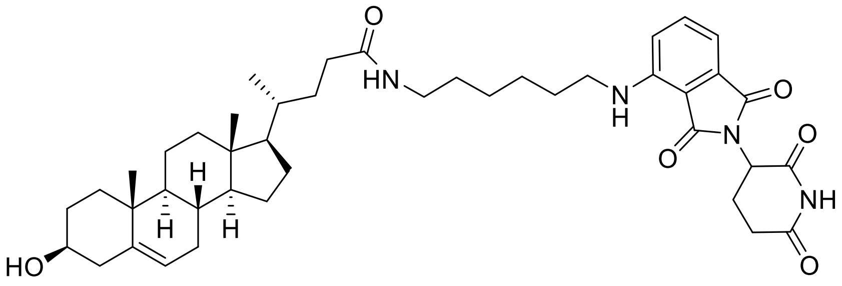


**4** (80 mg, 0.21 mmol) and **11** (120 mg, 0.252 mmol) were combined according to general procedure III. The product was a yellow solid. (73 mg, 48%).

**$^{1}$H NMR** (400 MHz, $CDCl_3$) δ 8.48 (d, *J* = 5.6 Hz, 1H), 7.47 (dd, *J* = 8.5, 7.1 Hz, 1H), 7.06 (d, *J* = 7.1 Hz, 1H), 6.85 (d, *J* = 8.5 Hz, 1H), 6.21 (t, *J* = 5.6 Hz, 1H), 5.57 (d, *J* = 6.8 Hz, 1H), 5.32 (dt, *J* = 4.2, 1.7 Hz, 1H), 4.96 – 4.84 (m, 1H), 3.50 (m, 1H), 3.29 – 3.18 (m, 4H), 2.91 – 2.64 (m, 3H), 2.31 – 2.16 (m, 3H), 2.15 – 2.00 (m, 2H), 2.00 – 1.92 (m, 2H), 1.91 – 1.76 (m, 5H), 1.68 – 1.60 (m, 2H), 1.58 – 1.24 (m, 15H), 1.18 – 1.02 (m, 4H), 0.98 (s, 3H), 0.91 (d, *J* = 6.4 Hz, 4H), 0.65 (s, 3H).

**$^{13}$C NMR** (101 MHz, $CDCl_3$) δ 173.63, 171.23, 169.51, 168.51, 167.61, 146.93, 140.79, 136.12, 132.45, 121.55, 116.63, 111.39, 109.82, 71.69, 56.70, 55.80, 50.05, 48.85, 42.51, 42.34, 42.23, 39.73, 39.32, 37.24, 36.45, 35.49, 33.64, 31.84, 31.58, 31.40, 29.58, 29.09, 28.16, 26.59, 26.56, 24.23, 22.78, 21.04, 19.38, 18.41, 11.86.

**HRMS** (ESI+) *m/z* found 729.4585 $[M+H]^+$, $C_{43}H_{60}N_4O_6$ calculated 729.4591 (Δ = -0.82 ppm)

| Retention time | Area | Rel. Area(%) | Found (m/z) |
|---|---|---|---|
| 2.18 | 2263774 | 100 | 729.26 |

**(4R)-N-(2-(2-(2-((2-(2,6-dioxopiperidin-3-yl)-1,3-dioxoisoindolin-4-yl)amino)ethoxy)ethoxy)ethyl)-4-((3S,8S,9S,10R,13R,14S,17R)-3-hydroxy-10,13-dimethyl-2,3,4,7,8,9,10,11,12,13,14,15,16,17-tetradecahydro-1H-cyclopenta[a]phenanthren-17-yl)pentanamide (C2)**

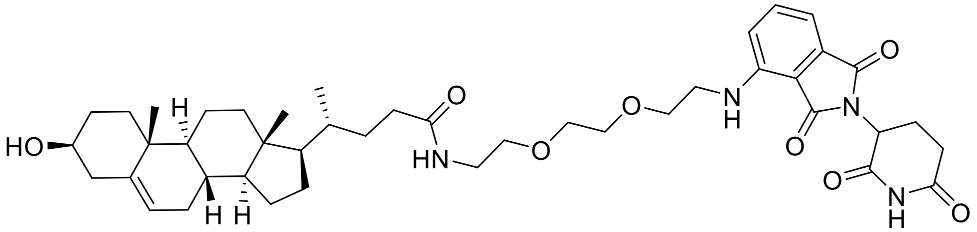

**4** (80 mg, 0.21 mmol) and **12** (127 mg, 0.252 mmol) were combined according to general procedure III. The product was a yellow solid. (83 mg, 52%).

**$^1$H NMR** (400 MHz, $CDCl_3$) δ 8.89 (s, 1H), 7.47 (dd, *J* = 8.5, 7.1 Hz, 1H), 7.09 (d, *J* = 7.1 Hz, 1H), 6.88 (d, *J* = 8.5 Hz, 1H), 6.50 (s, 1H), 6.14 (t, *J* = 5.4 Hz, 1H), 5.36 – 5.25 (m, 1H), 4.98 – 4.85 (m, 1H), 3.71 (t, *J* = 5.3 Hz, 2H), 3.64 (d, *J* = 2.2 Hz, 4H), 3.56 (t, *J* = 5.1 Hz, 2H), 3.52 – 3.47 (m, 1H), 3.47 – 3.40 (m, 4H), 2.89 – 2.66 (m, 3H), 2.26 – 2.19 (m, 2H), 2.14 – 2.02 (m, 3H), 1.97 – 1.90 (m, 2H), 1.85 – 1.77 (m, 3H), 1.60 – 1.35 (m, 7H), 1.35 – 1.10 (m, 4H), 1.10 – 0.98 (m, 4H), 0.97 (s, 3H), 0.87 (d, *J* = 6.3 Hz, 4H), 0.63 (s, 3H).

**$^{13}$C NMR** (101 MHz, $CDCl_3$) δ 174.00, 171.52, 169.48, 168.74, 167.65, 146.80, 140.89, 136.19, 132.62, 121.65, 116.78, 111.85, 110.45, 71.76, 70.72, 70.14, 70.08, 69.32, 56.78, 55.88, 50.15, 48.98, 42.41, 42.38, 42.32, 39.82, 39.33, 37.34, 36.56, 35.58, 33.58, 31.94, 31.81, 31.67, 31.48, 28.23, 24.34, 22.95, 21.14, 19.48, 18.47, 11.95.

**HRMS** (ESI+) *m/z* found 761.4483 $[M+H]^+$, $C_{43}H_{60}N_4O_8$ calculated 761.4489 (Δ = -0.78 ppm)

| Retention time | Area | Rel. Area(%) | Found (m/z) |
|---|---|---|---|
| 1.97 | 1411014 | 100 | 761.41 |

**(4R)-N-(3-(4-(3-((2-(2,6-dioxopiperidin-3-yl)-1,3-dioxoisoindolin-4-yl)amino)propoxy)butoxy)propyl)-4-((3S,8S,9S,10R,13R,14S,17R)-3-hydroxy-10,13-dimethyl-2,3,4,7,8,9,10,11,12,13,14,15,16,17-tetradecahydro-1H-cyclopenta[a]phenanthren-17-yl)pentanamide (C3)**

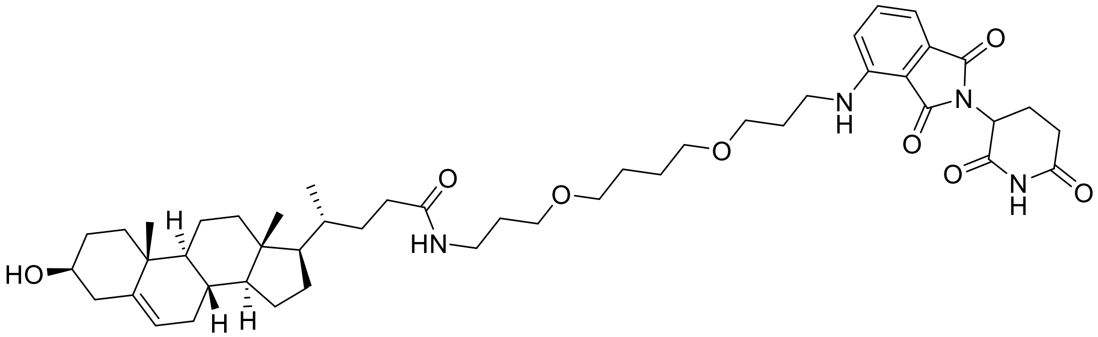


**4** (80 mg, 0.21 mmol) and **13** (145 mg, 0.252 mmol) were combined according to general procedure III. The product was a yellow solid. (51 mg, 30%).

**$^1$H NMR** (400 MHz, $CDCl_3$) δ 8.62 (s, 1H), 7.47 (dd, *J* = 8.5, 7.1 Hz, 1H), 7.06 (d, *J* = 7.1 Hz, 1H), 6.89 (d, *J* = 8.6 Hz, 1H), 6.17 (t, *J* = 5.5 Hz, 1H), 5.34 – 5.29 (m, 1H), 4.92 – 4.84 (m, 1H), 3.54 – 3.47 (m, 5H), 3.44 (d, *J* = 5.8 Hz, 4H), 3.39 – 3.31 (m, 4H), 2.88 – 2.70 (m, 3H), 2.30 – 2.09 (m, 4H), 2.06 – 1.99 (m, 2H), 1.98 – 1.88 (m, 4H), 1.84 – 1.72 (m, 6H), 1.67 – 1.61 (m, 4H), 1.49 – 1.38 (m, 7H), 1.32 – 1.23 (m, 2H), 1.16 – 1.02 (m, 4H), 0.98 (s, 3H), 0.90 (d, *J* = 6.5 Hz, 4H), 0.65 (s, 3H).

**$^{13}$C NMR** (101 MHz, $CDCl_3$) δ 173.84, 171.44, 169.43, 168.67, 167.75, 147.05, 140.90, 136.17, 132.60, 121.65, 116.69, 111.44, 109.98, 71.79, 71.09, 70.96, 69.78, 68.55, 56.82, 55.90, 50.17, 48.94, 42.45, 42.33, 40.46, 39.85, 38.24, 37.35, 36.57, 35.61, 33.75, 31.97, 31.96, 31.68, 31.50, 29.46, 29.21, 28.26, 26.58, 26.50, 24.35, 22.93, 21.16, 19.49, 18.50, 11.97.

**HRMS** (ESI+) *m/z* found 817.5110 $[M+H]^+$, $C_{47}H_{68}N_4O_8$ calculated 817.5115 (Δ = -0.61 ppm)

| Retention time | Area | Rel. Area(%) | Found (m/z) |
|---|---|---|---|
| 1.93 | 9198 | <1 | |
| 2.2 | 1967868 | >99 | 817.58 |

**(4R)-4-((3S,8S,9S,10R,13R,14S,17R)-3-hydroxy-10,13-dimethyl-2,3,4,7,8,9,10,11,12,13,14,15,16,17-tetradecahydro-1H-cyclopenta[a]phenanthren-17-yl)-N-(3-(4-(3-((2-(1-methyl-2,6-dioxopiperidin-3-yl)-1,3-dioxoisoindolin-4-yl)amino)propoxy)butoxy)propyl)pentanamide (C3_Me)**

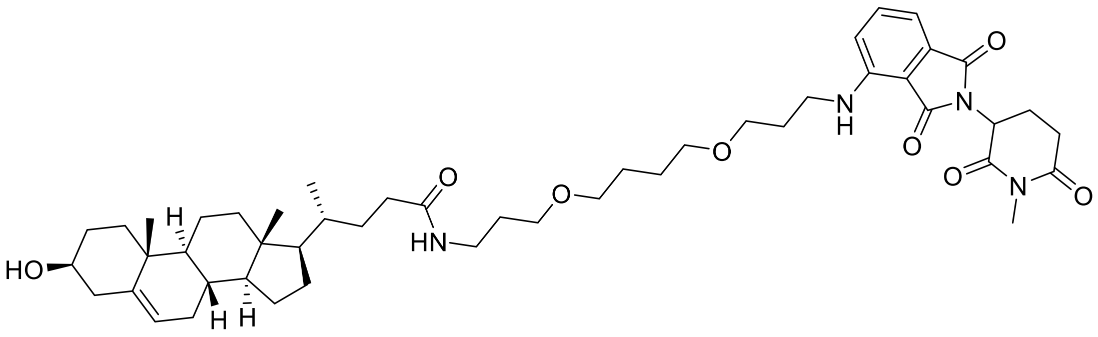

**4** (80 mg, 0.21 mmol) and **14** (141 mg, 0.252 mmol) were combined according to general procedure III. The product was a yellow solid. (70 mg, 41%).

**$^{1}$H NMR** (400 MHz, $CDCl_3$) δ 7.47 (dd, *J* = 8.6, 7.1 Hz, 1H), 7.07 (d, *J* = 7.1, 0.6 Hz, 1H), 6.90 (dd, *J* = 8.6, 0.6 Hz, 1H), 6.19 – 6.07 (m, 1H), 5.34 – 5.30 (m, 1H), 4.93 – 4.85 (m, 1H), 3.54 – 3.45 (m, 5H), 3.45 – 3.40 (m, 4H), 3.40 – 3.31 (m, 4H), 3.19 (s, 3H), 2.98 – 2.93 (m, 1H), 2.78 – 2.70 (m, 2H), 2.29 – 2.15 (m, 3H), 2.09 – 1.97 (m, 3H), 1.96 – 1.86 (m, 4H), 1.86 – 1.70 (m, 6H), 1.67 – 1.61 (m, 4H), 1.52 – 1.38 (m, 6H), 1.39 – 1.21 (m, 3H), 1.17 – 1.02 (m, 4H), 0.98 (s, 3H), 0.91 (d, *J* = 6.5 Hz, 4H), 0.65 (s, 3H).

**$^{13}$C NMR** (101 MHz, $CDCl_3$) δ 173.72, 171.37, 169.64, 169.16, 167.88, 147.05, 140.90, 136.14, 132.66, 121.68, 116.65, 111.44, 110.10, 71.81, 71.03, 69.99, 68.35, 56.84, 55.93, 50.19, 49.71, 42.47, 42.38, 40.29, 39.87, 38.39, 37.37, 36.59, 35.64, 33.77, 32.03, 31.98, 31.73, 29.60, 29.24, 28.29, 27.36, 26.68, 26.53, 24.37, 22.26, 21.17, 19.51, 18.53, 11.99.

**HRMS** (ESI+) *m/z* found 831.5270 $[M+H]^+$, $C_{48}H_{70}N_4O_8$ calculated 831.5272 (Δ = -0.24 ppm)

| Retention time | Area | Rel. Area(%) | Found (m/z) |
|---|---|---|---|
| 1.32 | 72600 | 4.5 | |
| 2.33 | 1550000 | 95.5 | 831.41 |

***tert*-butyl (3-(4-(3-(2-(adamantan-1-yl)acetamido)propoxy)butoxy)propyl)carbamate (16)**

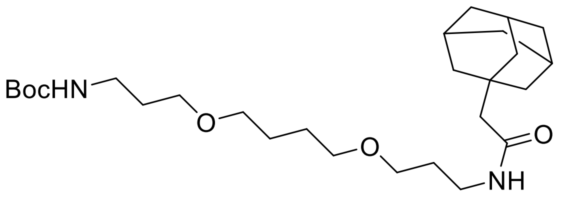

1-Adamantylacetic acid (200 mg, 1.03 mmol) and **9** (376 mg, 1.23 mmol) were combined according to general procedure III. The product was a colorless oil. (344 mg, 69%).

**$^{1}$H NMR** (400 MHz, DMSO-*d*$_6$) δ 7.64 (t, *J* = 5.6 Hz, 1H), 6.76 (t, *J* = 5.7 Hz, 1H), 3.54 – 3.45 (m, 1H), 3.38 – 3.33 (m, 7H), 3.06 (q, *J* = 6.6 Hz, 2H), 2.96 (q, *J* = 6.6 Hz, 2H), 1.91 (s, 3H), 1.80 (s, 2H), 1.69 – 1.48 (m, 20H), 1.38 (s, 9H).

**$^{13}$C NMR** (101 MHz, DMSO-*d*$_6$): δ 170.2, 155.4, 132.7, 70.3, 68.2, 50.6, 42.6, 37.7, 36.9, 36.1, 32.6, 30.2, 30.0, 28.7, 28.5, 26.5.

**MS** *m/z* (ES+) calcd for $C_{27}H_{48}N_2O_5$ [M+ H$^+$] 481.4, found 481.4.

**(R)-N-(3-(4-(3-(2-(adamantan-1-yl)acetamido)propoxy)butoxy)propyl)-4-((3S,8S,9S,10R,13R,14S,17R)-3-hydroxy-10,13-dimethyl-2,3,4,7,8,9,10,11,12,13,14,15,16,17-tetradecahydro-1H-cyclopenta[a]phenanthren-17-yl)pentanamide (C3_HyT, Oxybipin-1)**

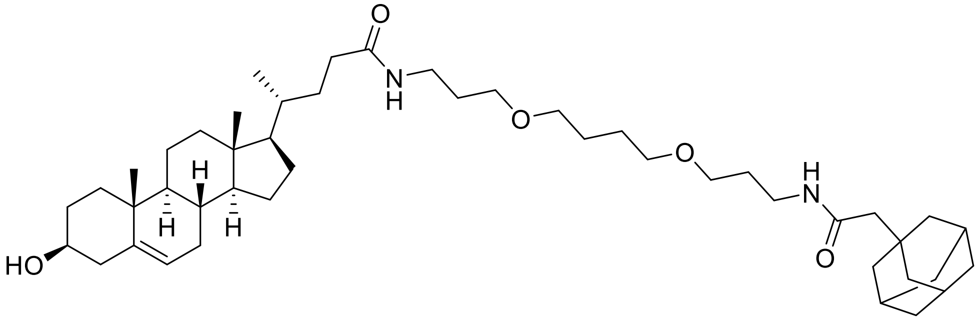

**4** (45 mg, 0.12 mmol) and **16** (55 mg, 0.144 mmol) were combined according to general procedure III. The product was a transparent oil. (44 mg, 50%).

**$^{1}$H NMR** (400 MHz, CDCl$_3$) δ 6.15 (t, *J* = 5.4 Hz, 1H), 6.04 (t, *J* = 5.4 Hz, 1H), 5.35 – 5.29 (m, 1H), 3.53 – 3.45 (m, 5H), 3.45 – 3.37 (m, 4H), 3.32 (q, *J* = 6.3 Hz, 4H), 2.31 – 2.13 (m, 4H), 2.06 – 1.98 (m, 2H), 1.97 – 1.91 (m, 4H), 1.88 (s, 2H), 1.84 – 1.77 (m, 3H), 1.77 – 1.70 (m, 5H), 1.69 (s, 1H), 1.66 (s, 2H), 1.64 – 1.60 (m, 6H), 1.60 – 1.58 (m, 7H), 1.56 – 1.36 (m, 7H), 1.34 – 1.17 (m, 2H), 1.15 – 1.02 (m, 4H), 0.98 (s, 3H), 0.91 (d, *J* = 6.5 Hz, 4H), 0.65 (s, 3H).

**$^{13}$C NMR** (101 MHz, CDCl$_3$) δ 173.55, 170.89, 140.85, 121.52, 71.64, 70.82, 69.81, 69.74, 56.73, 55.83, 51.93, 50.08, 42.67, 42.36, 42.28, 39.76, 38.08, 37.98, 37.27, 36.77, 36.49, 35.53, 33.67, 32.70, 31.87, 31.61, 29.36, 29.22, 28.64, 28.18, 26.58, 26.54, 24.26, 21.06, 19.40, 18.42, 11.88.

**HRMS** (ESI+) *m/z* found 737.5822 [M+H]$^+$, $C_{46}H_{76}N_2O_5$ calculated 737.5834 (Δ = -1.62 ppm)

$[\alpha]_D^{20}$: -24° (c=0.9, EtOH)

**_tert_-butyl (3-(4-(3-((R)-4-((3S,8S,9S,10R,13R,14S,17R)-3-hydroxy-10,13-dimethyl-2,3,4,7,8,9,10,11,12,13,14,15,16,17-tetradecahydro-1H-cyclopenta[a]phenanthren-17-yl)pentanamido)propoxy)butoxy)propyl)carbamate (C3_Boc, Oxybipin-2)**

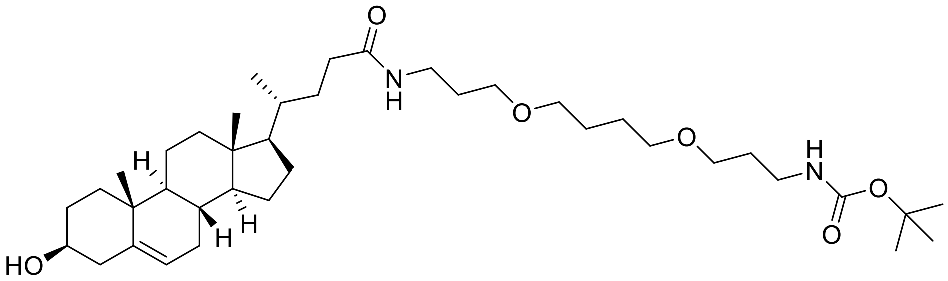

**4** (90 mg, 0.24 mmol) and **9** (88 mg, 0.288 mmol) were combined according to general procedure III. The product was a transparent oil. (130 mg, 82%).

**$^1$H NMR** (400 MHz, $CDCl_3$) δ 6.13 (s, 1H), 5.37 – 5.30 (m, 1H), 4.88 (s, 1H), 3.53 – 3.45 (m, 5H), 3.45 – 3.39 (m, 4H), 3.34 (q, *J* = 6.0 Hz, 2H), 3.20 (q, *J* = 6.1 Hz, 2H), 2.31 – 2.16 (m, 3H), 2.04 – 1.92 (m, 4H), 1.86 – 1.71 (m, 8H), 1.65 – 1.60 (m, 4H), 1.59 – 1.45 (m, 5H), 1.43 (s, 10H), 1.33 – 1.24 (m, 2H), 1.24 – 1.00 (m, 5H), 0.99 (s, 3H), 0.92 (d, *J* = 6.5 Hz, 4H), 0.66 (s, 3H).

**$^{13}$C NMR** (101 MHz, $CDCl_3$) δ 173.71, 156.13, 140.92, 121.72, 71.84, 70.99, 70.82, 69.99, 69.31, 56.86, 55.96, 50.21, 42.49, 42.40, 39.89, 38.37, 37.39, 36.61, 35.66, 33.78, 32.00, 31.75, 29.88, 29.22, 28.56, 28.30, 26.65, 26.61, 24.39, 21.19, 19.52, 18.54, 12.01.

**HRMS** (ESI+) *m/z* found 661.5153 $[M+H]^+$, $C_{39}H_{68}N_2O_6$ calculated 661.5156 (Δ = -0.45 ppm)

$[\alpha]_D^{20}$: -31.2° (c=1.00, EtOH)

**(4R)-N-(3-(2-(2-(3-((2-(2,6-dioxopiperidin-3-yl)-1,3-dioxoisoindolin-4-yl)amino)propoxy)ethoxy)ethoxy)propyl)-4-((3S,8S,9S,10R,13R,14S,17R)-3-hydroxy-10,13-dimethyl-2,3,4,7,8,9,10,11,12,13,14,15,16,17-tetradecahydro-1H-cyclopenta[a]phenanthren-17-yl)pentanamide (C4)**

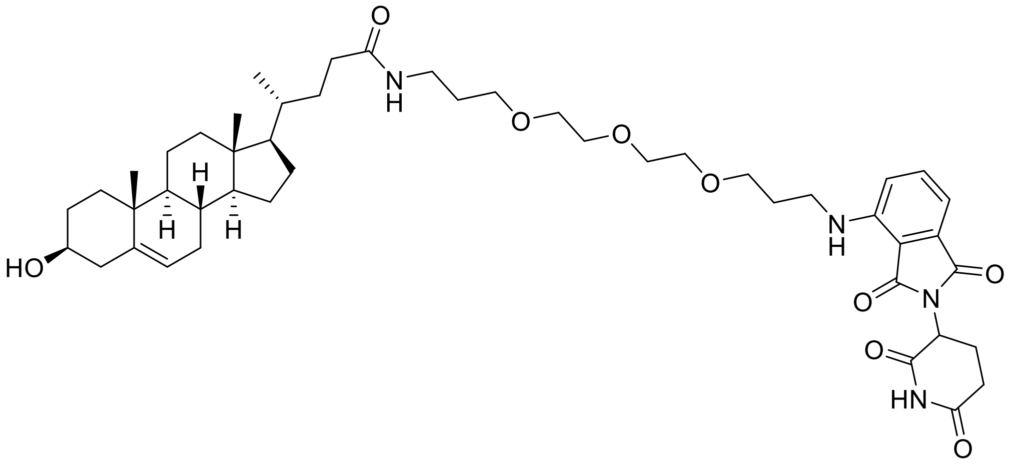

**4** (80 mg, 0.21 mmol) and **15** (210 mg, 0.252 mmol) were combined according to general procedure III. The product was a yellow solid. (100 mg, 57%).

**$^{1}$H NMR** (400 MHz, $CDCl_3$) δ 8.53 (s, 1H), 7.47 (dd, *J* = 8.5, 7.1 Hz, 1H), 7.07 (d, *J* = 7.1 Hz, 1H), 6.92 (d, *J* = 8.5 Hz, 1H), 6.26 (t, *J* = 5.5 Hz, 1H), 5.33 – 5.31 (m, 1H), 4.92 – 4.87 (m, 1H), 3.68 – 3.52 (m, 12H), 3.52 – 3.44 (m, 1H), 3.40 – 3.30 (m, 4H), 2.88 – 2.67 (m, 3H), 2.26 – 2.16 (m, 3H), 2.14 – 2.00 (m, 3H), 1.97 – 1.88 (m, 4H), 1.85 – 1.70 (m, 6H), 1.57 – 1.39 (m, 7H), 1.33 – 1.21 (m, 2H), 1.14 – 1.01 (m, 4H), 0.98 (s, 3H), 0.97 – 0.90 (m, 4H), 0.65 (s, 3H).

**$^{13}$C NMR** (101 MHz, $CDCl_3$) δ 173.98, 171.35, 169.45, 168.63, 167.74, 147.06, 140.90, 136.22, 132.61, 121.68, 116.75, 111.49, 110.00, 71.81, 70.59, 70.56, 70.54, 70.18, 70.09, 69.05, 56.83, 55.95, 50.18, 48.96, 42.46, 42.36, 40.36, 39.86, 38.02, 37.36, 36.59, 35.64, 33.63, 31.97, 31.71, 31.52, 29.37, 29.03, 28.27, 24.37, 22.94, 21.17, 19.51, 18.53, 11.99.

**HRMS** (ESI+) *m/z* found 833.5055 $[M+H]^+$, $C_{47}H_{68}N_4O_9$ calculated 833.5065 (Δ = -1.19 ppm)

| Retention time | Area | Rel. Area(%) | Found (m/z) |
|---|---|---|---|
| 2.06 | 1795766 | 100 | 833.72 |

**(R)-4-((3S,8S,9S,10R,13R,14S,17R)-3-hydroxy-10,13-dimethyl-2,3,4,7,8,9,10,11,12,13,14,15,16,17-tetradecahydro-1H-cyclopenta[a]phenanthren-17-yl)-N-methylpentanamide (C5)**

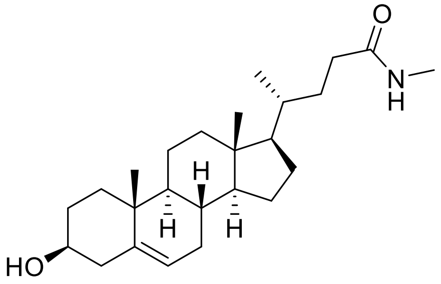

**4** (50 mg, 0.133 mmol) and methylamine (5 mg, 0.16 mmol) were combined according to general procedure III. The product was a white powder. (34 mg, 65%).

**$^1$H NMR** (400 MHz, $CDCl_3$) δ 5.63 – 5.46 (m, 1H), 5.39 – 5.31 (m, 1H), 3.58 – 3.47 (m, 1H), 2.81 (d, *J* = 3.3 Hz, 3H), 2.29 – 2.23 (m, 2H), 2.10 – 1.98 (m, 4H), 1.87 – 1.81 (m, 3H), 1.62 – 1.41 (m, 7H), 1.35 – 1.24 (m, 4H), 1.15 – 1.02 (m, 4H), 1.00 (s, 3H), 0.95 – 0.91 (m, 4H), 0.68 (d, *J* = 2.4 Hz, 3H).

**$^{13}$C NMR** (101 MHz, $CDCl_3$) δ 174.63, 140.88, 121.82, 71.94, 56.87, 55.96, 50.22, 42.52, 42.43, 39.90, 37.39, 36.64, 35.70, 33.63, 32.02, 31.79, 28.32, 26.57, 24.41, 21.22, 19.54, 18.56, 12.03.

**HRMS** (ESI+) *m/z* found 388.3216 $[M+H]^+$, $C_{25}H_{41}NO_2$ calculated 388.3212 (Δ = -0.07 ppm)

$[\alpha]_D^{20}$: -33.8° (c=0.7, EtOH)

***tert*-butyl (3-(4-(3-(5-((3aS,4S,6aR)-2-oxohexahydro-1H-thieno[3,4-d]imidazol-4-yl)pentanamido)propoxy)butoxy)propyl)carbamate (17)**

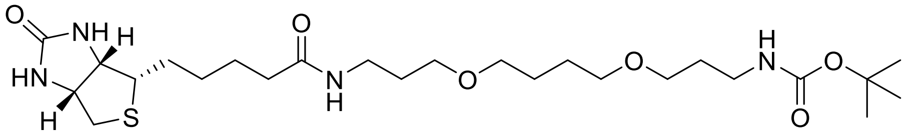

D-biotin (200 mg, 0.82 mmol) and **9** (300 mg, 0.98 mmol) were combined according to general procedure III. The product was a white powder. (265 mg, 61%)

The synthesis of this compound has been reported previously[7].

**$^1$H NMR** (400 MHz, $CDCl_3$) δ 4.58 – 4.50 (m, 1H), 4.38 – 4.30 (m, 1H), 3.56 – 3.46 (m, 6H), 3.44 (s, 4H), 3.25 – 3.18 (m, 2H), 3.18 – 3.16 (m, 1H), 2.95 – 2.88 (m, 1H), 2.80 – 2.74 (m, 1H), 2.24 – 2.18 (m, 2H), 1.80 – 1.72 (m, 5H), 1.71 – 1.57 (m, 7H), 1.44 (s, 11H).

**MS** *m/z* (ES+) calcd for $C_{25}H_{46}N_4O_6S$ $[M+H^+]$ 531.3 found 531.1.

**_tert_-butyl ((S)-1-((2S,4R)-4-hydroxy-2-((4-(4-methylthiazol-5-yl)benzyl)carbamoyl)pyrrolidin-1-yl)-3,3-dimethyl-1-oxobutan-2-yl)carbamate (18)**

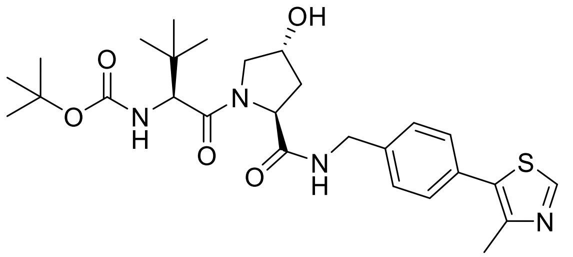

This compound was synthesized following a procedure described in the literature[8].

**$^1$H NMR** (400 MHz, $CDCl_3$): δ 8.73 (s, 1H), 7.45 (t, J = 5.9 Hz, 1H), 7.38 – 7.31 (m, 4H), 5.18 (d, J = 9.0 Hz, 1H), 4.76 (t, J = 7.9 Hz, 1H), 4.60 – 4.49 (m, 2H), 4.31 (dd, J = 15.0, 5.1 Hz, 1H), 4.15 (d, J = 9.1 Hz, 1H), 4.07 (d, J = 11.4 Hz, 1H), 3.58 (dd, J = 11.4, 3.6 Hz, 1H), 2.59 – 2.53 (m, 1H), 2.52 (s, 3H), 2.12 (dd, J = 13.6, 7.9 Hz, 1H), 1.40 (s, 9H), 0.91 (s, 9H).

**$^{13}$C NMR** (101 MHz, $CDCl_3$) δ 172.85, 170.74, 156.58, 150.63, 138.44, 132.12, 130.74, 129.64, 128.31, 80.59, 70.26, 59.04, 58.37, 56.59, 43.37, 35.80, 34.82, 28.43, 26.47, 15.93.

**General procedure IV: Boc-deprotection and HATU-mediated amide coupling.**

The Boc-protected amine (1 equiv.) was dissolved in DCM followed by the addition of 4 M HCl in dioxane to make a 1 M HCl solution. The solution was set to stir at rt under $N_2$ atmosphere for 2 h. This produced the deprotected amine as a precipitated HCl salt.

A solution/suspension of the carboxylic acid (1.3 equiv.), HATU (1.4 equiv.), and DIPEA (5 equiv.) in 10 mL DCM was set to stir for 5 min before dissolving the amine (1 equiv.) in DCM and adding this amine solution to the acid-HATU solution. The mixture was set to stir at rt overnight. The mixture was diluted 4-fold and washed with equal volumes of sat. $NaHCO_3$, 0.5 M HCl, and brine respectively. The organic solution was dried with anhydrous $Na_2SO_4$, filtered, and evaporated under reduced pressure. The crude product was purified by flash chromatography and dried under vacuum.

***tert*-Butyl (6-(((S)-1-((2S,4R)-4-hydroxy-2-((4-(4-methylthiazol-5-yl)benzyl)carbamoyl)pyrrolidin-1-yl)-3,3-dimethyl-1-oxobutan-2-yl)amino)-6-oxohexyl)carbamate (19)**

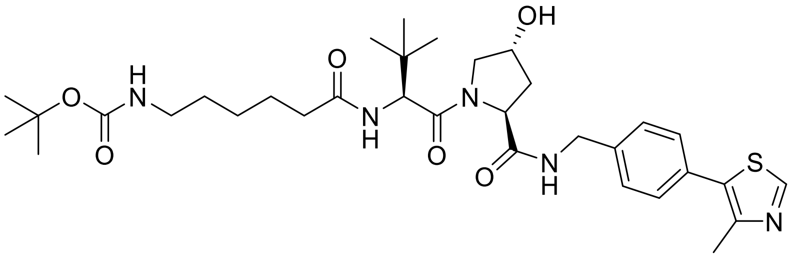


6-((*tert*-butoxycarbonyl)amino)hexanoic acid (213.5 mg, 0.92 mmol) and **18** (365.0 mg, 0.69 mmol) were combined according to general procedure IV. The product was a creamy-white solid (314.8 mg, 71%).

**$^{1}$H NMR** (400 MHz, MeOD): δ 8.88 (s, 1H), 7.49 – 7.39 (m, 4H), 4.63 (s, 1H), 4.60 – 4.47 (m, 3H), 4.36 (d, J = 15.5 Hz, 1H), 3.91 (d, J = 10.9 Hz, 1H), 3.80 (dd, J = 11.0, 3.9 Hz, 1H), 3.02 (t, J = 7.0 Hz, 2H), 2.48 (s, 3H), 2.35 – 2.18 (m, 3H), 2.12 – 2.04 (m, 1H), 1.68 – 1.56 (m, 2H), 1.52 – 1.44 (m, 2H), 1.42 (s, 9H), 1.38 – 1.28 (m, 2H), 1.04 (s, 9H).

**$^{13}$C NMR** (101 MHz, MeOD): δ 175.96, 174.49, 172.36, 152.85, 149.04, 140.30, 133.43, 131.52, 130.37, 128.99, 71.09, 60.82, 59.00, 58.01, 43.70, 41.22, 38.92, 36.54, 30.64, 28.79, 27.49, 27.04, 26.70, 15.80.

The data matched those reported in the literature[9].

***tert*-Butyl (2-(2-(2-(((S)-1-((2S,4R)-4-hydroxy-2-((4-(4-methylthiazol-5-yl)benzyl)carbamoyl)pyrrolidin-1-yl)-3,3-dimethyl-1-oxobutan-2-yl)amino)-2-oxoethoxy)ethoxy)ethyl)carbamate (20)**

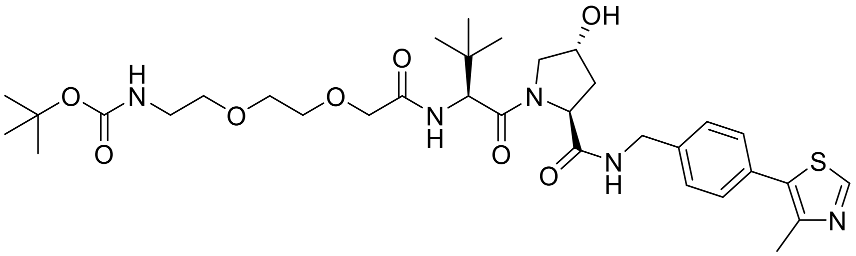

2,2-dimethyl-4-oxo-3,8,11-trioxa-5-azatridecan-13-oic acid (165.0 mg, 0.63 mmol) and **18** (253.3 mg, 0.48 mmol) were combined according to general procedure IV. The product was a white solid (103.3 mg, 32%).

**$^{1}$H NMR** (400 MHz, DMSO-*d*$_6$): δ 8.98 (s, 1H), 8.57 (t, J = 6.0 Hz, 1H), 7.46 – 7.42 (m, 1H), 7.41 – 7.37 (m, 4H), 6.76 (t, J = 5.8 Hz, 1H), 5.15 (d, J = 3.6 Hz, 1H), 4.57 (d, J = 9.6 Hz, 1H), 4.47 – 4.32 (m, 3H), 4.26 (dd, J = 15.8, 5.7 Hz, 1H), 3.96 (s, 2H), 3.67 (dd, J = 10.7, 4.0 Hz, 1H), 3.63 – 3.57 (m, 3H), 3.57 – 3.50 (m, 2H), 3.44 – 3.36 (m, 2H), 3.08 (q, J = 6.0 Hz, 2H), 2.44 (s, 3H), 2.06 (dd, J = 13.1, 7.7 Hz, 1H), 1.94 – 1.87 (m, 1H), 1.35 (s, 9H), 0.94 (s, 9H).

**$^{13}$C NMR** (101 MHz, DMSO-*d*$_6$): δ 172.18, 169.63, 169.07, 156.04, 151.93, 148.22, 139.88, 131.61, 130.19, 129.18, 127.95, 78.05, 70.86, 70.03, 69.75, 69.33, 59.19, 57.04, 56.15, 42.16, 38.38, 36.19, 28.68, 26.66, 16.40.

The data matched those reported in the literature[10].

***tert*-Butyl ((S)-17-((2S,4R)-4-hydroxy-2-((4-(4-methylthiazol-5-yl)benzyl)carbamoyl)pyrrolidine-1-carbonyl)-18,18-dimethyl-15-oxo-3,6,9,12-tetraoxa-16-azanonadecyl)carbamate (21)**

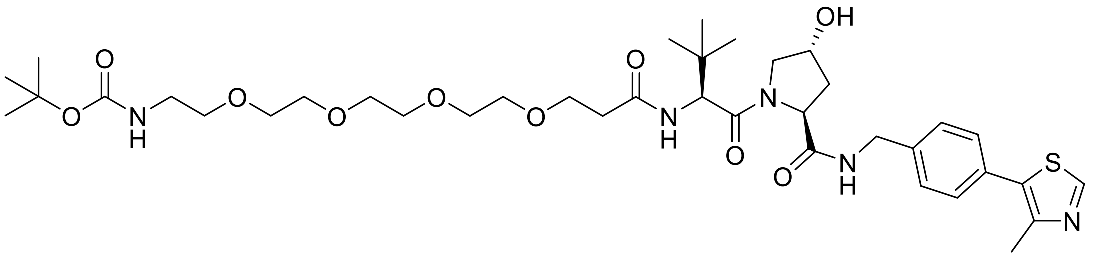

2,2-dimethyl-4-oxo-3,8,11,14,17-pentaoxa-5-azaicosan-20-oic acid (197.9 mg, 0.54 mmol) and **18** (218.3 mg, 0.41 mmol) were combined according to general procedure IV. The product was a white solid (166.6 mg, 52%).

**$^1$H NMR** (400 MHz, MeOD): δ 8.88 (s, 1H), 7.49 – 7.39 (m, 4H), 4.65 (s, 1H), 4.59 – 4.48 (m, 3H), 4.36 (d, J = 15.5 Hz, 1H), 3.89 (d, J = 10.8 Hz, 1H), 3.83 – 3.68 (m, 3H), 3.66 – 3.56 (m, 12H), 3.49 (t, J = 5.6 Hz, 2H), 3.21 (t, J = 5.6 Hz, 2H), 2.62 – 2.54 (m, 1H), 2.51 – 2.43 (m, 4H), 2.22 (m, 1H), 2.08 (m, 1H), 1.43 (s, 9H), 1.04 (s, 9H).

**$^{13}$C NMR** (101 MHz, MeOD): δ 174.49, 173.74, 172.11, 152.86, 149.04, 140.30, 133.42, 131.51, 130.37, 128.99, 71.56, 71.52, 71.43, 71.26, 71.08, 68.29, 60.81, 58.91, 58.00, 43.69, 41.29, 38.93, 37.36, 36.80, 28.77, 27.03, 15.82.

The data matched those reported in the literature[11].

**(2S,4R)-4-hydroxy-1-((S)-2-(6-((R)-4-((3S,8S,9S,10R,13R,14S,17R)-3-hydroxy-10,13-dimethyl-2,3,4,7,8,9,10,11,12,13,14,15,16,17-tetradecahydro-1H-cyclopenta[a]phenanthren-17-yl)pentanamido)hexanamido)-3,3-dimethylbutanoyl)-N-(4-(4-methylthiazol-5-yl)benzyl)pyrrolidine-2-carboxamide (ORD027)**

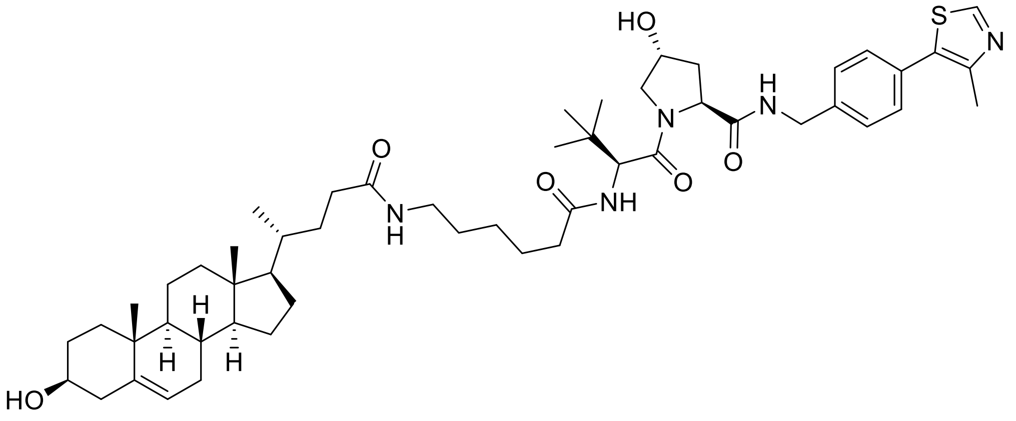

**4** (52.4 mg, 0.14 mmol) and **19** (77.9 mg, 0.12 mmol) were combined according to general procedure IV. The product was a white solid (44.2 mg, 41%).

**$^1$H-NMR** (400 MHz, $CDCl_3$): δ 8.77 (s, 1H), 7.44 (t, J = 6.0 Hz, 1H), 7.35 (s, 4H), 6.27 (d, J = 8.8 Hz, 1H), 5.74 (t, J = 5.9 Hz, 1H), 5.35 – 5.31 (m, 1H), 4.72 (t, J = 8.1 Hz, 1H), 4.59 – 4.48 (m, 3H), 4.35 (m, 1H), 4.07 (d, J = 11.6 Hz, 1H), 3.60 (m, 1H), 3.50 (m, 1H), 3.18 (q, J = 6.7 Hz, 2H), 2.53 (s, 3H), 2.51 – 2.43 (m, 1H), 2.31 – 2.11 (m, 6H), 2.07 – 1.90 (m, 3H), 1.89 – 1.69 (m, 4H), 1.66 – 1.35 (m, 11H), 1.34 – 1.20 (m, 4H), 1.18 – 1.00 (m, 5H), 0.99 (s, 3H), 0.93 (s, 9H), 0.90 (d, J = 6.4 Hz, 4H), 0.65 (s, 3H).

**$^{13}$C NMR** (101 MHz, $CDCl_3$): δ 173.89, 173.74, 171.98, 171.06, 150.70, 147.84, 140.92, 138.61, 132.21, 130.57, 129.60, 128.26, 121.72, 71.83, 70.12, 58.70, 57.59, 57.00, 56.85, 55.94, 50.20, 43.31, 42.49, 42.38, 39.88, 39.38, 37.39, 36.61, 36.26, 35.65, 35.14, 33.76, 31.99, 31.98, 31.73, 29.41, 28.31, 26.55, 26.30, 25.23, 24.38, 21.19, 19.53, 18.56, 15.89, 12.01.

**HRMS** (ESI+) *m/z* found 900.5659 $[M+H]^+$, $C_{52}H_{77}N_5O_6S$ calculated 900.5673 (Δ = 1.55 ppm)

$[\alpha]_D^{20}$: -25.0° (c=0.46, EtOH)

| Retention time | Area | Rel. Area(%) | Found (m/z) |
|---|---|---|---|
| 2.91 | 4274573 | 100 | 900.46 |

**(4R)-1-((2S,16R)-2-(*tert*-butyl)-16-((3S,8S,9S,10R,13R,14S,17R)-3-hydroxy-10,13-dimethyl-2,3,4,7,8,9,10,11,12,13,14,15,16,17-tetradecahydro-1H-cyclopenta[a]phenanthren-17-yl)-4,13-dioxo-6,9-dioxa-3,12-diazaheptadecanoyl)-4-hydroxy-N-(4-(4-methylthiazol-5-yl)benzyl)pyrrolidine-2-carboxamide (ORD033)**

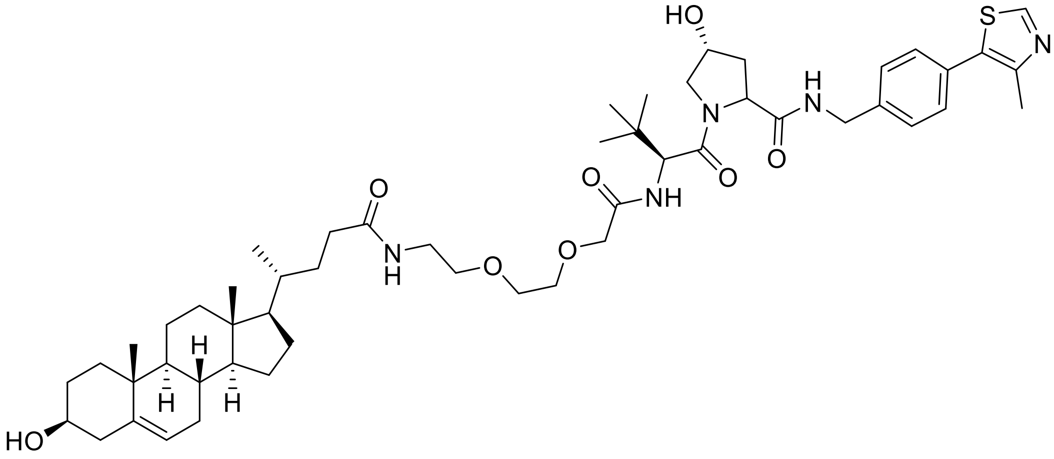

**4** (24.2 mg, 0.064 mmol) and **20** (30.1 mg, 0.044 mmol) were combined according to general procedure IV. The product was a white solid (24.0 mg, 58%)

**$^{1}$H NMR** (400 MHz, MeOD): δ 8.88 (s, 1H), 7.46 – 7.40 (m, 4H), 5.35 – 5.31 (m, 1H), 4.75 (s, 1H), 4.61 – 4.48 (m, 3H), 4.39 – 4.31 (m, 1H), 4.04 (s, 2H), 3.89 – 3.79 (m, 2H), 3.72 (t, *J* = 4.3 Hz, 2H), 3.67 – 3.62 (m, 2H), 3.58 – 3.51 (m, 2H), 3.49 – 3.34 (m, 2H), 3.30 – 3.25 (m, 1H), 2.49 (s, 3H), 2.28 – 1.72 (m, 12H), 1.63 – 1.09 (m, 12H), 1.06 (s, 9H), 1.04 – 1.02 (m, 1H), 1.01 (s, 3H), 0.94 (d, *J* = 6.5 Hz, 3H), 0.91 – 0.87 (m, 1H), 0.70 (s, 3H).

**$^{13}$C NMR** (101 MHz, MeOD): δ 176.94, 174.23, 172.20, 171.70, 152.86, 149.07, 142.22, 140.21, 133.40, 131.60, 130.40, 128.99, 122.43, 72.43, 72.40, 71.22, 70.98, 70.91, 60.88, 58.20, 58.15, 58.11, 57.35, 51.69, 43.76, 43.54, 43.02, 41.12, 40.43, 39.02, 38.54, 37.67, 37.34, 36.83, 34.15, 33.37, 33.25, 33.01, 32.29, 29.23, 27.05, 25.30, 22.18, 19.86, 18.94, 15.89, 12.37.

**HRMS** (ESI+) *m/z* found 932.5575 $[M+H]^+$, $C_{52}H_{77}N_5O_8S$ calculated 932.5571 (Δ = 0.43 ppm)

$[\alpha]_D^{20}$: -37.1° (c=0.36, EtOH).

| Retention time | Area | Rel. Area(%) | Found (m/z) |
|---|---|---|---|
| 2.85 | 1737938 | 100 | 932.53 |

**(2S,4R)-1-((2S,23R)-2-(*tert*-butyl)-23-((3S,8S,9S,10R,13R,14S,17R)-3-hydroxy-10,13-dimethyl-2,3,4,7,8,9,10,11,12,13,14,15,16,17-tetradecahydro-1H-cyclopenta[a]phenanthren-17-yl)-4,20-dioxo-7,10,13,16-tetraoxa-3,19-diazatetracosanoyl)-4-hydroxy-N-(4-(4-methylthiazol-5-yl)benzyl)pyrrolidine-2-carboxamide(ORD031)**

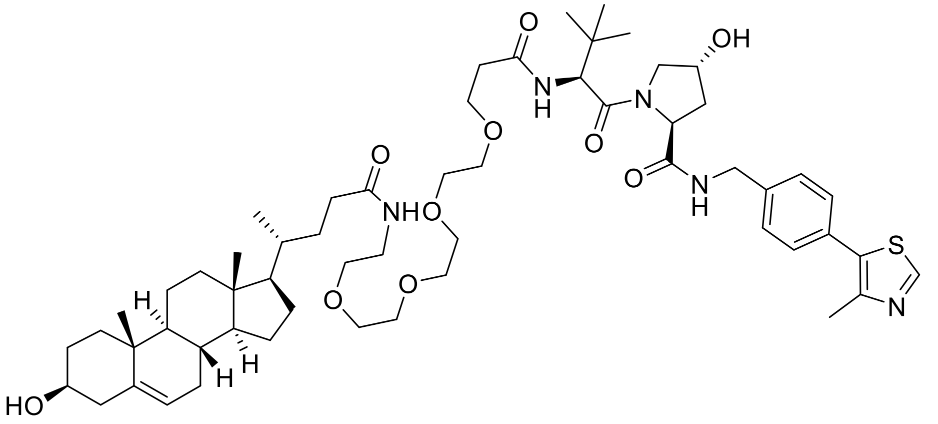


**4** (36.9 mg, 0.098 mmol) and **21** (60.4 mg, 0.077 mmol) were combined according to general procedure IV. The product was a white solid (53.6 mg, 67%).

**$^{1}$H NMR** (400 MHz, $CDCl_3$): δ 8.97 (s, 1H), 7.43 (t, J = 5.9 Hz, 1H), 7.42 – 7.33 (m, 4H), 6.97 (d, J = 8.1 Hz, 1H), 6.24 (t, J = 5.2 Hz, 1H), 5.34 (d, J = 5.5 Hz, 1H), 4.73 (t, J = 8.1 Hz, 1H), 4.59 (dd, J = 15.1, 6.7 Hz, 1H), 4.51 (s, 1H), 4.45 (d, J = 8.2 Hz, 1H), 4.35 (dd, J = 15.1, 5.2 Hz, 1H), 4.13 (d, J = 11.4 Hz, 1H), 3.72 (t, J = 5.7 Hz, 2H), 3.66 – 3.46 (m, 16H), 3.42 (q, J = 5.0 Hz, 2H), 2.59 (s, 3H), 2.58 – 2.42 (m, 3H), 2.33 – 1.72 (m, 12H), 1.62 – 1.21 (m, 9H), 1.19 – 1.01 (m, 5H), 1.00 (s, 3H), 0.94 (s, 9H), 0.92 (d, J = 6.5 Hz, 3H), 0.66 (s, 3H).

**$^{13}$C NMR** (101 MHz, $CDCl_3$): δ 173.98, 172.20, 172.00, 171.02, 151.18, 140.92, 139.42, 129.58, 128.53, 121.78, 71.89, 70.65, 70.58, 70.28, 70.24, 70.13, 67.28, 58.50, 57.95, 56.88, 56.80, 56.00, 50.22, 43.30, 42.51, 42.41, 39.90, 39.34, 37.39, 36.81, 36.63, 36.01, 35.69, 34.86, 33.66, 32.01, 31.91, 31.77, 28.31, 26.58, 24.41, 21.21, 19.54, 18.58, 15.16, 12.03.

**HRMS** (ESI+) *m/z* found 1056.6057 $[M+Na]^+$, $C_{57}H_{88}N_5O_{10}S$ calculated 1056.6071 (Δ = -1.32 ppm)

$[\alpha]_D^{20}$: -18.4° (c=0.48, EtOH).

| Retention time | Area | Rel. Area(%) | Found (m/z) |
|---|---|---|---|
| 2.83 | 4654739 | 100 | 1034.57 |

# C1

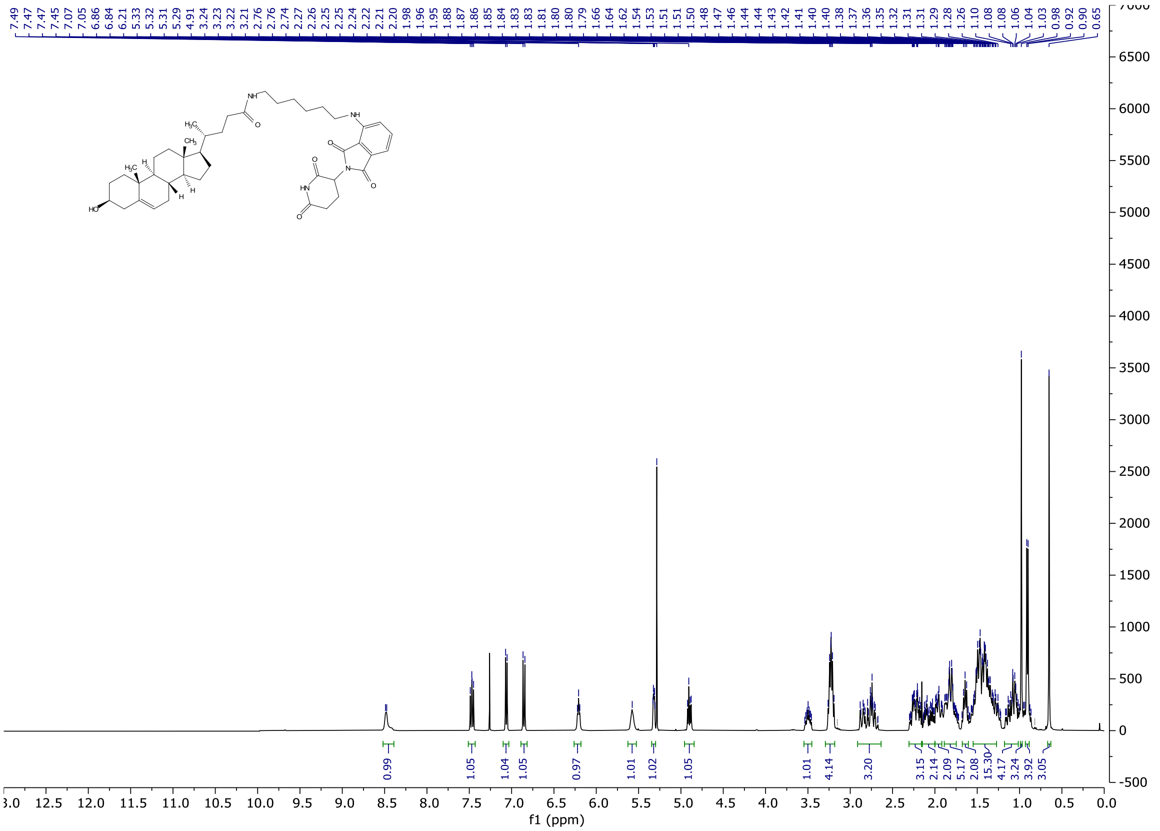
7.49 7.47 7.47 7.45 7.07 7.05 6.86 6.84 6.21 5.33 5.32 5.31 5.29 4.91 3.24 3.23 3.22 3.21 2.76 2.76 2.74 2.27 2.26 2.25 2.25 2.24 2.22 2.21 2.20 1.98 1.96 1.95 1.88 1.87 1.86 1.85 1.84 1.83 1.83 1.81 1.80 1.79 1.66 1.64 1.62 1.54 1.53 1.51 1.51 1.50 1.48 1.47 1.46 1.44 1.44 1.43 1.42 1.41 1.40 1.38 1.37 1.36 1.35 1.32 1.31 1.29 1.28 1.26 1.10 1.08 1.08 1.06 1.04 1.03 0.98 0.92 0.90 0.65
0.99 1.05 1.04 1.05 0.97 1.01 1.02 1.05 1.01 4.14 3.20 3.15 2.14 2.09 5.17 2.08 15.30 4.17 3.24 3.92 3.05
f1 (ppm)

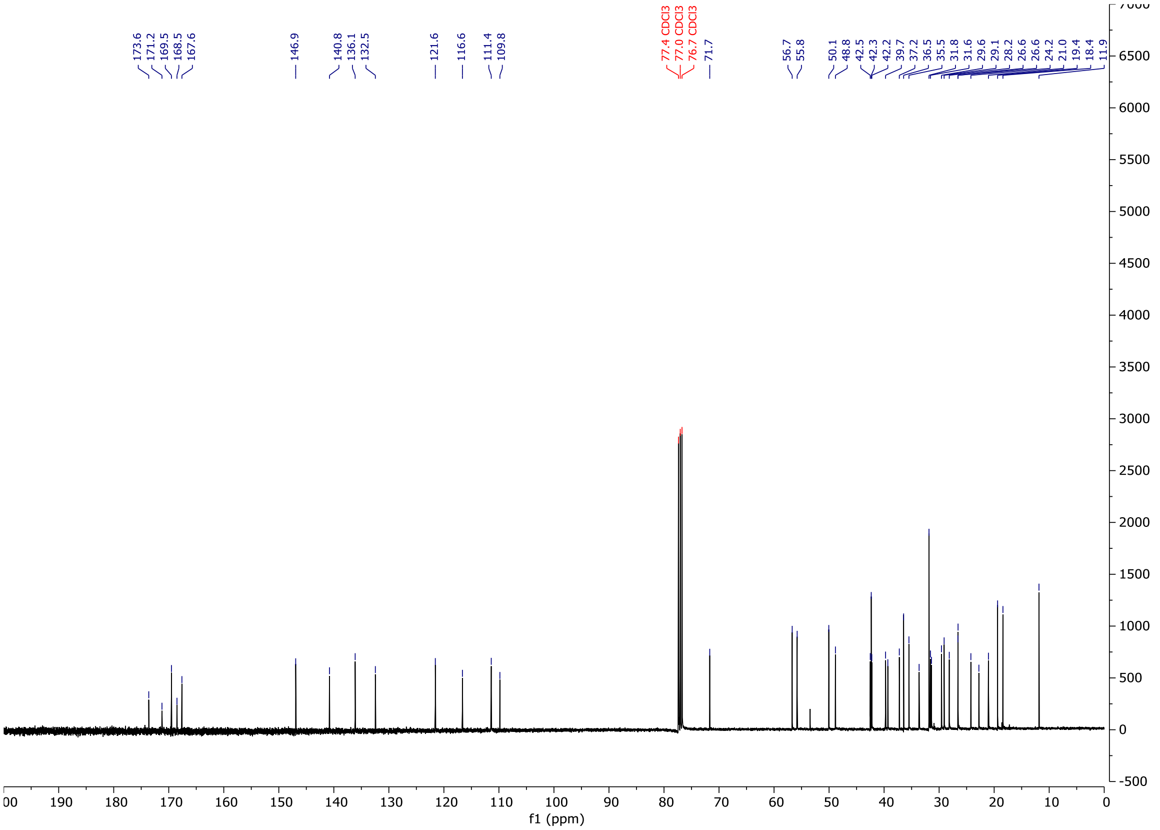
173.6 171.2 169.5 168.5 167.6 146.9 140.8 136.1 132.5 121.6 116.6 111.4 109.8
77.4 CDCl3 77.0 CDCl3 76.7 CDCl3
71.7 56.7 55.8 50.1 48.8 42.5 42.3 42.2 39.7 37.2 36.5 35.5 31.8 31.6 29.6 29.1 28.2 26.6 26.6 24.2 21.0 19.4 18.4 11.9
f1 (ppm)

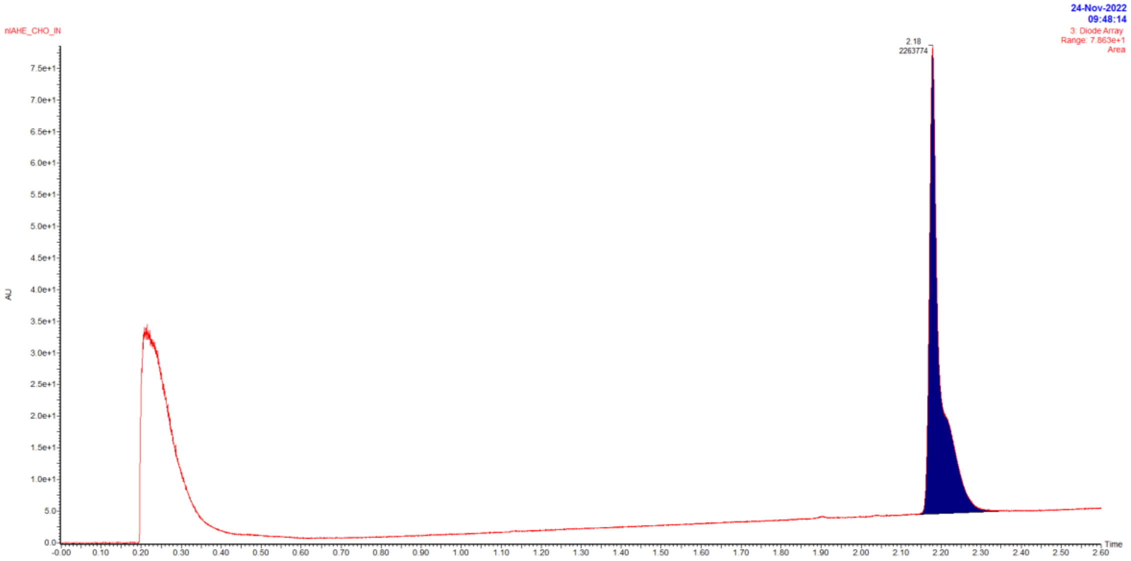


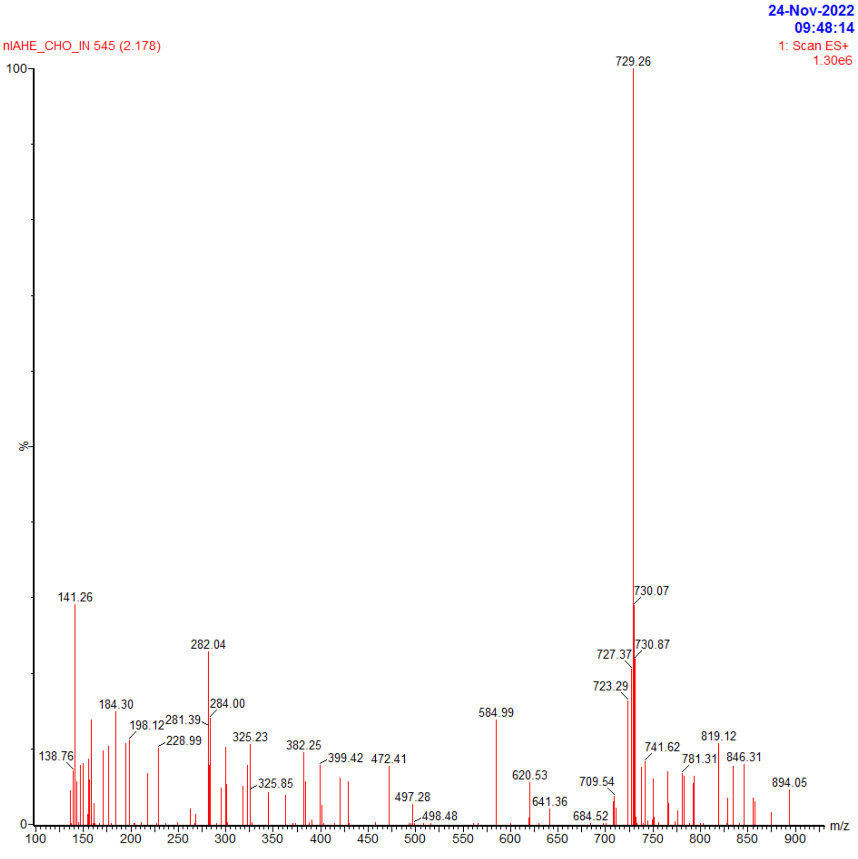


| Retention time | Area | Rel. Area(%) | Found (m/z) |
|---|---|---|---|
| 2.18 | 2263774 | 100 | 729.26 |

# C2

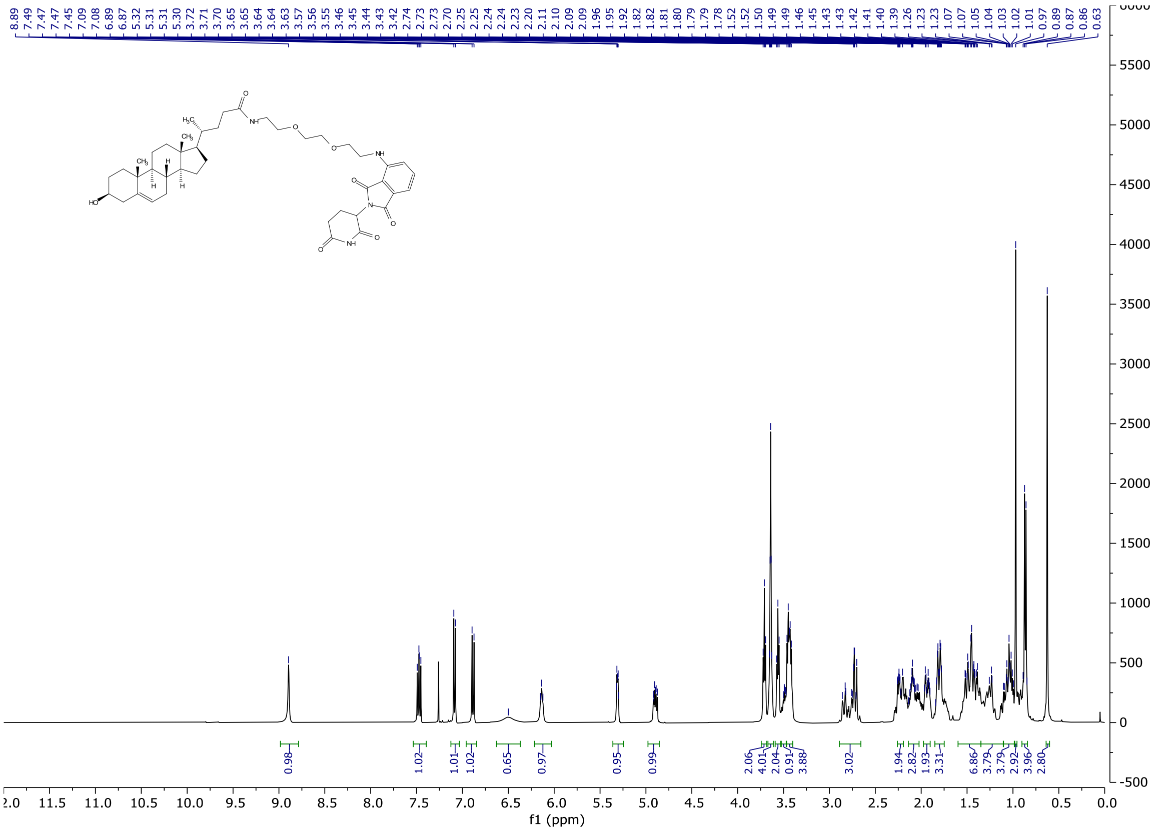
8.89 7.49 7.47 7.47 7.45 7.09 7.08 6.89 6.87 5.32 5.31 5.30 3.72 3.71 3.70 3.65 3.65 3.64 3.63 3.57 3.56 3.55 3.46 3.45 3.44 3.43 3.42 2.74 2.73 2.73 2.70 2.25 2.25 2.24 2.24 2.23 2.20 2.11 2.10 2.09 1.96 1.95 1.92 1.82 1.82 1.81 1.80 1.79 1.78 1.52 1.52 1.50 1.49 1.49 1.46 1.45 1.43 1.43 1.42 1.41 1.40 1.39 1.26 1.23 1.23 1.07 1.07 1.05 1.04 1.03 1.02 1.01 0.97 0.89 0.87 0.86 0.63
0.98 1.02 1.01 1.02 0.65 0.97 0.95 0.99 2.06 4.01 2.04 0.91 3.88 3.02 1.94 2.82 1.93 3.31 6.86 3.79 3.79 2.92 3.96 2.80
f1 (ppm)

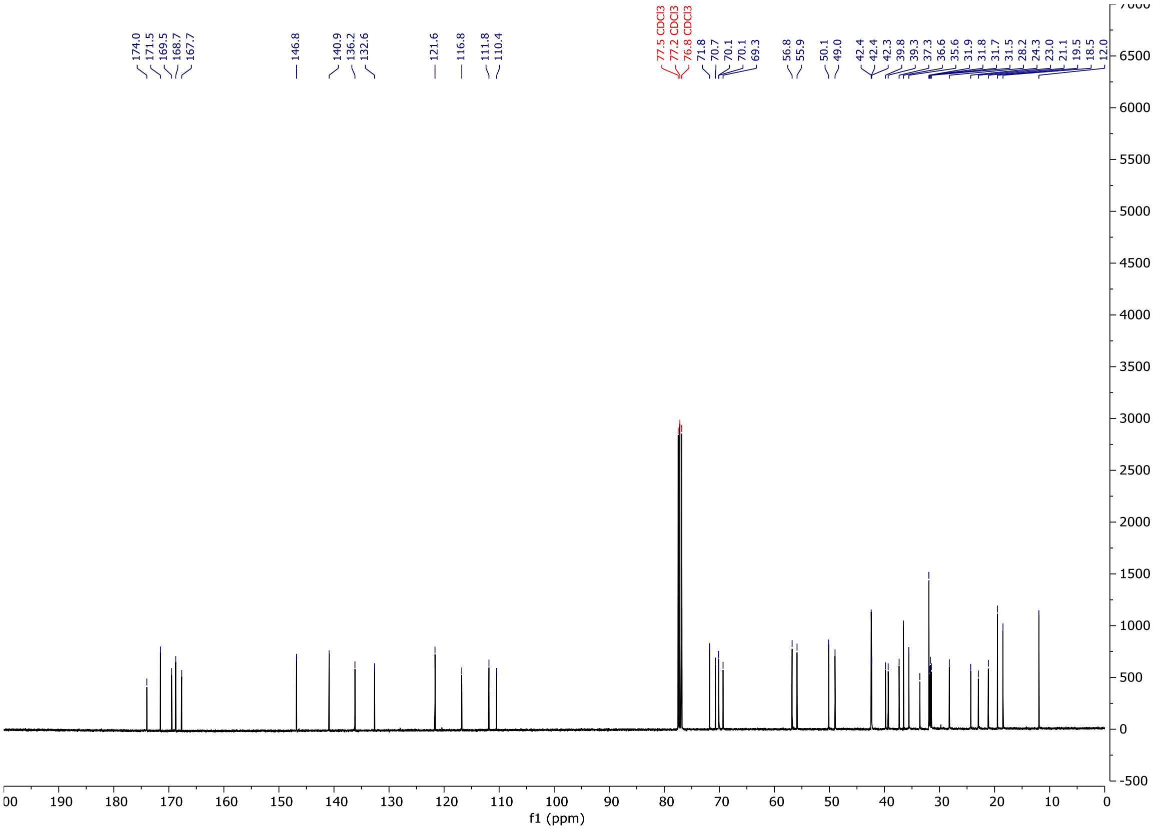
174.0 171.5 169.5 168.7 167.7 146.8 140.9 136.2 132.6 121.6 116.8 111.8 110.4
77.5 CDCl3
77.2 CDCl3
76.8 CDCl3
71.8 70.7 70.1 70.1 69.3 56.8 55.9 50.1 49.0 42.4 42.4 42.3 39.8 39.3 37.3 36.6 35.6 31.9 31.8 31.7 31.5 28.2 24.3 23.0 21.1 19.5 18.5 12.0
f1 (ppm)

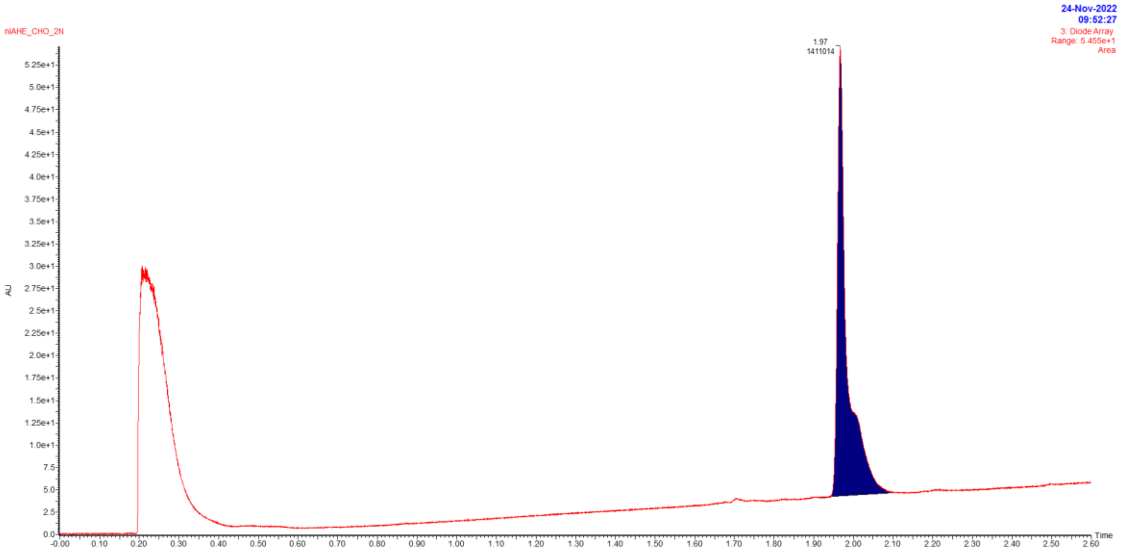


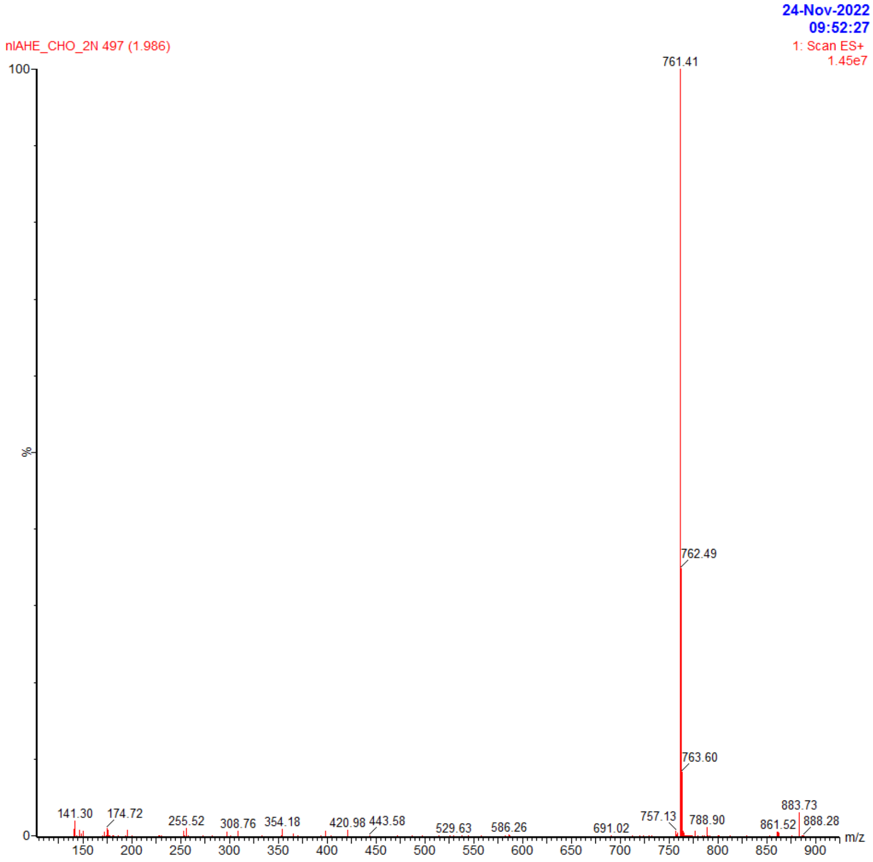


| Retention time | Area | Rel. Area(%) | Found (m/z) |
|---|---|---|---|
| 1.97 | 1411014 | 100 | 761.41 |

**C3**

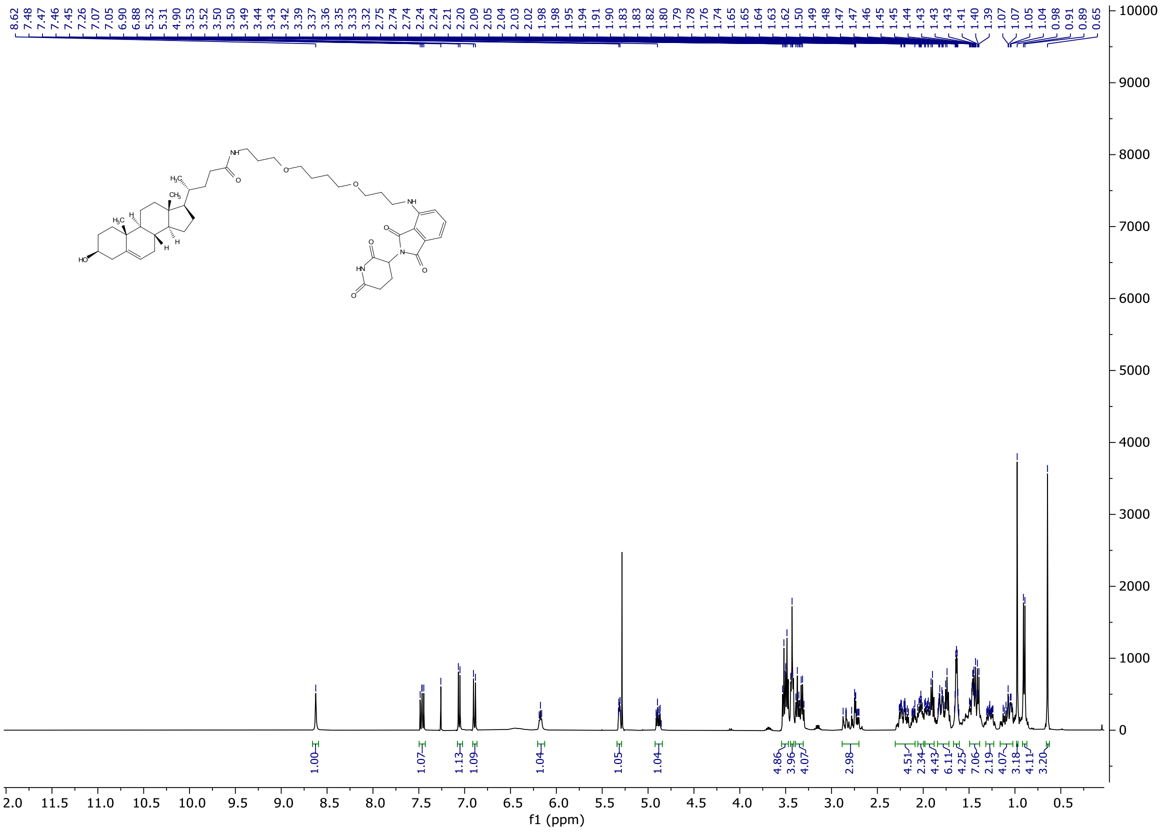

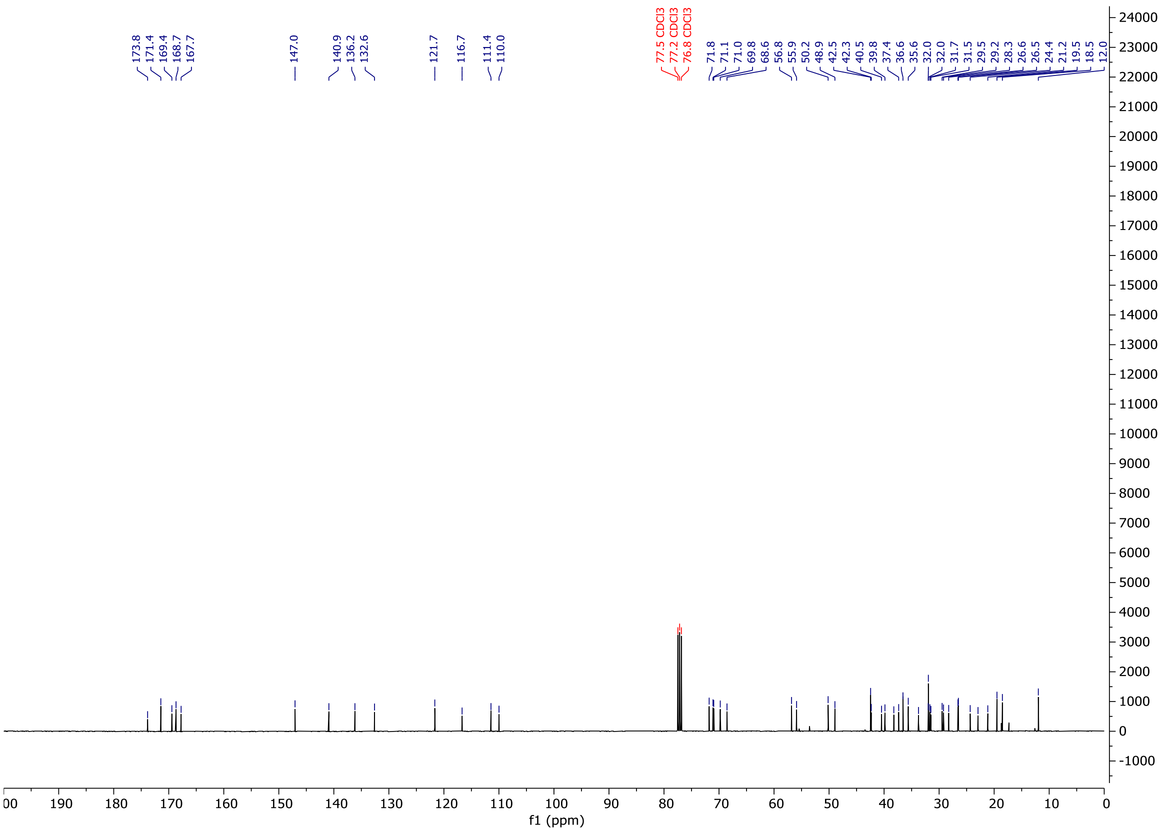

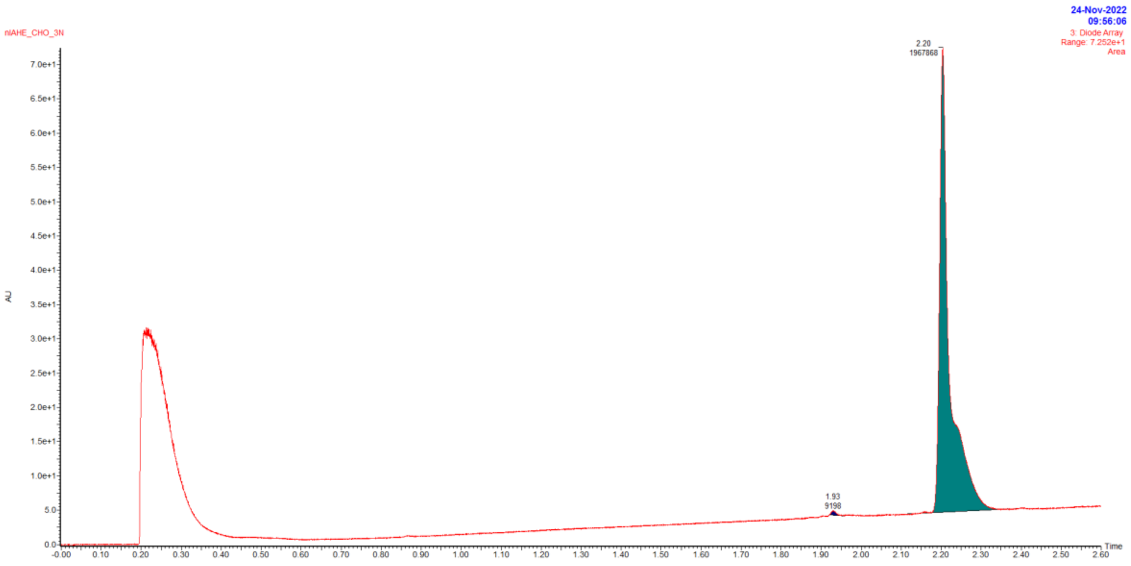


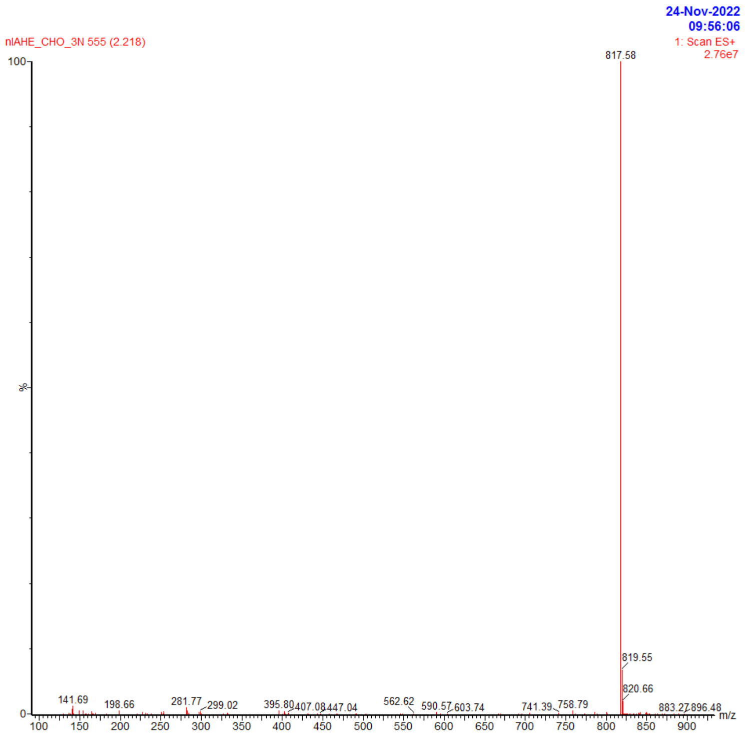


| Retention time | Area | Rel. Area(%) | Found (m/z) |
|---|---|---|---|
| 1.93 | 9198 | <1 | |
| 2.2 | 1967868 | >99 | 817.58 |

# C3_Me

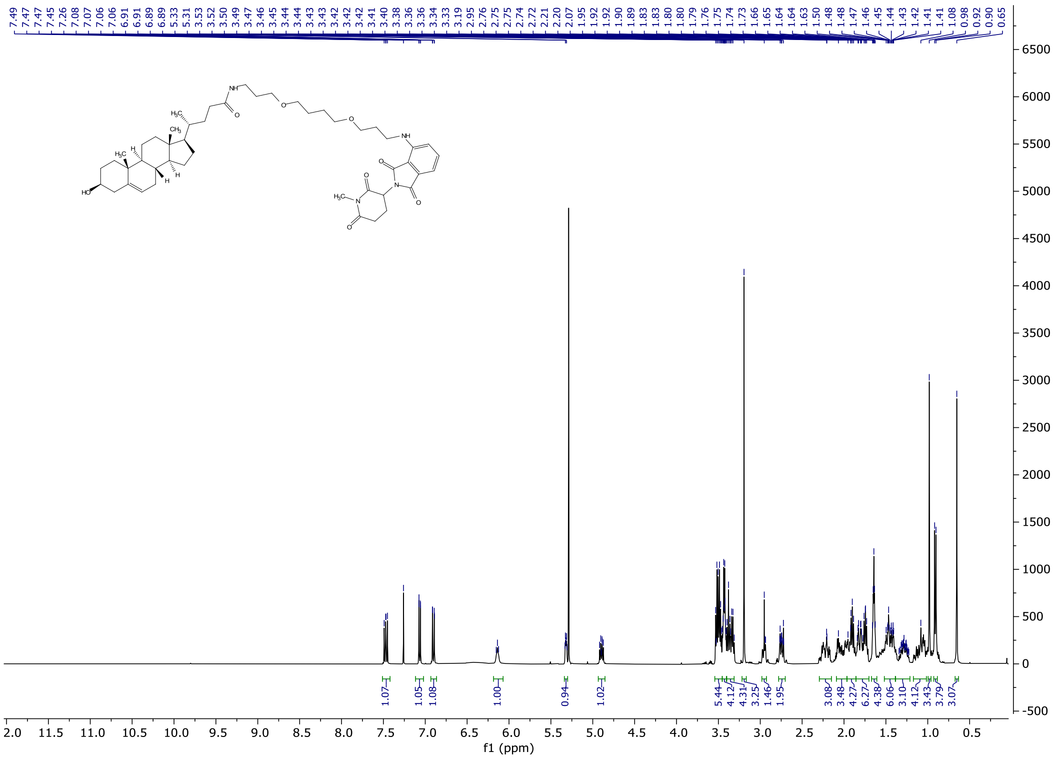

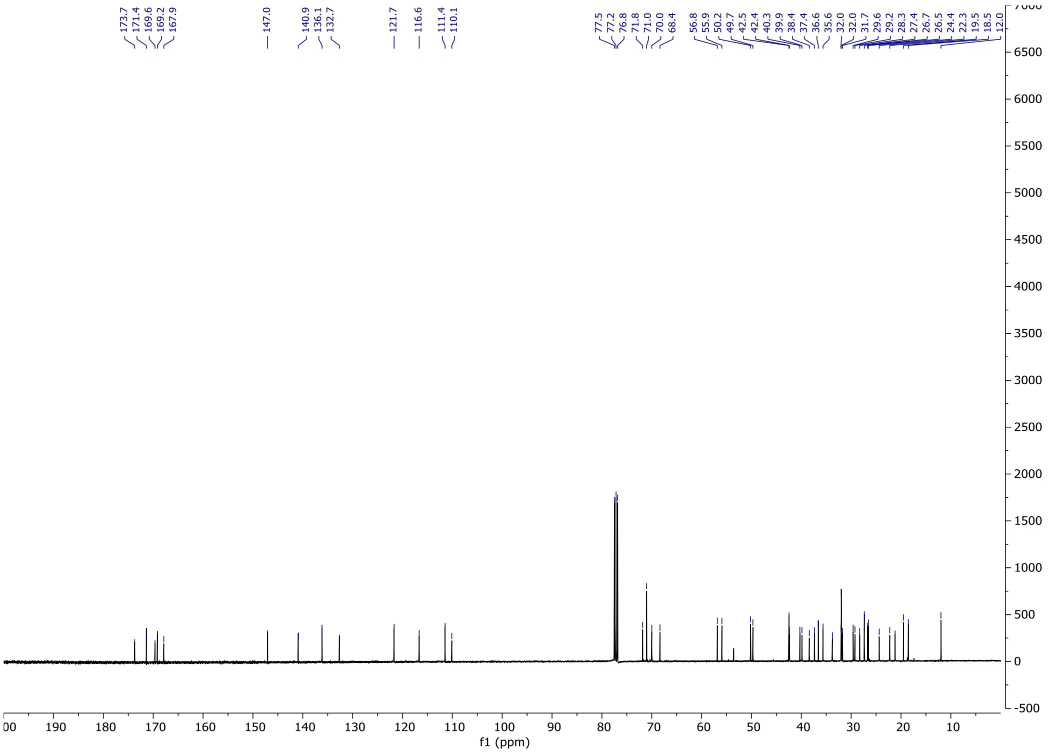

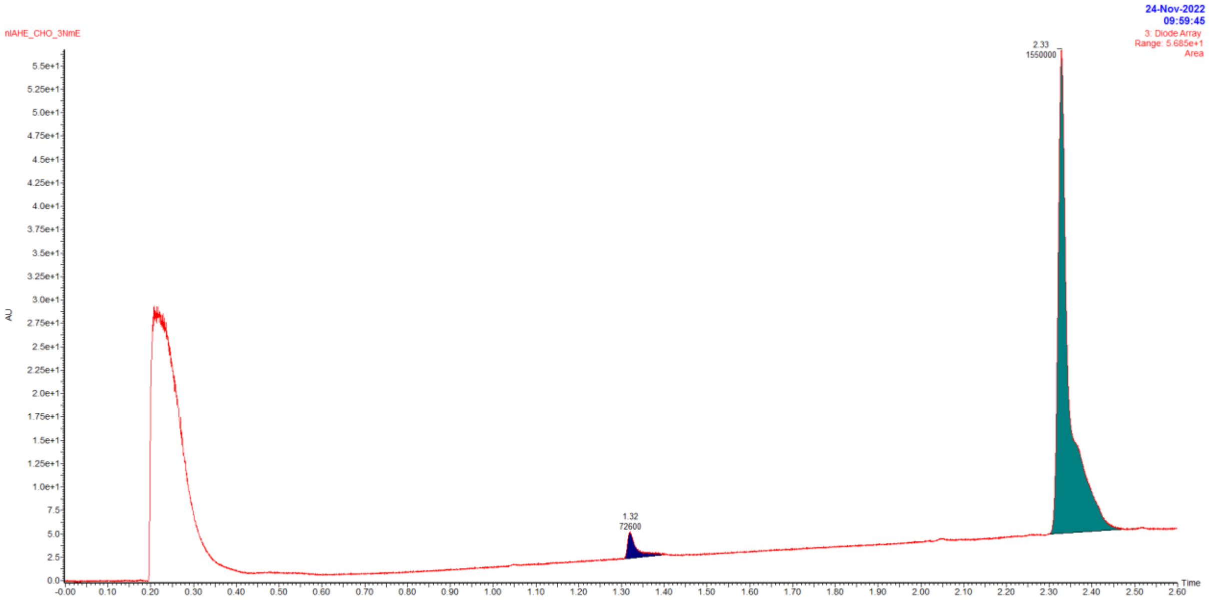


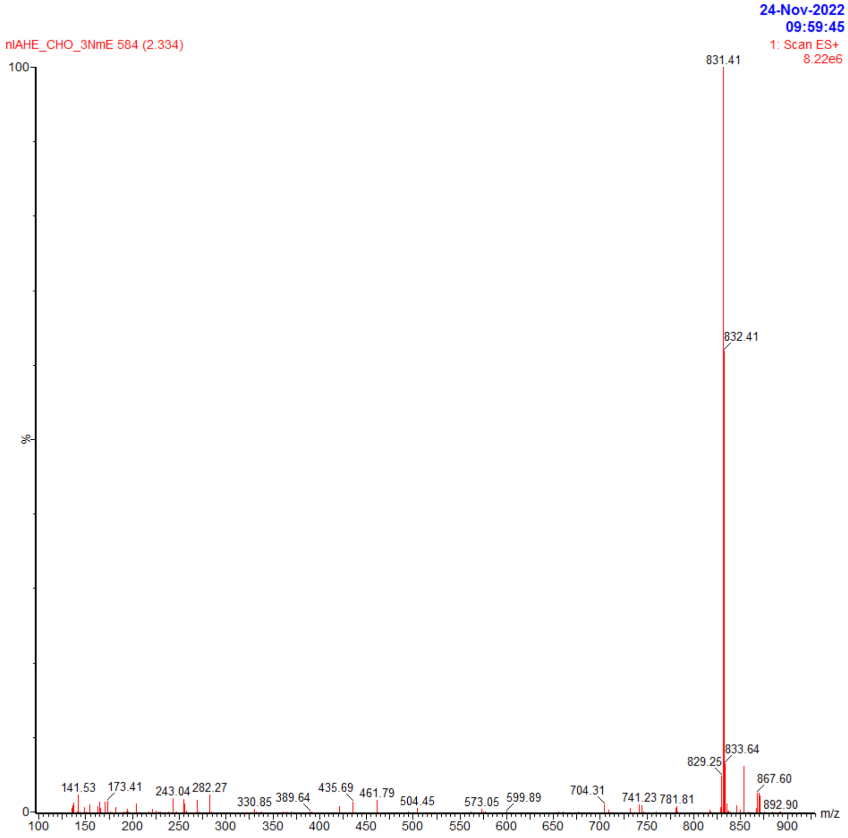


| Retention time | Area | Rel. Area(%) | Found (m/z) |
|---|---|---|---|
| 1.32 | 72600 | 4.5 | |
| 2.33 | 1550000 | 95.5 | 831.41 |

# C4

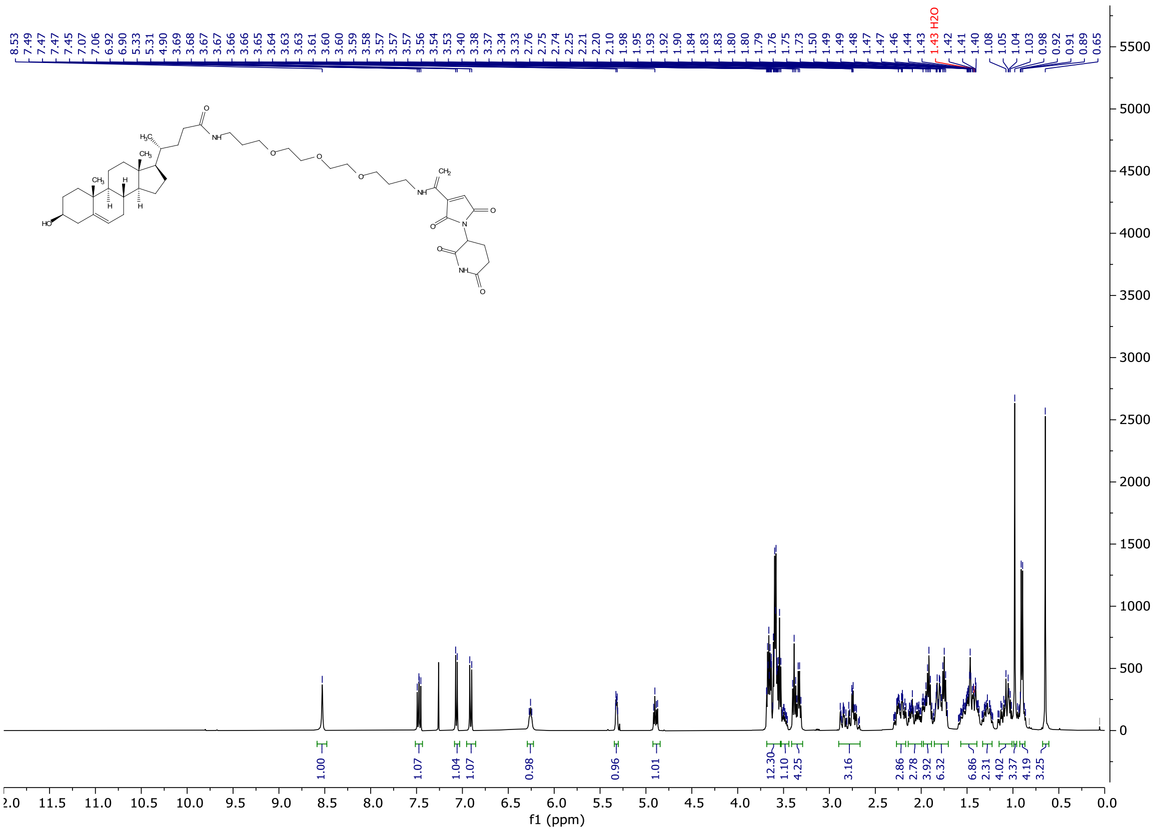
8.53 7.49 7.47 7.47 7.45 7.07 7.06 6.92 6.90 5.33 5.31 4.90 3.69 3.68 3.67 3.67 3.66 3.65 3.64 3.63 3.63 3.61 3.60 3.60 3.59 3.58 3.57 3.57 3.56 3.54 3.53 3.40 3.38 3.37 3.34 3.33 2.76 2.75 2.74 2.25 2.21 2.20 2.10 1.98 1.95 1.93 1.92 1.90 1.84 1.83 1.83 1.80 1.79 1.76 1.75 1.73 1.50 1.49 1.48 1.47 1.47 1.46 1.44 1.43 1.43 H2O 1.42 1.41 1.40 1.08 1.05 1.04 1.03 0.98 0.92 0.91 0.89 0.65
1.00 1.07 1.04 1.07 0.98 0.96 1.01 12.30 1.10 4.25 3.16 2.86 2.78 3.92 6.32 6.86 2.31 4.02 3.37 4.19 3.25
f1 (ppm)

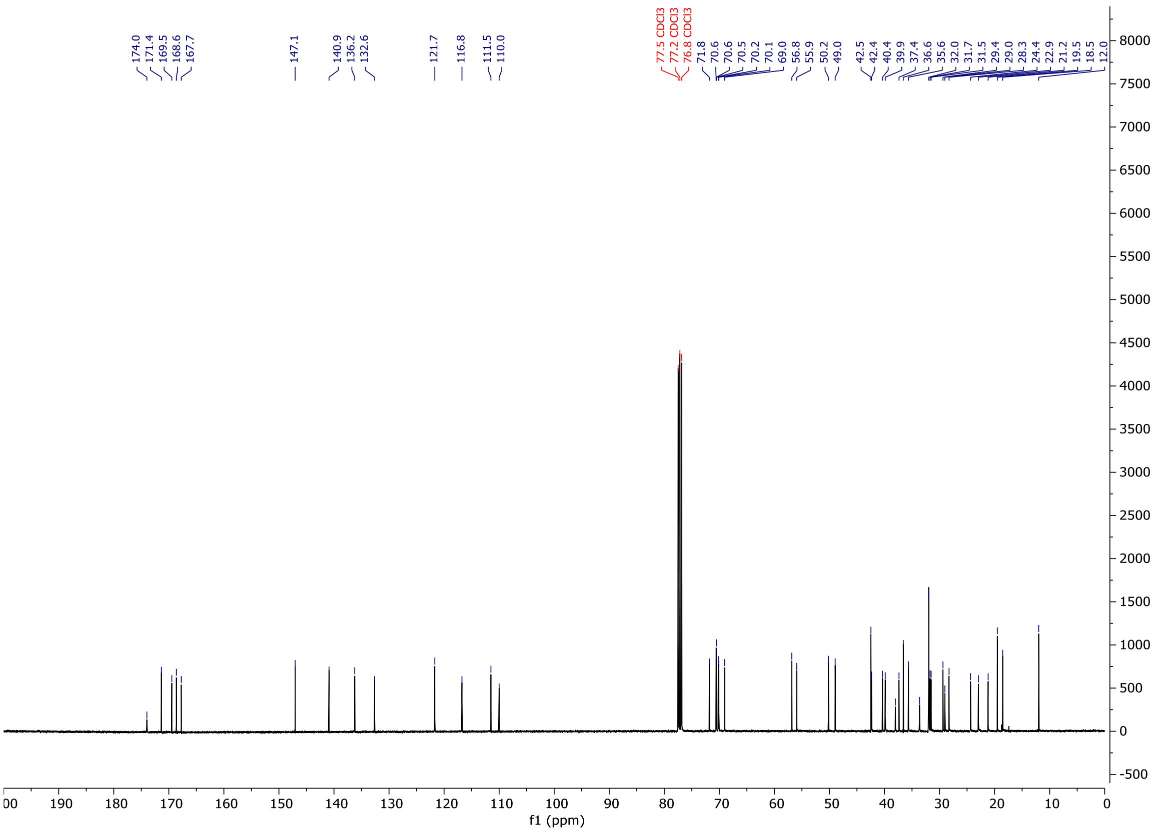
174.0 171.4 169.5 168.6 167.7 147.1 140.9 136.2 132.6 121.7 116.8 111.5 110.0 77.5 CDCl3 77.2 CDCl3 76.8 CDCl3 71.8 70.6 70.6 70.5 70.2 70.1 69.0 56.8 55.9 50.2 49.0 42.5 42.4 40.4 39.9 37.4 36.6 35.6 32.0 31.7 31.5 29.4 29.0 28.3 24.4 22.9 21.2 19.5 18.5 12.0
f1 (ppm)

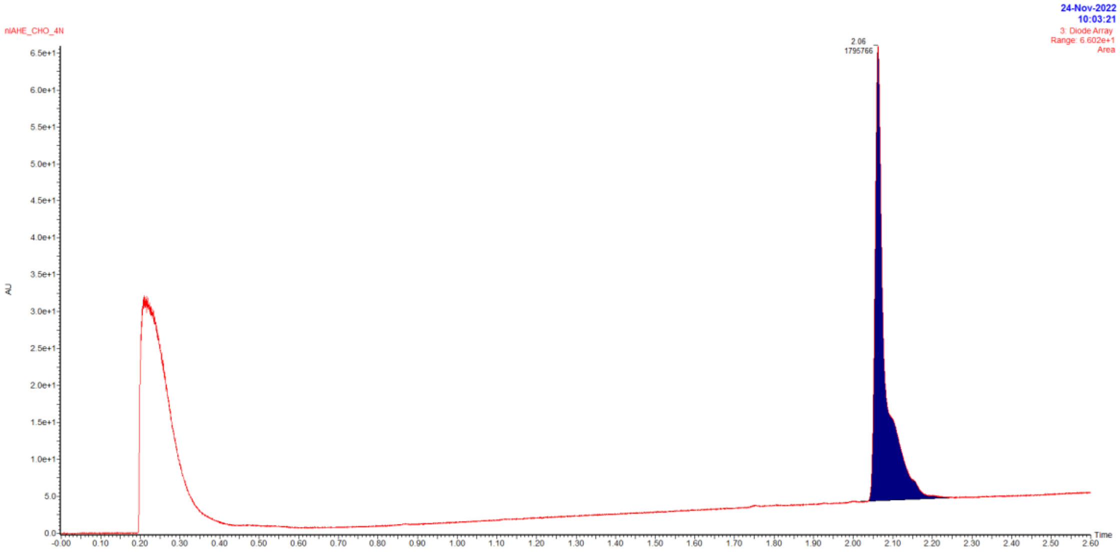


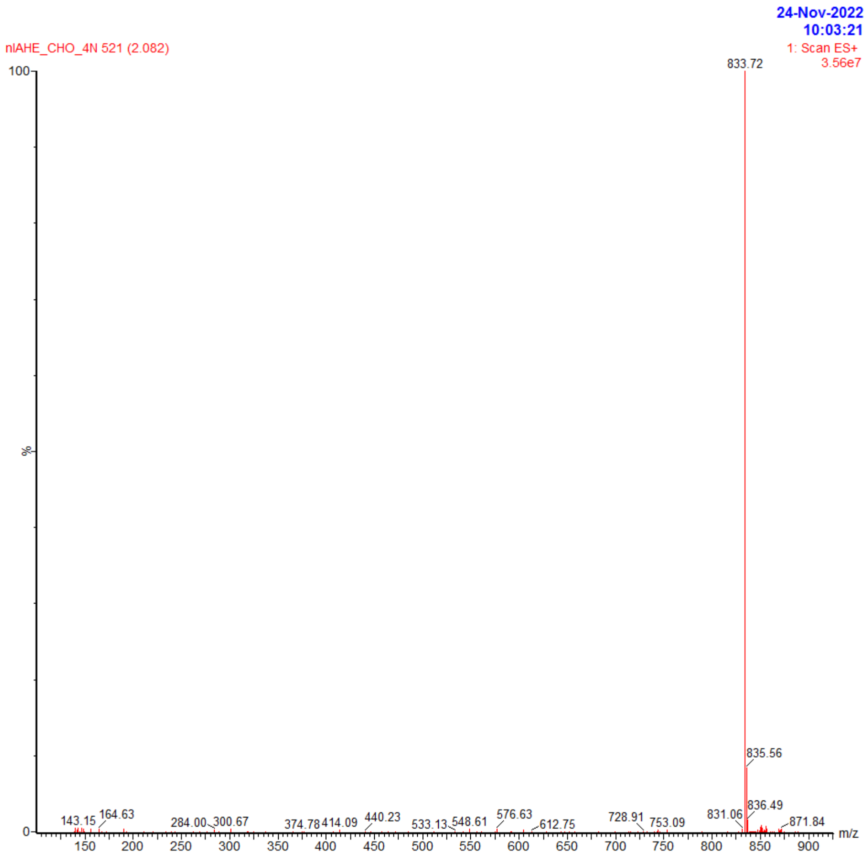


| Retention time | Area | Rel. Area(%) | Found (m/z) |
|---|---|---|---|
| 2.06 | 1795766 | 100 | 833.72 |

# C5

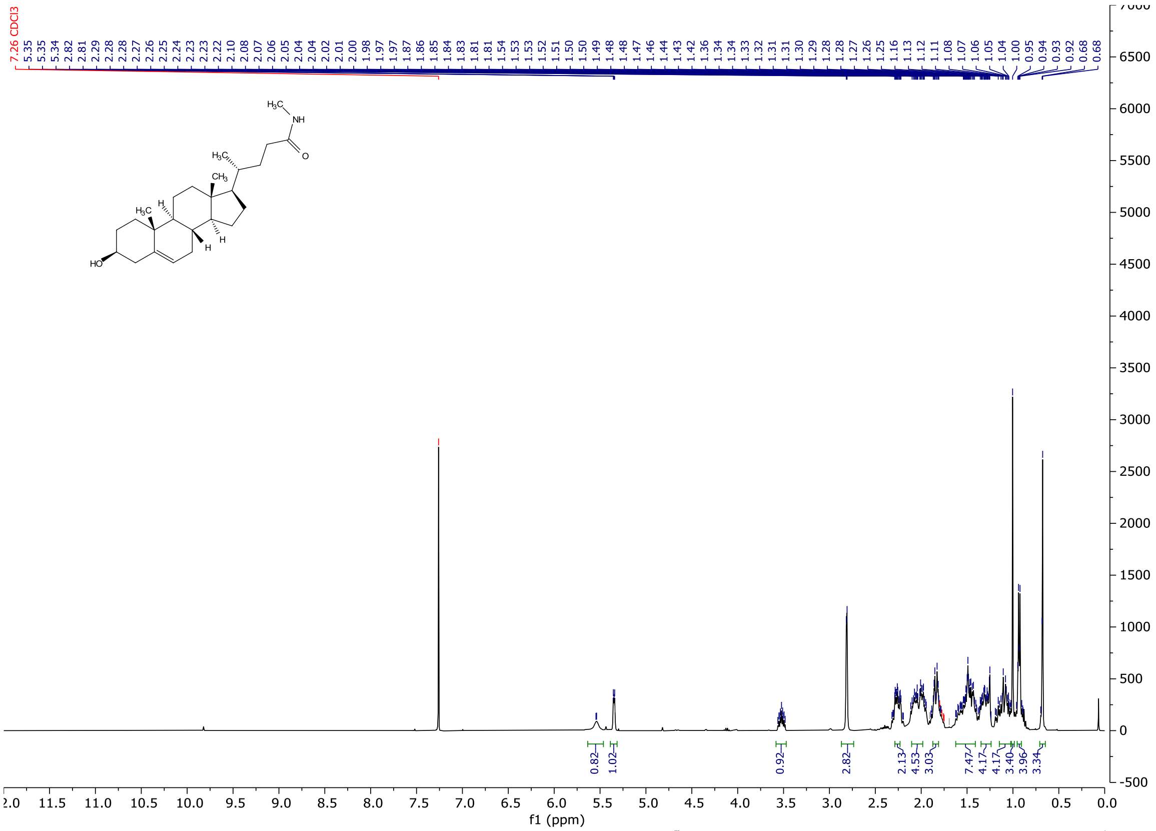
7.26 CDCl3
5.35 5.35 5.34 2.82 2.81 2.29 2.28 2.28 2.27 2.26 2.25 2.24 2.23 2.23 2.22 2.10 2.08 2.07 2.06 2.05 2.04 2.04 2.02 2.01 2.00 1.98 1.97 1.97 1.87 1.86 1.85 1.84 1.83 1.81 1.81 1.54 1.53 1.53 1.52 1.51 1.50 1.49 1.48 1.47 1.46 1.44 1.43 1.42 1.36 1.34 1.34 1.33 1.32 1.31 1.31 1.30 1.29 1.28 1.28 1.27 1.26 1.25 1.16 1.13 1.12 1.11 1.08 1.07 1.06 1.05 1.04 1.00 0.95 0.94 0.93 0.92 0.68 0.68
H3C
NH
O
H3C
CH3
H
H3C
H
H
HO
0.82 1.02 0.92 2.82 2.13 4.53 3.03 7.47 4.17 4.17 3.40 3.96 3.34
f1 (ppm)

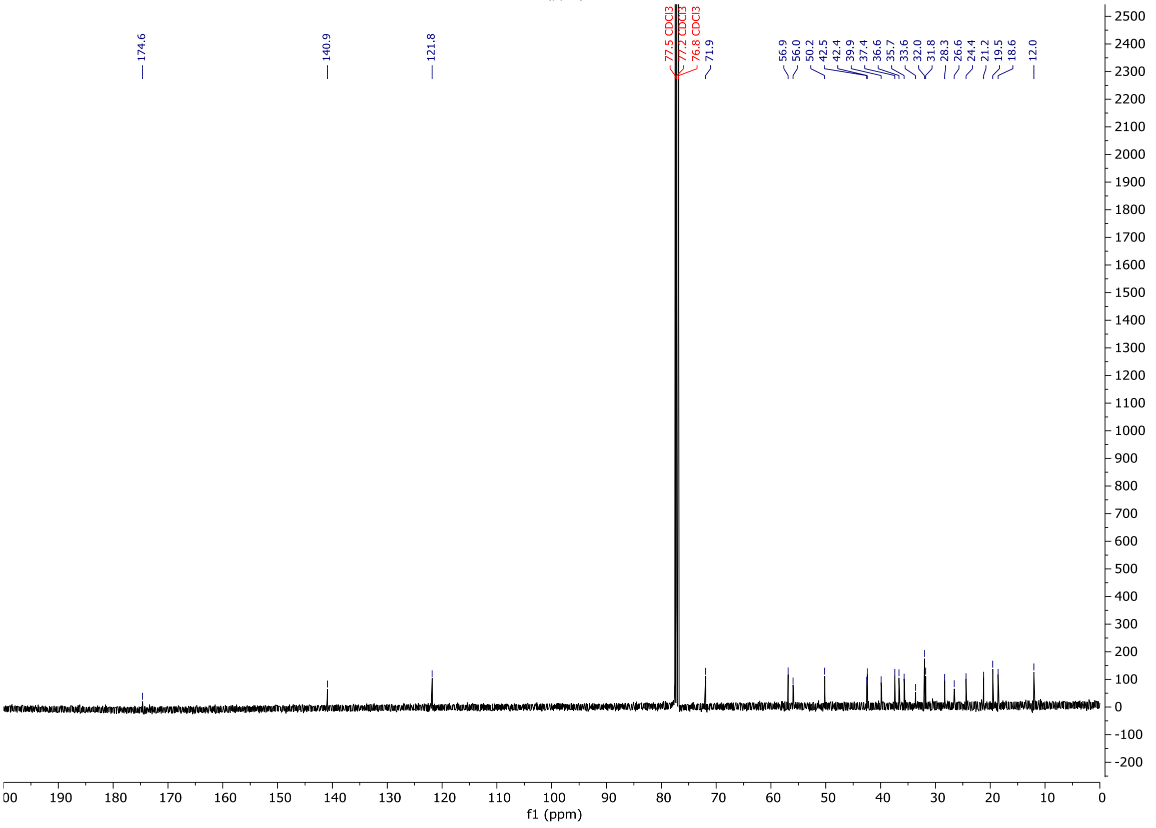
174.6
140.9
121.8
77.5 CDCl3
77.2 CDCl3
76.8 CDCl3
71.9
56.9 56.0 50.2 42.5 42.4 39.9 37.4 36.6 35.7 33.6 32.0 31.8 28.3 26.6 24.4 21.2 19.5 18.6 12.0
f1 (ppm)

### C3_Boc (Oxybipin-2)

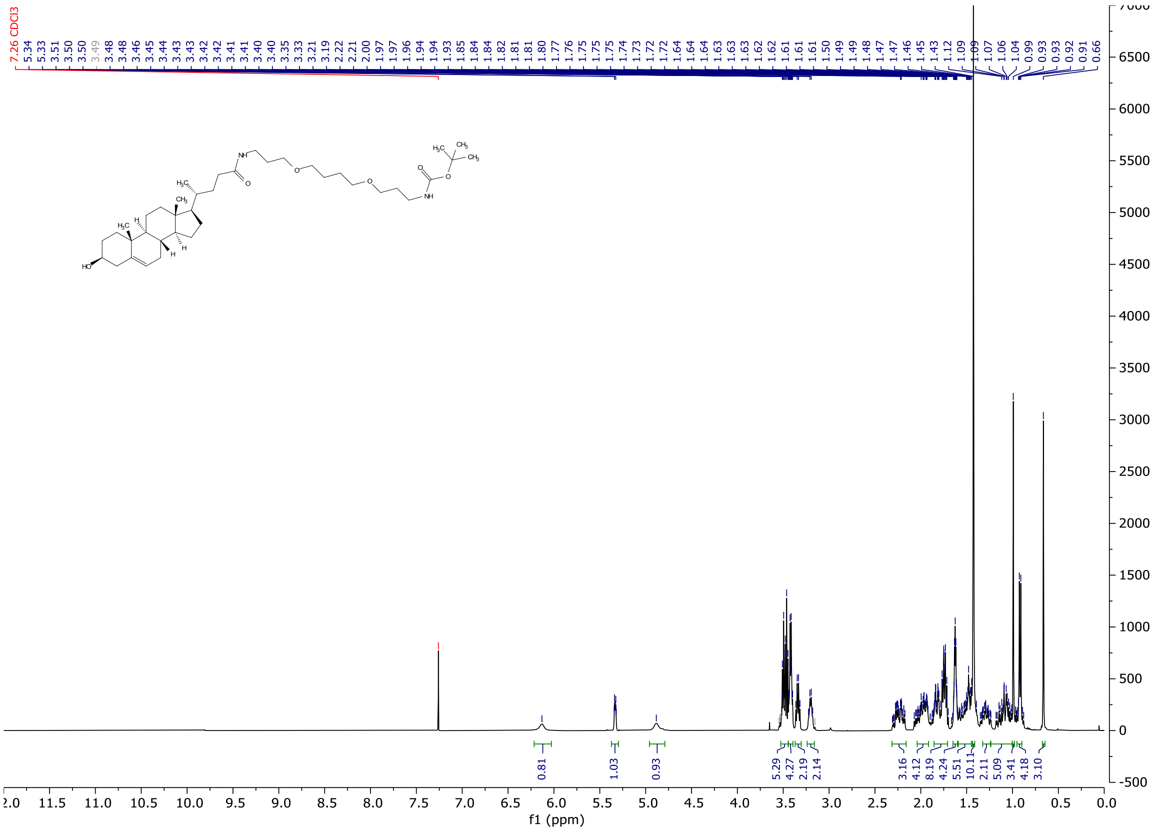


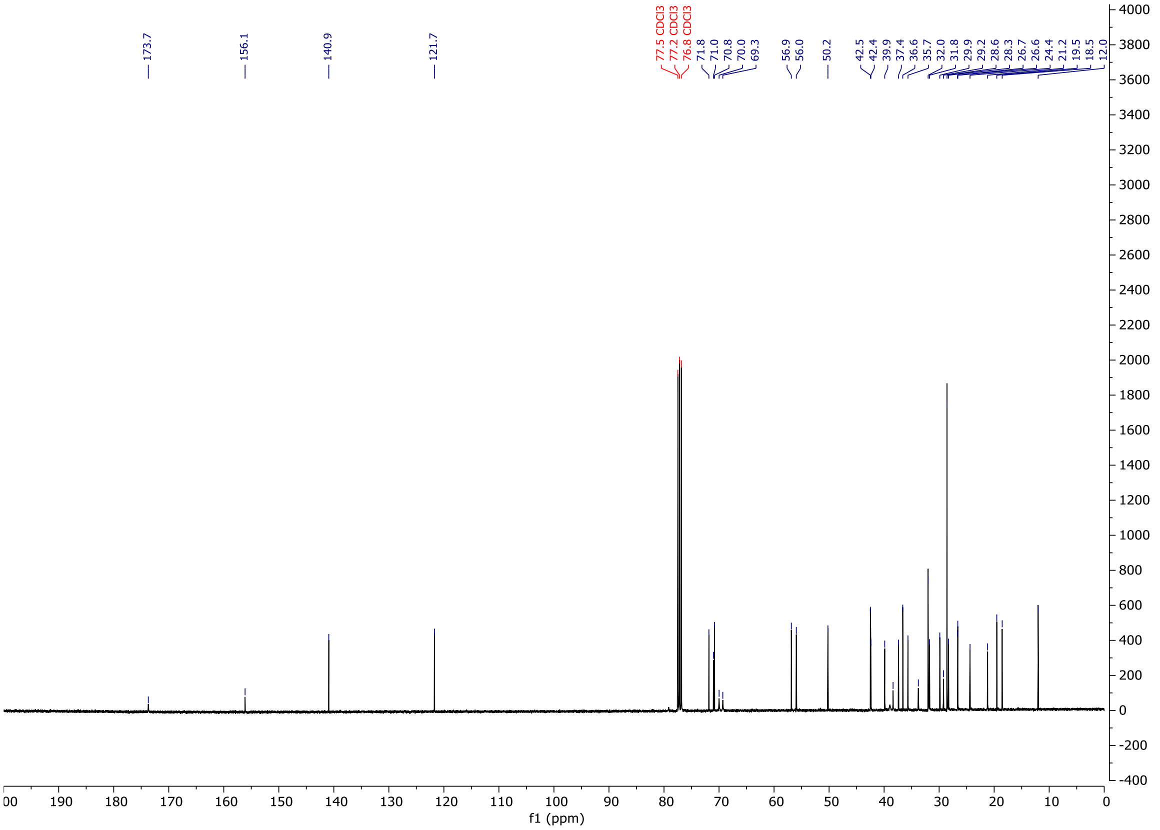

### C3_HyT (Oxybipin-1)

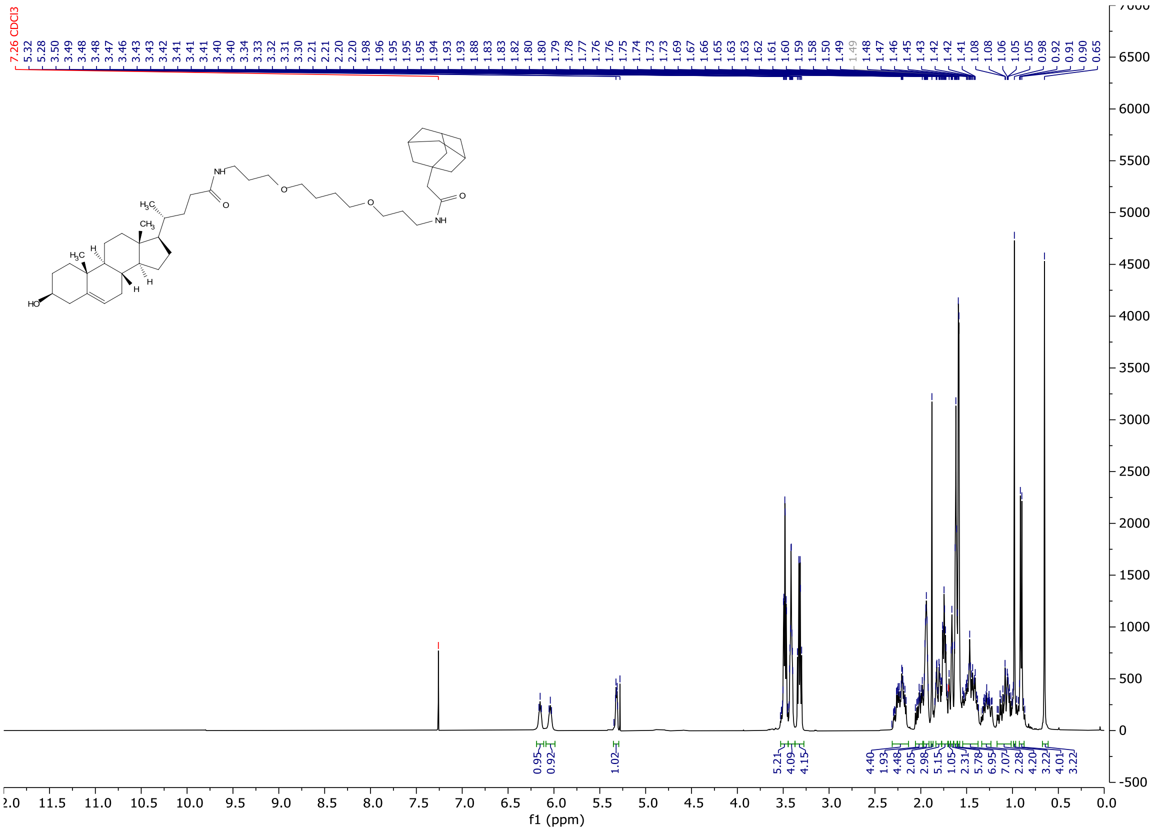


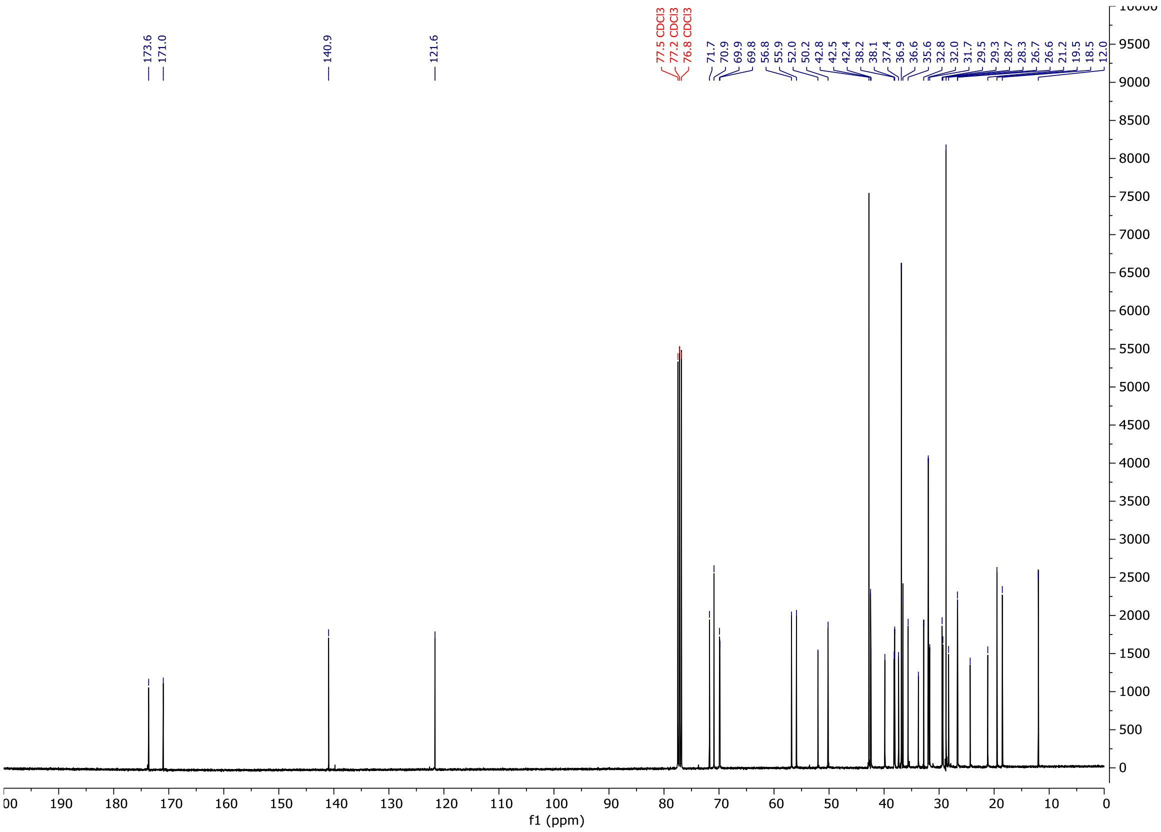

### ORD_027

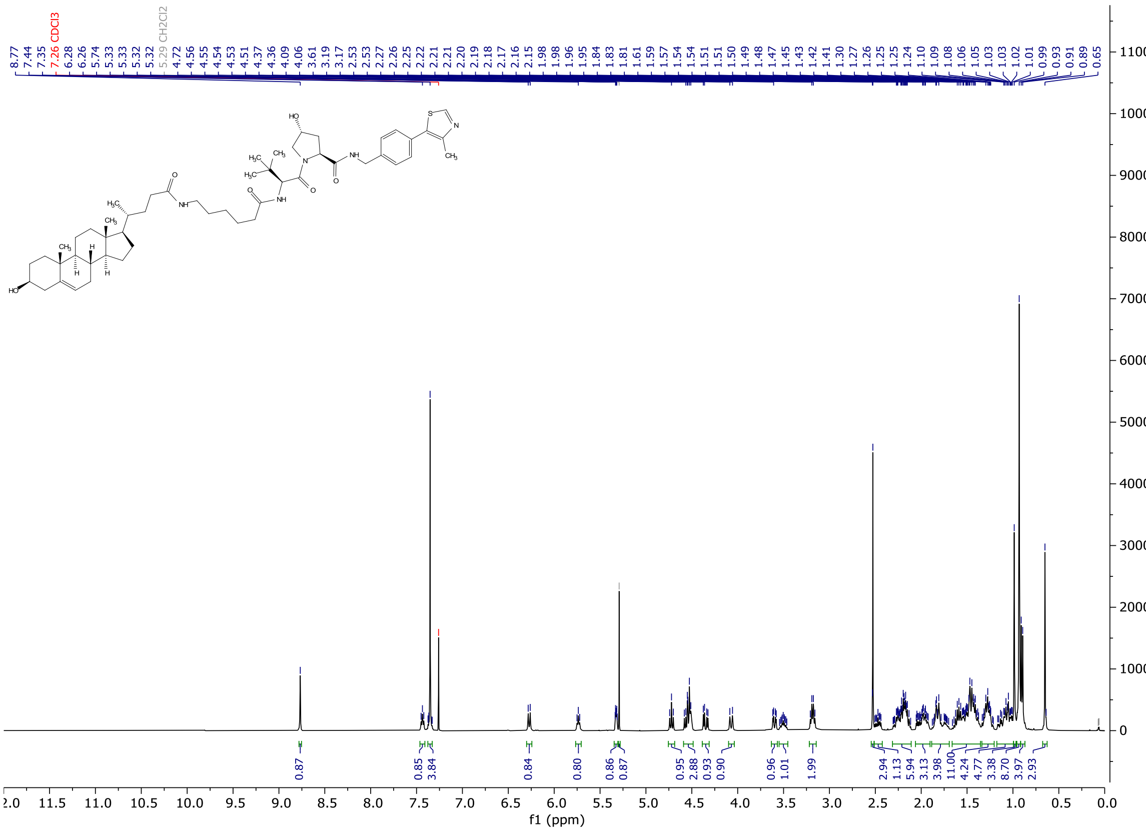

8.77
7.44
7.35
7.26 CDCl3
6.28
6.26
5.74
5.33
5.33
5.32
5.29 CH2Cl2
4.72
4.56
4.55
4.54
4.53
4.51
4.37
4.36
4.09
4.06
3.61
3.19
3.17
2.53
2.53
2.27
2.26
2.25
2.22
2.21
2.21
2.20
2.19
2.18
2.17
2.16
2.15
1.98
1.98
1.96
1.95
1.84
1.83
1.83
1.81
1.61
1.59
1.57
1.54
1.54
1.51
1.51
1.50
1.49
1.48
1.47
1.45
1.43
1.42
1.41
1.30
1.27
1.26
1.25
1.25
1.24
1.10
1.09
1.08
1.06
1.05
1.03
1.03
1.02
1.01
0.99
0.93
0.91
0.89
0.65
0.87
0.85
3.84
0.84
0.80
0.86
0.87
0.95
2.88
0.93
0.90
0.96
1.01
1.99
2.94
1.13
5.94
3.13
3.98
11.00
4.24
4.77
3.38
8.70
3.97
2.93
f1 (ppm)


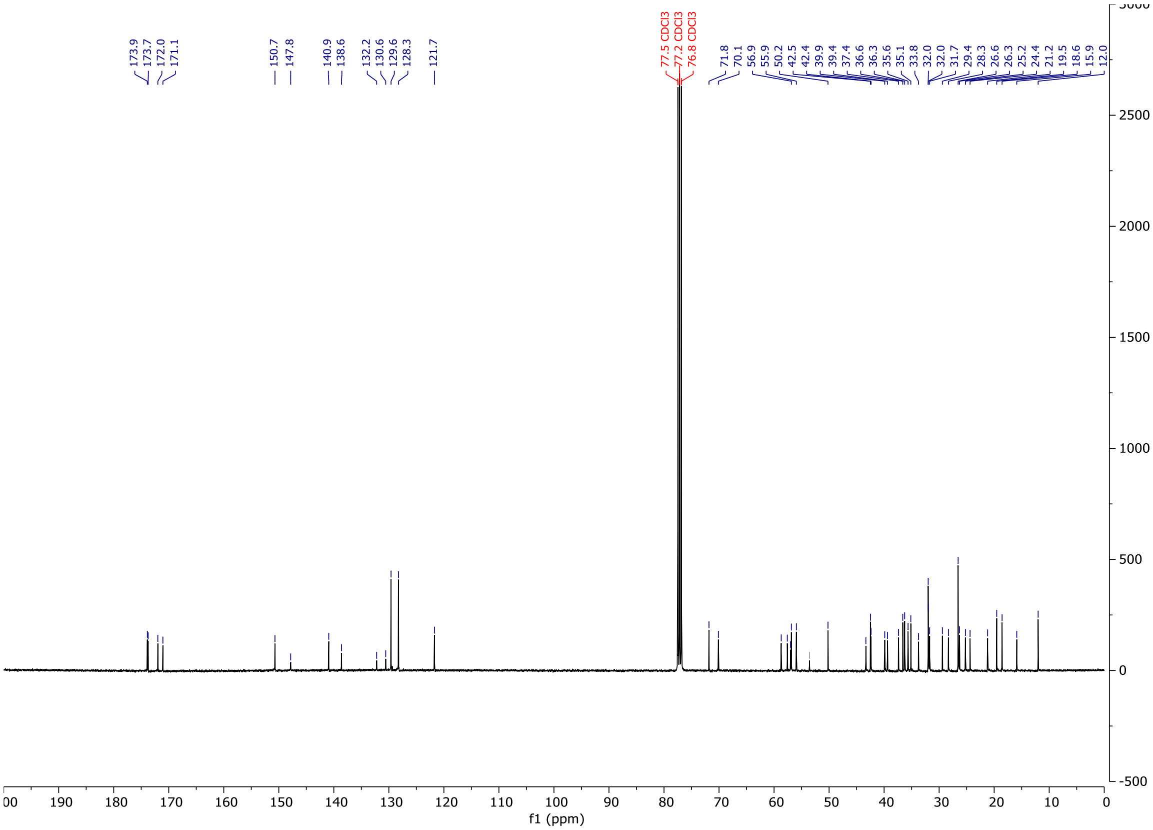

173.9
173.7
172.0
171.1
150.7
147.8
140.9
138.6
132.2
130.6
129.6
128.3
121.7
77.5 CDCl3
77.2 CDCl3
76.8 CDCl3
71.8
70.1
56.9
55.9
50.2
42.5
42.4
39.9
39.4
37.4
36.6
36.3
35.6
35.1
33.8
32.0
32.0
31.7
29.4
28.3
26.6
26.3
25.2
24.4
21.2
19.5
18.6
15.9
12.0
f1 (ppm)

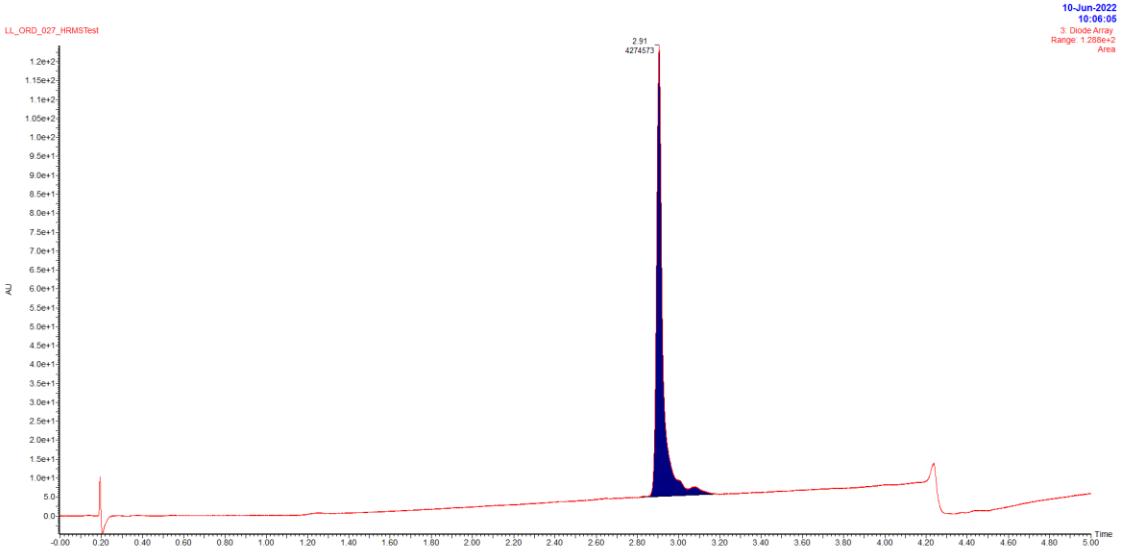


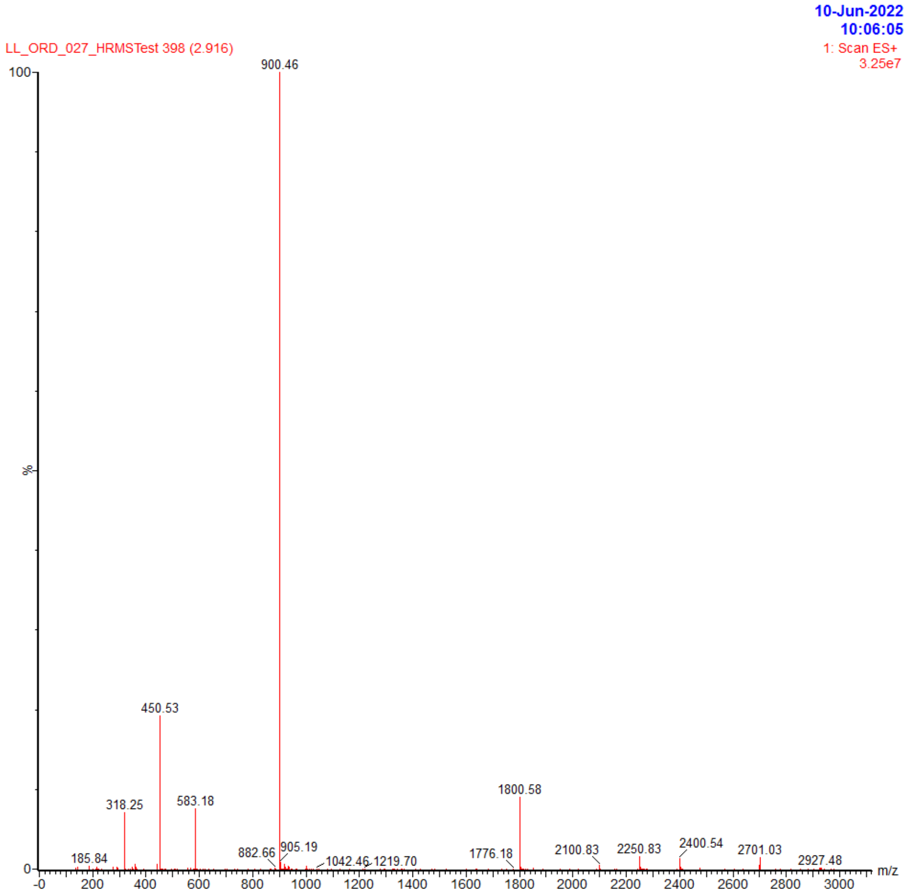


| Retention time | Area | Rel. Area(%) | Found (m/z) |
|---|---|---|---|
| 2.91 | 4274573 | 100 | 900.46 |

### ORD_031

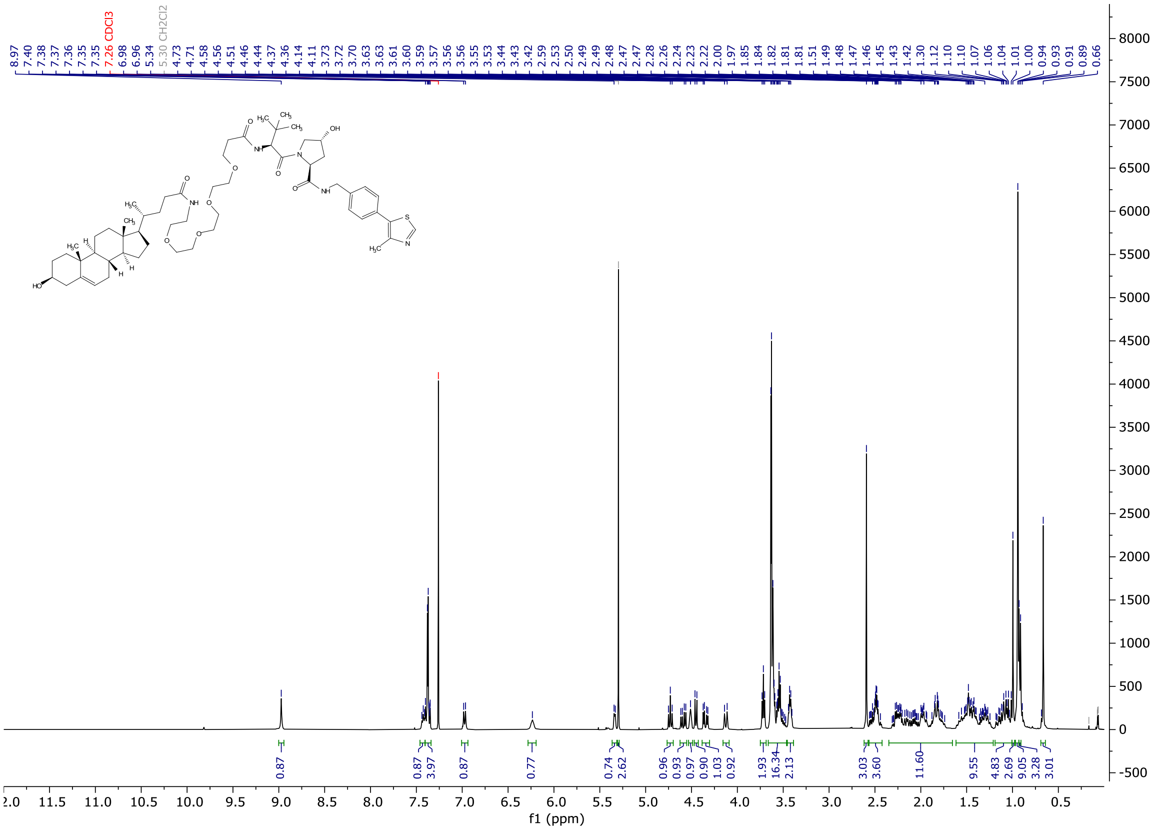
8.97 7.40 7.38 7.37 7.36 7.35 7.35 7.26 CDCl3 6.98 6.96 5.34 5.30 CH2Cl2 4.73 4.71 4.58 4.56 4.51 4.46 4.44 4.37 4.36 4.14 4.11 3.73 3.72 3.70 3.63 3.63 3.61 3.60 3.59 3.57 3.56 3.55 3.53 3.44 3.43 3.42 2.59 2.53 2.50 2.49 2.48 2.47 2.28 2.26 2.24 2.23 2.22 2.00 1.97 1.85 1.84 1.82 1.81 1.81 1.51 1.49 1.48 1.47 1.46 1.45 1.45 1.43 1.42 1.30 1.12 1.10 1.10 1.07 1.06 1.04 1.01 1.00 0.94 0.93 0.91 0.89 0.66
0.87 0.87 3.97 0.87 0.77 0.74 2.62 0.96 0.93 0.97 0.90 1.03 0.92 1.93 16.34 2.13 3.03 3.60 11.60 9.55 4.83 2.69 9.05 3.28 3.01
f1 (ppm)

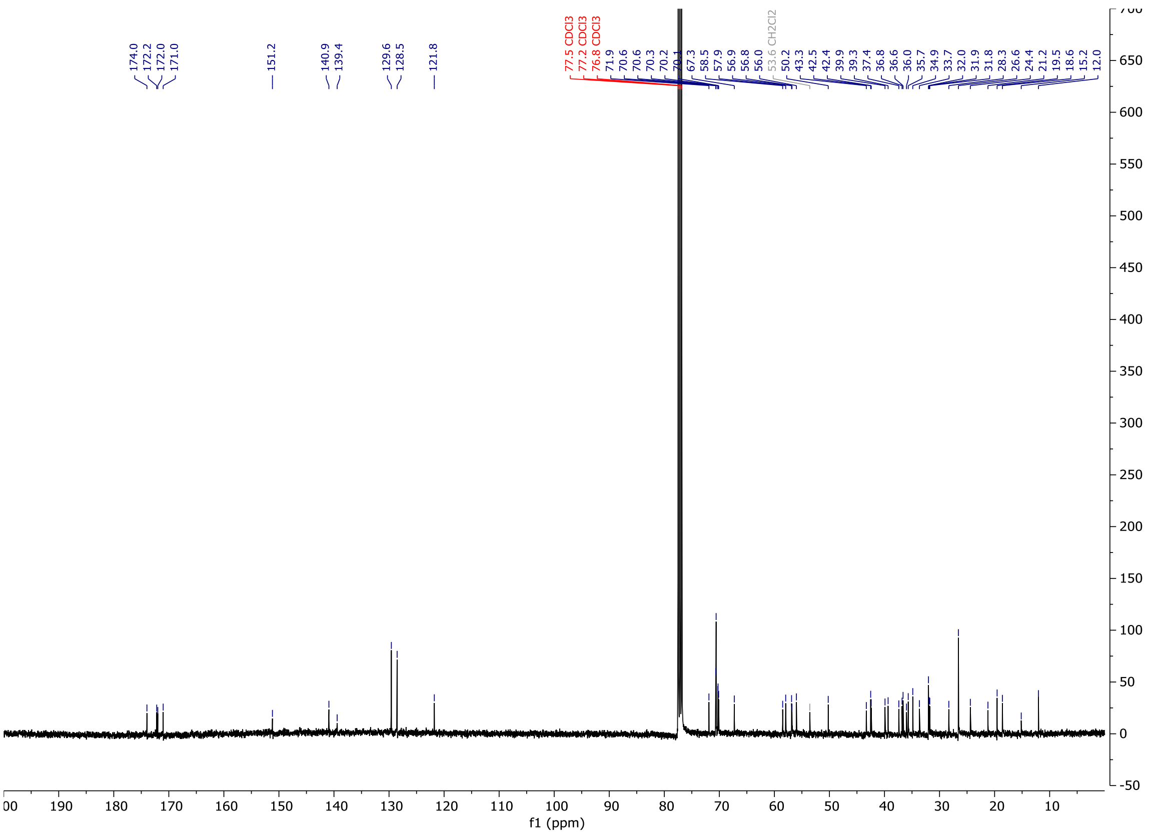
174.0 172.2 172.0 171.0 151.2 140.9 139.4 129.6 128.5 121.8
77.5 CDCl3 77.2 CDCl3 76.8 CDCl3
71.9 70.6 70.6 70.3 70.2 70.1 67.3 58.5 57.9 56.9 56.8 56.0 53.6 CH2Cl2 50.2 43.3 42.5 42.4 39.9 39.3 37.4 36.8 36.6 36.0 35.7 34.9 33.7 32.0 31.9 31.8 28.3 26.6 24.4 21.2 19.5 18.6 15.2 12.0
f1 (ppm)

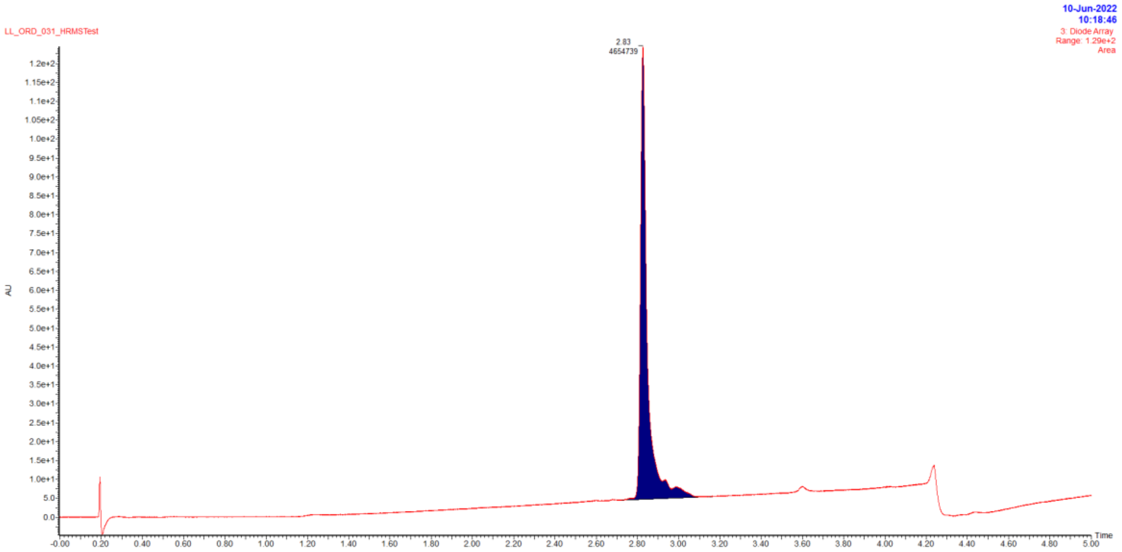


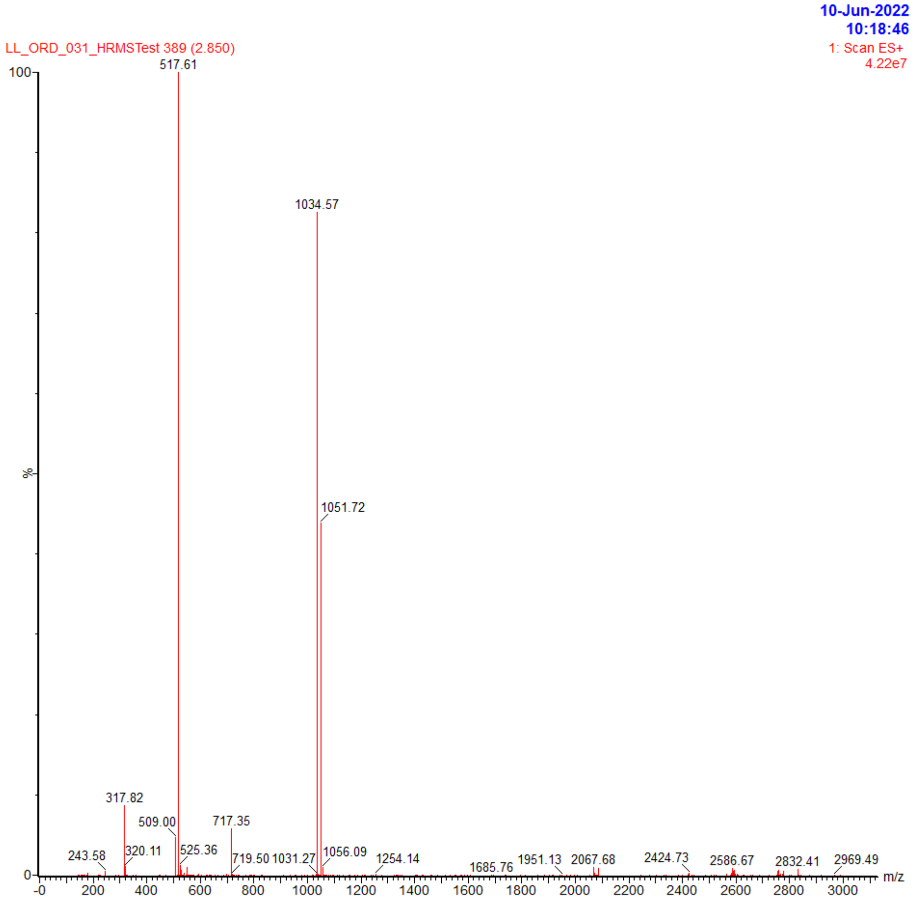


| Retention time | Area | Rel. Area(%) | Found (m/z) |
|---|---|---|---|
| 2.83 | 4654739 | 100 | 1034.57 |

### ORD_033

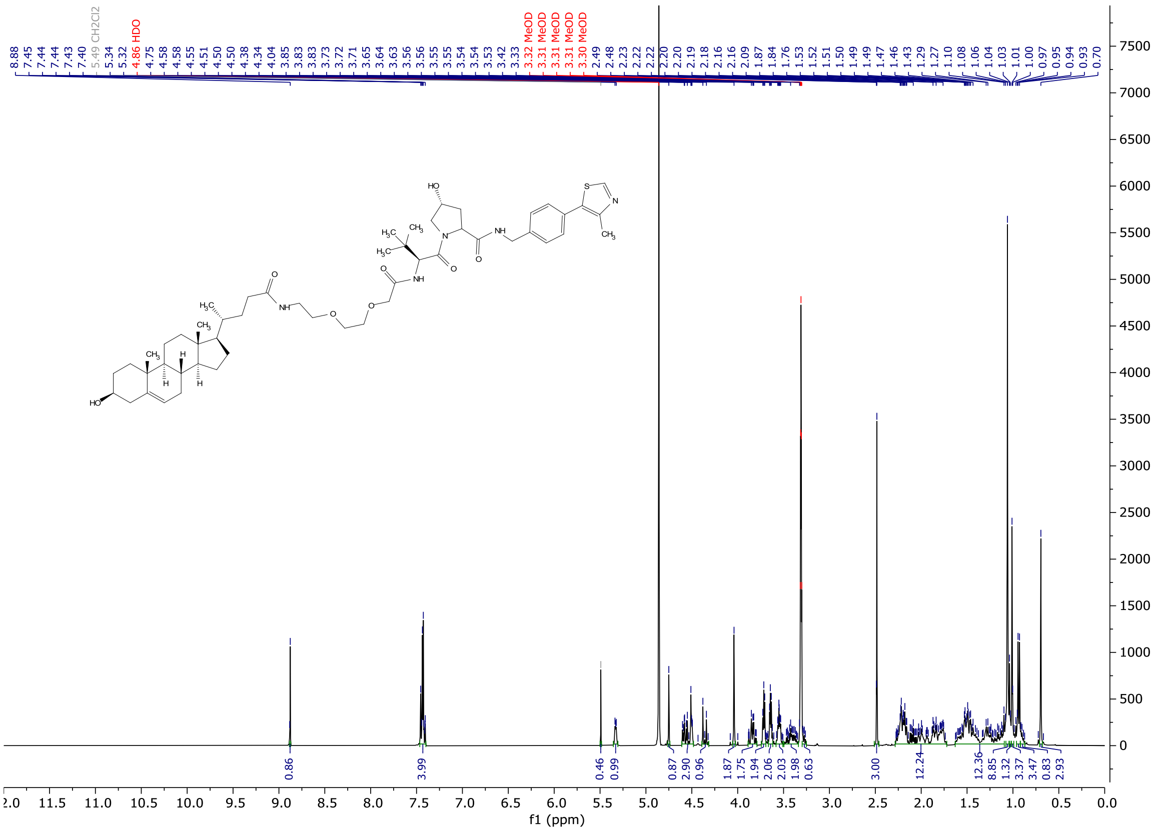

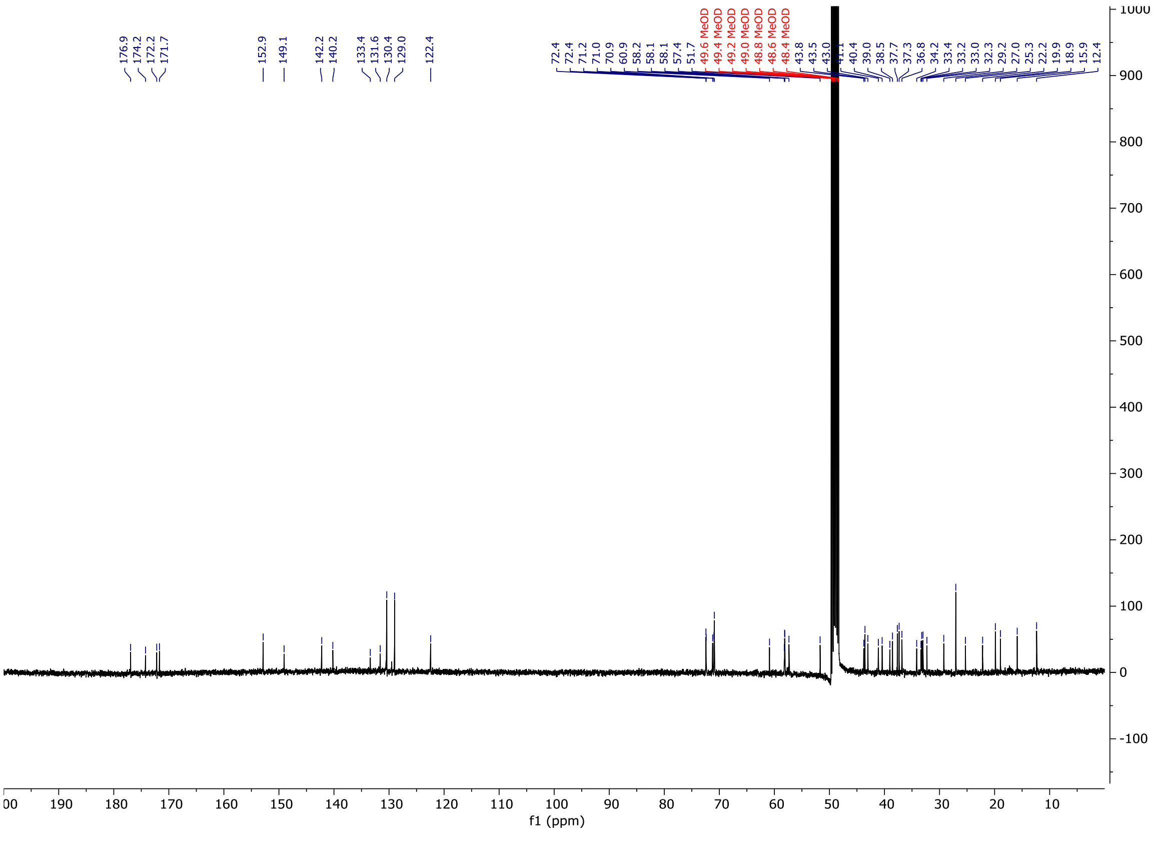

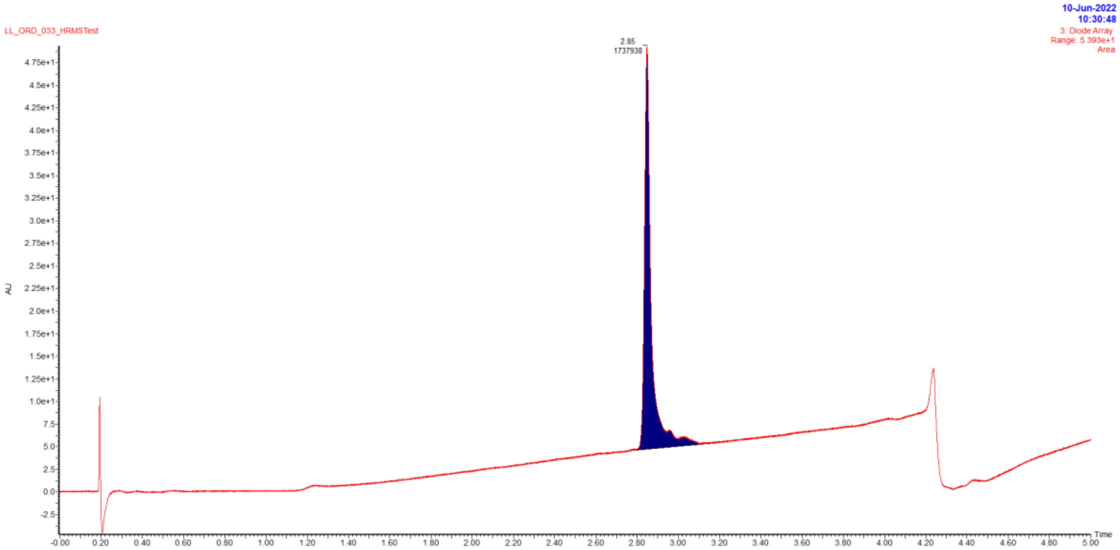


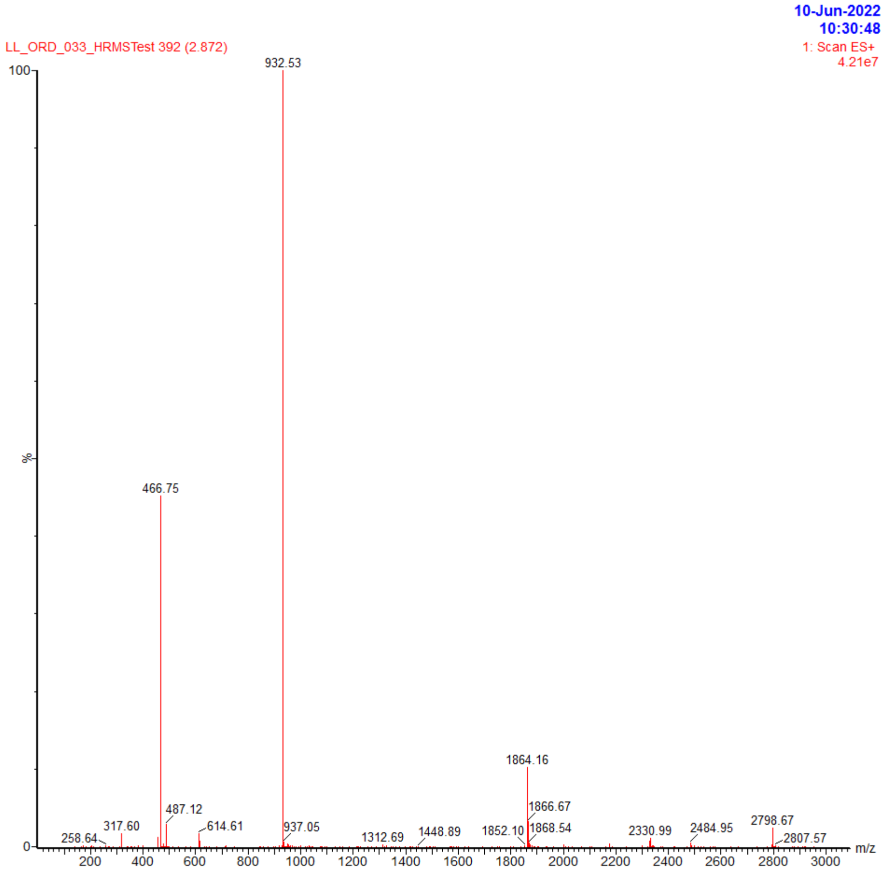


| Retention time | Area | Rel. Area(%) | Found (m/z) |
|---|---|---|---|
| 2.85 | 1737938 | 100 | 932.53 |
